# Semi-stability and unfolding of a KH domain modulate RNA binding and functions of MEX3A

**DOI:** 10.64898/2026.11.15.766395

**Authors:** Philipp H.O. Mayer, Charlotte Crauwels, Peijian Zou, Clara Hipp, Sam Asami, Donato Santovito, Christian Weber, Wim Vranken, Michael Sattler

## Abstract

MEX3A is essential for binding to Ago2/miR-126-5p and inhibition of apoptosis in endothelial cells implicated in cardiovascular health. Here, we discovered that the KH1 domain within the RNA binding region of MEX3A exists in equilibrium with an unfolded conformation. Using NMR spectroscopy and biophysical analysis, we dissect the KH1 folding kinetics and show that increased KH1 stability correlates with higher RNA affinity. The unfolded state of KH1 enables phosphorylation of T162, locking the domain in the unfolded conformation and enables recognition by 14-3-3 ζ. Residues in MEX3A KH1 that reduce its stability are located outside the RNA-binding interface and are evolutionary selected compared to human MEX3 paralogs. Semi-stability of KH domains in MEX3A and other multi-KH proteins may provide a general mechanism to tune RNA binding, enable phosphorylation and regulate biological function.

**Teaser:** MEX3A KH1 semi-stability modulates RNA binding and biological function.

## Introduction

Protein-RNA interactions regulate key functions in the cell, where multi-domain RNA binding proteins recognize cis-regulatory RNA sequences to modulate gene expression and RNA processing. The molecular processes responsible for this regulation are complex, involving multidomain RNA binding proteins with disordered regions, partial folding and post-translational modifications [1]. Many RNA binding proteins exhibit multiple KH domains, the second most abundant RNA binding fold [2, 3]. KH domains recognize 3-5 nucleotide RNA binding motifs [4, 5] involving the second β-strand and a conserved GxxG loop [6, 7]. Extensions of the KH domain as found in the STAR family proteins include an additional helix to recognize 6 nucleotide motifs [8]. The combination of multiple KH domains enables the recognition of clustered RNA motifs to enable specific binding and guide specific RNA processing functions [9–12].

Recently, the RNA binding protein MEX3A has been reported to play a key role for a non-canonical function of Ago2 depending on its interaction with miR-126-p. In endothelial cells suffering high sheer stress, the MEX3A tandem KH domains bind to a complex comprising Argonaute 2 (Ago2) and miR-126-5p (**Figure 1A**). The complex shuttles into the nucleus, where the miRNA suppresses apoptosis by binding and inhibiting Caspase-3. This suppresses apoptosis and contributes to the maintenance of cardiovascular health. The underlying molecular and structural mechanisms remain largely unresolved. [13, 14].

**Fig 1:**
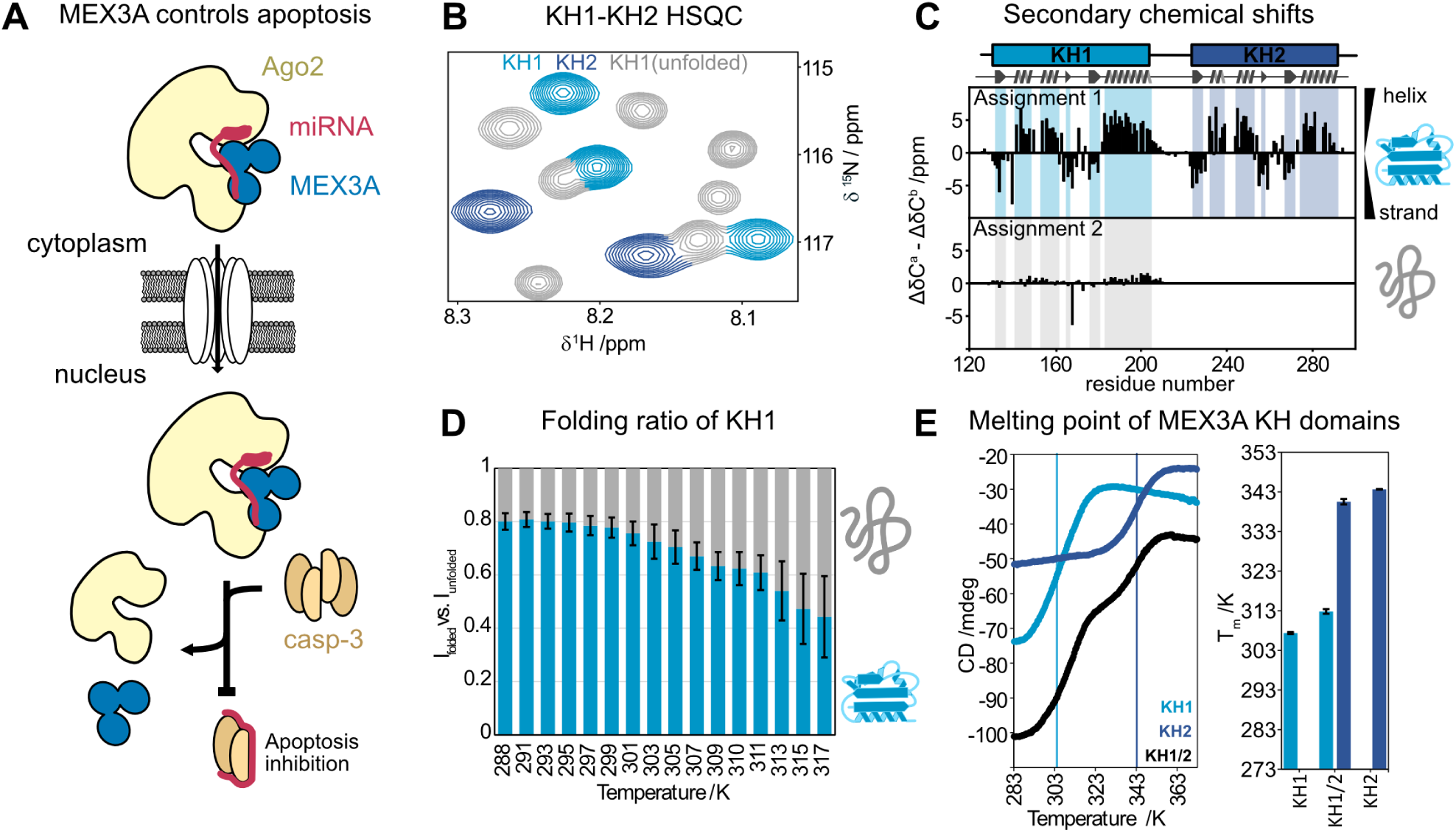
MEX3A KH1 exhibits an unfolded conformation in solution. **(A)** Schematic of the biological pathway in which MEX3A supports cardiovascular health. **(B)** ^1^H,^15^N HSQC spectra of KH1-KH2 show second set of NMR signals of an unfolded KH1 domain. **(C)** ^13^C secondary chemical shifts [(Δδ^13^C^α^) − (Δδ^13^C^β^)] of the second sets of signals (Assignment 2) indicate the presence of an unfolded conformation of KH1 with residual secondary structure. **(D)** The ratios of amide signal intensities of the folded and unfolded KH1 conformations decrease with increasing temperature between 288 K and 317 K and demonstrate that the unfolded conformation remains present. Peak height was measured, compared, and then averaged for 8 residues approximately equally spaced along the sequence. Error bars represent their standard deviation of the ratio between different residues. The increased standard deviations likely result from differential changes in solvent exchange at higher temperatures. **(E)** Thermal protein unfolding data observed in CD experiments show that KH1 and KH2 fold and unfold independently. At pH 5.5 KH1 unfolds around 312 K and 308 K in the tandem domain and isolation, respectively. KH2 folds at 340 K and 343 K in the tandem domain and in isolation, respectively. At pH 6.5, KH1 in isolation unfolds around 316 K (**Figure S4**).

The four human paralogs of MEX3A share a common domain architecture comprising two highly conserved RNA-binding KH domains flanked by long variable intrinsically disordered regions (IDR), and a C-terminal RING finger domain. The biological roles of the MEX3 homologs range from the regulation of stem cell properties [15] to functions in innate antiviral response [16].

Here, we investigated the RNA binding of the MEX3A tandem KH domains and discovered that the folded KH1 domain exists in equilibrium with a significant population of an unfolded conformation. We combined NMR spectroscopy, biochemical, and biophysical experiments to characterize the thermodynamic stability, folding kinetics and RNA binding of MEX3A KH1. We find that KH1 unfolding modulates RNA binding and enables phosphorylation of T162, which is not solvent accessible in the structure of the folded KH1 domain. Notably, KH1 phosphorylation locks it in an unfolded conformation, and the phosphorylated unfolded KH1 domain can be recognized by 14-3-3 ζ. Our data suggest that in a subset of KH domains in our proteome semi-stability is a feature to tune RNA binding and modulate interaction and function, providing a salient example of the complexity of RNA regulation by proteins, which goes beyond folded structures alone.

## Results

### MEX3A KH1 populates folded and unfolded conformations

The RNA binding of MEX3A is essential for its role in apoptosis regulation and cardiovascular protection. To study the molecular details of MEX3A RNA binding by NMR, we expressed the core RNA-binding region comprising its tandem KH domains. NMR ^1^H,^15^N HSQC correlation spectra of the MEX3A KH1-KH2 tandem domains exhibit more signals than expected (**Figure 1B; S1; S2**), suggesting that the protein adopts multiple conformations.

NMR backbone chemical shift assignments distinguish two conformations. The major state of the KH1 domain, as well as the KH2 domain correspond to a folded KH domain with the canonical βααββα secondary structure confirmed by NMR secondary chemical shifts (**Figure 1C**). In contrast, the second set of signals, which is in slow exchange with the folded form at the NMR chemical shift scale, corresponds to an alternative unstructured conformation, as indicated by random coil secondary chemical shifts, and high backbone flexibility seen in NMR relaxation data (**Figure 1C; S3**).

The exact population of this unfolded conformation is sensitive to temperature and pH (**Figure 1D; S4**), but all tested conditions showed an unfolded population of at least 20 % based on NMR signal intensities (**Figure 1D, S4**). Comparing the peak heights at different temperatures at pH 6.5 (**Figure 1D**) or pH 5.5 (**Figure S4**) shows unfolding of KH1 both by high temperatures and by low temperatures with the highest apparent stability at around 293 K. In line with this, thermal protein unfolding monitored by CD spectroscopy shows that KH1 has a significantly lower melting point than KH2 (313 K compared to 337 K), both in the tandem domain and in the independent isolated domains (**Figure 1E**). To rule out possible effects based on potentially truncated domain boundaries, we expressed protein constructs beyond the tandem KH domain. These included up to 129 residues of the unstructured N-terminal regions or up to an additional 20 residues C-terminally of the KH2 domain. We found that construct boundaries do not have a significant influence on the conformational equilibrium of the KH1 domain (**Figure S5**).

### Exchange kinetics between folded and unfolded conformations of MEX3A KH1

To quantify this persistent semi-stability, we first characterized the dynamic equilibrium between folded and unfolded conformation of the KH1 domain using ^15^N NMR exchange spectroscopy (EXSY) [17]. At room temperature, 298 K, EXSY shows fractions of 82.6 % (± 6 %) and 17.4 % (± 6 %) for the folded and unfolded state (**Figure 2A, S6**). These values are calculated from the folding and unfolding rates of 0.45 s^−1^ (± 0.22 s^−1^) and 0.09 s^−1^ (± 0.03 s^−1^), respectively. This implies a conformational stability of ΔG^0^ = 4.0 ± 1.5 kJ/mol, which qualifies the domain as ‘marginally stable’ according to Taverna & Goldstein [18].

**Fig 2:**
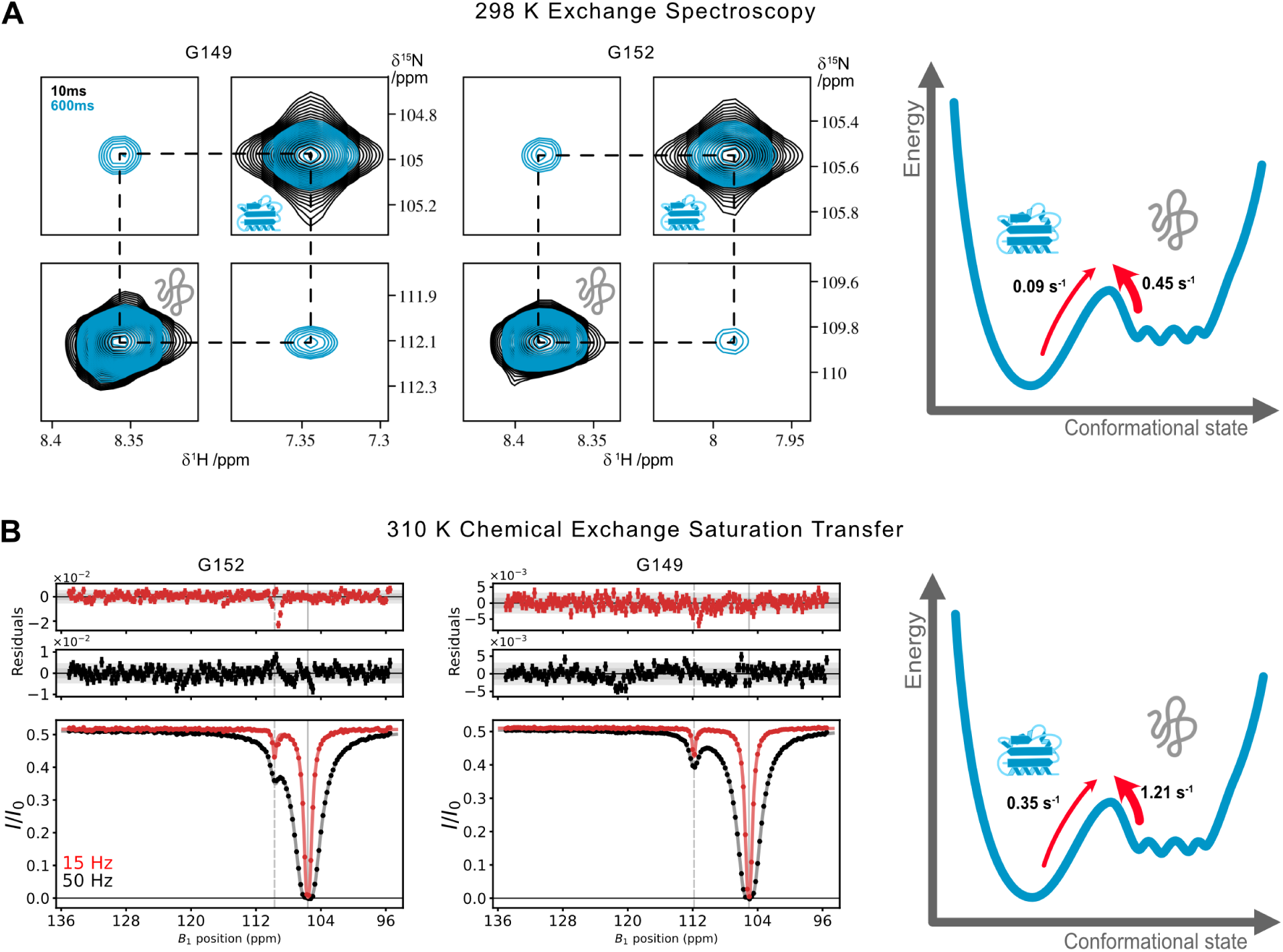
NMR exchange spectroscopy EXSY and CEST experiments provide un/folding kinetics of KH1 MEX3A KH1. **(A)** Zoomed view of auto- and cross-peaks of residues G152 and G149 in a ^15^N EXSY experiment recorded at 298 K. Spectra shown were recorded at pH 5.5, but quantification was based on spectra recorded at pH 6.5. Plots of the single residue fits used for quantification can be found in **Figure S7**. **(B)** ^15^N-CEST spectra of MEX3A KH1 recorded at 310 K at two *B*_1_ saturation fields with fits and residuals. Grey bars in the residuals plot indicate standard deviation and two times the standard deviation. The grey vertical line indicates the folded state resonance frequency. The dotted vertical line indicates the resonance frequency of the unfolded state. Plots for all residues can be found in the supplement (**Data S1, S2**).

We further characterized the conformational dynamics at 310 K [19], close to physiological temperatures. To minimize reliance on signals of the unfolded state, which are affected by proton exchange with the solvent, we used chemical exchange saturation transfer (CEST) spectroscopy analyzing the traces of the folded state peaks (**Figure 2B, Data S1, S2**). CEST traces of folded state peaks only show two dips in the signal along the ^15^N dimension, which indicate that a two-state folding model is appropriate. The fit for a two-state model converges with a reduced χ2 of 2.0. The CEST experiment shows that the unfolded state represents 22.5 % (± 0.5 % based on 100 Monte Carlo simulations), with an exchange rate of 1.56 s^−1^ (± 0.04 s^−1^). Based on these data, the unfolding and folding rates are 0.35 s^−1^(± 0.01 s^−1^) and 1.21 s^−1^(± 0.01 s^−1^), respectively (**Figure S8**). The conformational stability ΔG^0^ = 3.2 kJ/mol is in reasonable agreement with the stability measured at 298 K using EXSY experiments.

The persistence of the unfolded population, as well as the melting point close to physiological temperatures, suggests that the unfolded state might have a specific role for the biological function of MEX3A.

### Semi-stability modulates protein-RNA interaction

Since the unfolded state does not bind RNA (**Figure S9, S12**), the observed semi-stability would limit the binding-competent concentration of MEX3A KH1 in cells. But does the stability of the domain also affect the binding of RNA to the folded state of the protein? To answer this question, we created a strongly stabilized mutant (T140S, T188A) and a destabilized mutant of the KH domain (D185A) based on predictions by Protposer [20]. We then monitored RNA binding to a region of miR-126-5p (position 15-19; 5’ GUACG) that we have identified to represent a high affinity motif. Frequency shifts and line-broadening in NMR titrations of the RNA to ^15^N-labeled KH1 were analyzed based on the Bloch-McConnell equations with TITAN [21], employing a two-state binding equilibrium that incorporates a conformational selection model to account for the unfolded state not binding. The folding and unfolding rates required to fit the conformational selection model were extracted from EXSY experiments (**Figure 3A**).

**Fig 3:**
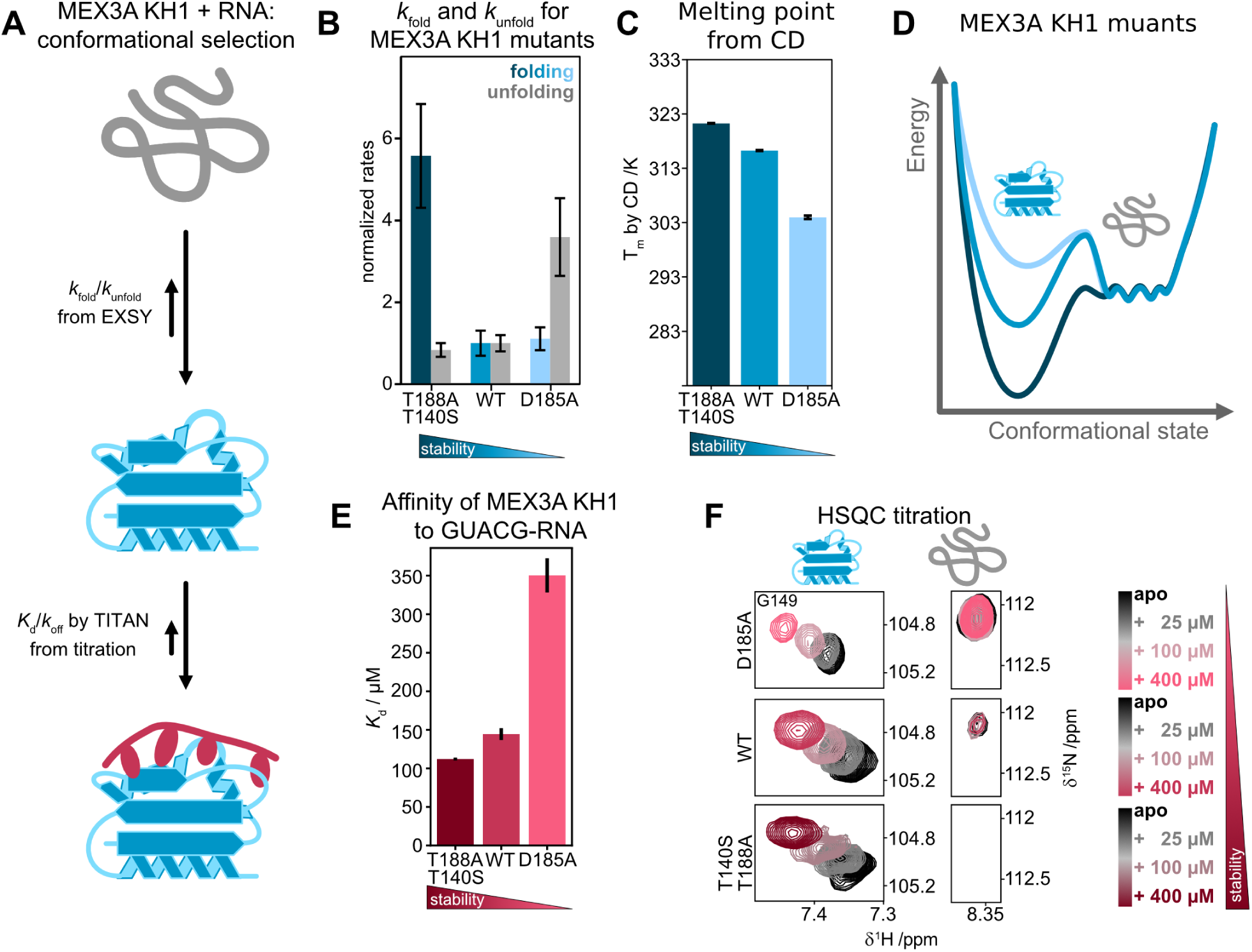
Mutations affecting the semi-stability modulate MEX3A KH1 affinity to RNA. **(A)** Schematic of the probed populations: unfolded state, folded state and RNA-bound state of the domain. Folding dynamics calculated from EXSY experiments, then used as input for the determination of *K*_d_ and *k*_off_ by TITAN. **(B)** Relative folding rate (blue) and unfolding rate (grey) of MEX3A KH1 WT, stabilized mutant T188A/T140S and the destabilized mutant D185A normalized to the rates of the WT protein, measured at 298 K. Errors were derived from the standard deviation of the single residue fits and error propagation after normalization. **(C)** Melting point of KH1 domain variants determined by CD. 321.3 K/316.3 K/304.0 K for T140S T188A/WT/D185A mutants. **(D)** Schematic energy diagram for the wildtype, stabilized and destabilized KH1 domains. **(E)** *K*_d_ for RNA (GUACG) binding of KH1 domain variants, based on NMR titration analyzed with NMR TITAN. **(F)** Zoomed view of the NMR titrations analyzed in (E) showing the folded and unfolded state of G149 for all three versions of KH1.

CD melting experiments, as well as EXSY-derived folding and unfolding rate demonstrate the strong de/stabilization of the mutants (**Figure 3B, C, D**). Compared to the wildtype protein (*K*_d_ = 145 µM) the affinity is increased for the stabilized mutant (*K*_d_ = 112 µM), while it iss decreased for the destabilized mutant (*K*_d_ = 356 µM) (**Figure 3E, F, S9**). Interestingly, stabilization and destabilization of KH1 are correlated with increased unfolding rates (*k*_unfold)_ and folding rates (*k*_fold_), respectively (**Figure 3B,C**).

### Semi-stability allows phosphorylation at otherwise inaccessible sites

The modulation of RNA binding by the folding kinetics is an interesting feature of fine-tuning RNA binding affinity. However, we wondered if there are other roles for the KH1 semi-stability; whether the unfolded state of MEX3A KH1 itself has a functional role. In fact, the presence of an unfolded conformation has been reported for the KH1 domain of FUBP2, where phosphorylation locks the KH domain in an unfolded conformation [22]. We thus explored if MEX3A KH1 might employ a similar mechanism (**Figure 4A**).

**Fig 4:**
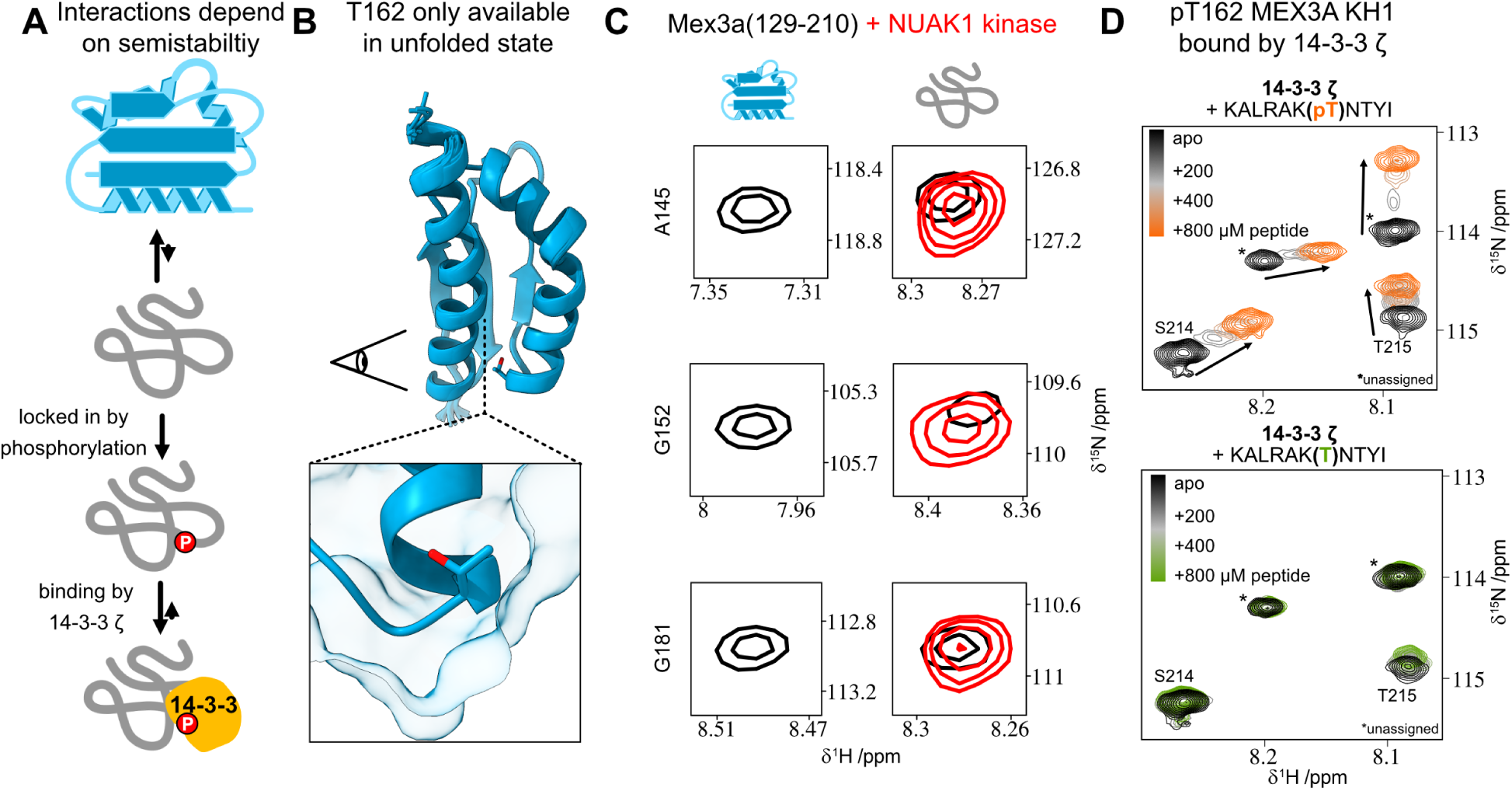
The unfolded state of MEX3A enables phosphorylation and subsequent 14-3-3 binding. **(A)** Schematic of a possible function of MEX3A KH1s unfolded population with phosphorylation and 14-3-3 binding being enabled. **(B)** NMR structural ensemble of the stabilized MEX3A KH1 domain. The hydroxyl-group of the phosphorylation site T162 is highlighted as being buried. **(C)** Zoomed views of ^1^H,^15^N-HSQC spectra of MEX3A KH1 signals representing the folded and unfolded state, before (black) and after (red) phosphorylation by NUAK1. **(D)** NMR titration shows 14-3-3ζ binding to a peptide representing pT162 at a *K*_d_ 144 µM, while the unphosphorylated peptide is not bound.

Prediction algorithms for binding sites of 14-3-3 proteins, which are well characterized phospho-peptide binding proteins, show a highly likely interaction site within the KH1 domain of MEX3A centered on pT162 [23] (**Figure S10**). T162 phosphorylation has been reported in a phosphoproteomics study in PC3 colorectal cancer cells [24], a tumor which shows upregulation of MEX3A [25], but as the sequence around T162 is conserved between the four human MEX3 paralogs, the eukaryotic phosphorylation site database (EPSD) [26] annotates the corresponding peptide as phosphorylated for all four MEX3 paralogs.

To evaluate this potential phosphorylation site, we determined the NMR structure of an extended MEX3A KH1 domain (**Table S1**), revealing that the T162 side chain is not solvent exposed and thus unavailable for post-translational modification in the folded state (**Figure 4B**). The T162 side chain, located at the end of helix α2, is turned towards the interior of the protein and engages hydrophobic contacts involving the side chain methyl group, confirmed by several NOEs (**Figure S11, Table S2**). The side chain of T162 is indeed essential for maintaining the fold of MEX3A KH1: When the residue is mutated to alanine or the phosphomimic T162E, the structure is entirely abolished (**Figure S13**). Accordingly, a phosphorylation of this residue could only occur in the unfolded state of the protein. We predicted potential kinase candidates for T162 phosphorylation using the kinase library prediction tool [27] of PhosphositePlus [28], which identified the kinase NUAK family SNF1-like kinase 1 (NUAK1) as a likely candidate.

NUAK1 does indeed phosphorylate MEX3A KH1 in vitro, which locks the domain into the unfolded conformation as judged by the NMR ^1^H,^15^N HSQC spectra, which show exclusively signals of the unfolded state after phosphorylation (**Figure 6C**). Based on minimal chemical shift changes, the C-terminal residue S210 also appeared to be phosphorylated off-target (**Figure S13E**). However, the conformational effect of pS210 is likely minimal, as phosphomimetic S210E shows only minimal changes on the melting point and folding state of KH1 (**Figure S13G**), while the phosphomimetic T162E is also locked in the unfolded state (**Figure S11**).

Next, we assessed whether the unfolded phosphorylated KH1 domain can bind to 14-3-3. To this end, deuterated 14-3-3 ζ was titrated with a phosphorylated peptide representing MEX3A. This peptide does indeed bind to 14-3-3 ζ with a *K*_d_ of 142 µM, as measured by NMR titration (**Figure 6D**). This is on the same order of magnitude as other affinities reported for the interaction of 14-3-3 ζ proteins with phosphorylated peptide [29]. An unphosphorylated peptide of the same sequence did not bind to 14-3-3 ζ (**Figure 6D**).

These results suggest a novel mechanism of how RNA affinity is fine-tuned by semi-stability and shows how the unfolded state of MEX3A KH1 is required for the phosphorylation of T162. The biological effects of this phosphorylation, including the role of 14-3-3 interactions in cells remain to be investigated.

### MEX3A is unique among its paralogs by both decreased folding and decreased stability

We next asked which sequence features might confer this and whether these features are evolutionary conserved. To answer this, we studied the KH1 domains of MEX3A’s human paralogs. The four investigated KH1 domains differ significantly in thermal stability, as indicated by CD-monitored melting points (**Figure 5A**). MEX3A KH1 has the lowest melting point (316.2 K), followed by MEX3B at 8 K higher (324.6 K), MEX3D (329.5 K), and finally MEX3C (331.1 K). This pattern is also apparent in the NMR spectra at 298 K (~25°C), where the unfolded conformation of MEX3A KH1 is clearly visible, while the other three proteins do show unfolded conformations, but in significantly reduced populations (**Figure 5B**).

**Fig 5:**
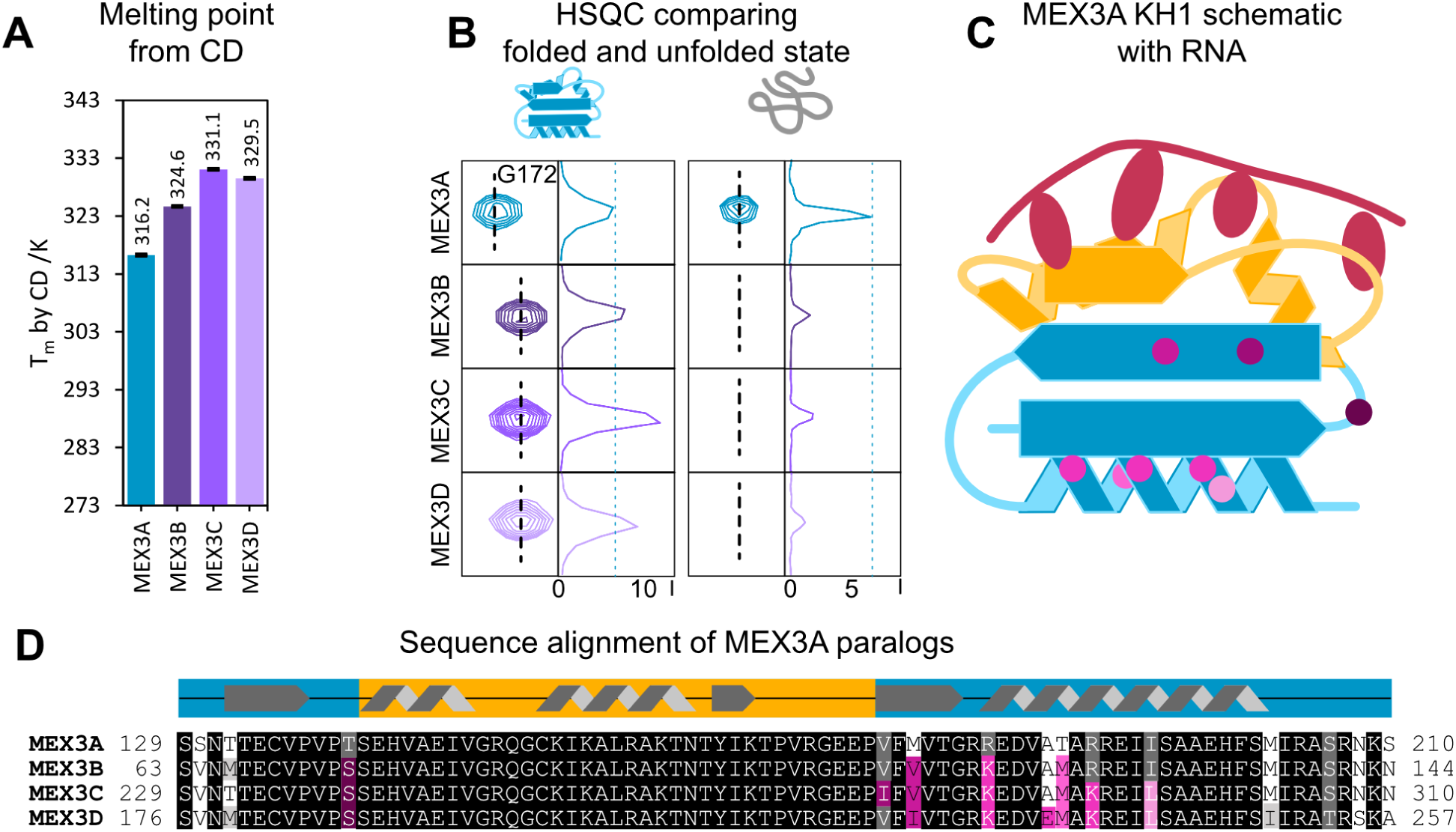
Semi-stability is a feature of MEX3A not conserved in its paralogs. **(A)** Melting point of KH1 of MEX3A paralogs with MEX3A being the least stable. **(B)** Amide NMR signals and 1D traces for G172 in the folded and unfolded state, and the corresponding residue in MEX3B/C/D. **(C)** Schematic of a KH domain (blue) showing the RNA-binding interface (yellow) and locations of sequence variations between the human MEX3A-D KH1 domains (shades of pink). **(D)** Multiple sequence alignment (MSA) of KH1 of human MEX3A paralogs. Color coding as in (C).

MEX3A’s human paralogs reveal that the primary sequence of the RNA binding interface is strictly conserved, suggesting that their RNA-binding specificity is conserved (**Figure 5C, D**, orange). However, there are sequence variations in the KH domain flanking the residues that form the core RNA-binding interface (**Figure 5C, D**, blue), as well as the distal, C-terminal helix.

To identify the sequence features that are responsible for the drastic differences in stability between the paralogs, we created point mutants of MEX3A KH1 that capture sequence variations from the other domains. For example, MEX3A contains alanine at position 187, whereas MEX3D contains glutamate at the corresponding position. We therefore introduced the A187E substitution into MEX3A KH1 (**Figure 6**). A187 is located on the solvent exposed side of the C-terminal helix α3. Helix projections suggest an increased stability of the mutant: The sidechain’s negative charge may serve to balance the charges of two arginine sidechains (R183 and R190) (**Figure 6A**).

**Fig 6:**
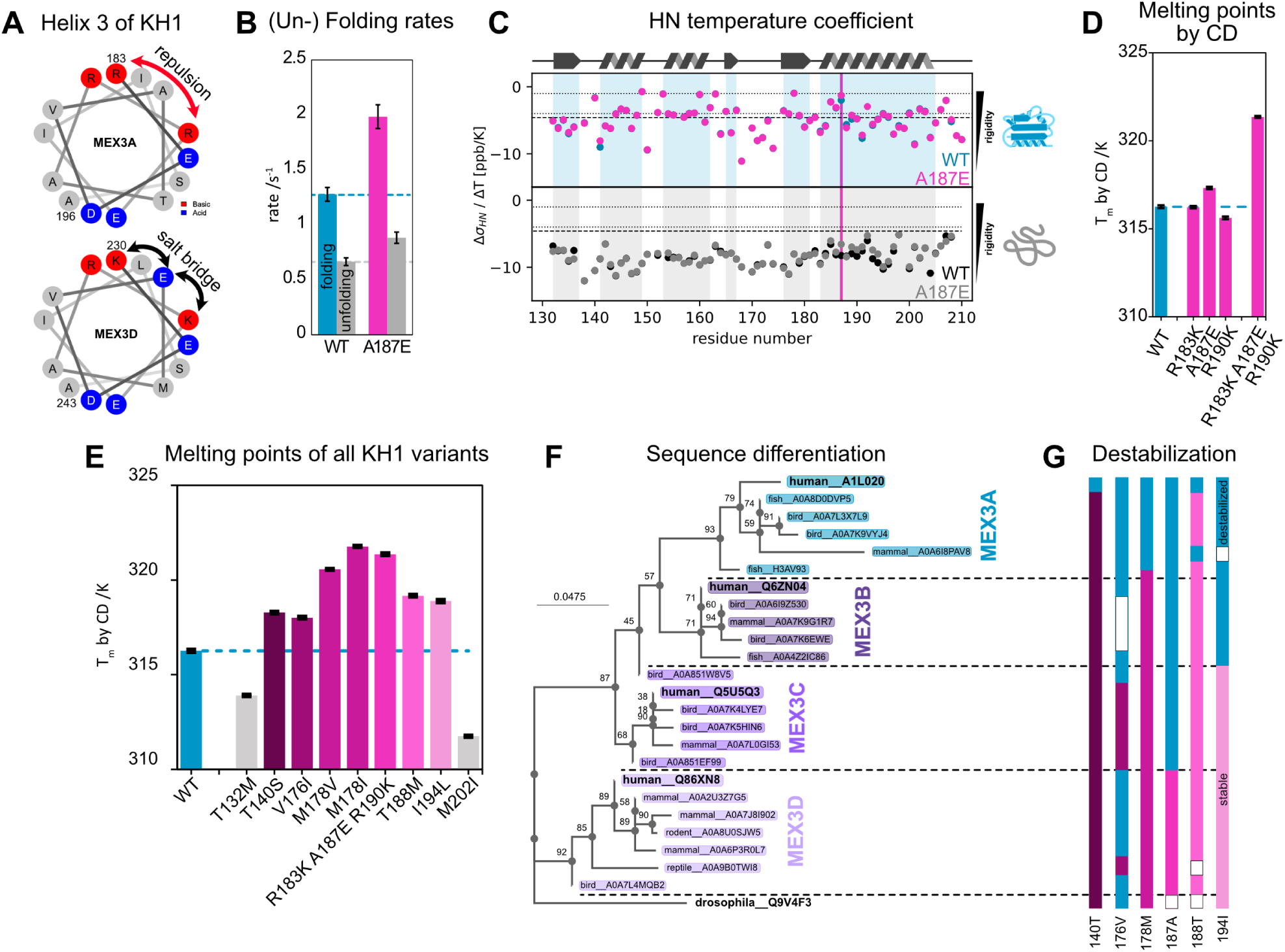
Sequence variations of MEX3 paralogs combine to specifically destabilize MEX3A KH1. **(A)** Helix projection of α3 of KH1 in MEX3A and MEX3D. Evenly distributed charges might stabilize MEX3D. **(B)** ZZ-exchange confirms higher folding rate in the MEX3D mimicking mutant A187E. **(C)** Amide proton temperature coefficients for the folded and unfolded states of MEX3A KH1 and the A187E mutant. Dashed lines indicate the range of likely hydrogen bonding [30, 31]. **(D)** CD-derived melting points of MEX3A KH1 and select mutants. R183K and R190K synergize with A187E. **(E)** Melting points of MEX3A KH1 with point mutations mimicking the more stable human paralogs. Core domain mutations in pink, peripheral mutations in grey. Subsets of combinations from (D) omitted for clarity. **(F)** Maximum-likelihood phylogenetic tree of MEX3 KH1 sequences. Node values indicate bootstrap support percentages. **(G)** Sequence variations of the phylogenetic tree in (F): Blue indicates the amino acid in MEX3A, shades of pink indicate comparatively stabilizing amino acids and white indicates amino acids with unverified effects. Destabilizing sequence variations are by far the most prevalent in MEX3A.

^15^N-EXSY experiments indeed show an increase in the folded state population of A187E compared to WT (**Figure 6B, S15**). Specifically, A187E substantially increases the folding rate of KH1, while the unfolding rate is only slightly increased. To assess the effects of the mutation on stability parameters at single residue resolution, we compared temperature coefficients for the backbone amide protons. These report on hydrogen bonding and solvent accessibility. Between A187E and WT, the unfolded state’s temperature coefficients are more significantly perturbed than those of the folded state (**Figure 6C**). These perturbations are limited to the residues surrounding A187 in the helix α3, suggesting that the effect of the mutation is primarily local, rather than global. The increases suggest that rigidified, hydrogen bonded conformations are more frequently sampled, even though the coefficients are still somewhat outside the range normally expected for full hydrogen bonding [30, 31]. Mutation A187E could thus increase the folding rate of the KH1 domain by pre-forming part of helix 3 in the unfolded state.

R183K and R190K are sequence variations that change the amino acid property less drastically, retaining the positive charge and long sidechain. These mutations indeed do not increase the stability of the MEX3A KH1 domain. However, in the combination with A187E, they significantly stabilize MEX3A KH1 (**Figure 6D**): The A187E mutation synergizes with R183K and R190K to form stronger electrostatic interactions and stabilize the C-terminal helix [32].

While mutations in the terminal regions of the domain are likely affected by boundary effects, every variation in the core domain of MEX3A KH1 decreases the thermal stability (**Figure 6E**). Overall, the effects and locations of the amino acid changes in the other three KH domains suggest that they drive an increase in stability compared to MEX3A KH1, while maintaining the RNA binding specificity across the MEX3 paralogs. Most destabilizing substitutions that distinguish MEX3A KH1 from the other MEX3 KH1 paralogs are shared among MEX3A homologs, suggesting that semi-stability may be conserved within this lineage (**Figure 6F**). The unfolded KH1 population at physiological temperatures could therefore reflect a functional adaptation specific to MEX3A.

### Semi-stability also occurs in other multi-KH domain proteins

Of the four MEX3 paralogs only MEX3A shows a significant fraction of unfolded conformation, while the KH1 domains of MEX3B/C/D have unfolded populations smaller than 5% (**Fig. 5B**). As mentioned, functional semi-stability has been previously reported for FUBP2 KH1 [22], with a melting point of 325 K, i.e. comparable to MEX3A KH1, but significantly lower than the melting point of other KH domains in the same protein (342.3 K, 335.2 K, and 334.2 K for FUBP2 KH2, KH3, and KH4 respectively) [11, 22]. We next wondered, whether other proteins also contain semi-stable KH domains.

Of all KH domains in the human proteome, (84 domains across 30 proteins according to InterPro, Pfam PF00013 [33]) the protein HDLBP alone contains 14 KH domains. Structures of isoalted HDLBP KH domains have been reported (PDB 2CTE, 2CTF, 1VIH, 2CTJ, 2CTK, 2CTL, 2CTM) [34, 35]. It is noteworthy that for KH7 and KH9 of HDLBP no structures have been reported, while experimental structures are available for KH6, KH8 and KH10. We hypothesized that this could indicate semi-stability of KH7 and KH9.

Indeed, when expressed and characterized, KH domains 7 and 9 presented unusually poor ^1^H,^15^N-HSQC spectra (**Figure S17**), with an unexpectedly high number of peaks, some with poor dispersion, and heterogeneous peak intensities. The same two KH domains also have drastically lower melting points compared to the surrounding KH domains (**Figure 7C**). Analogously to MEX3A KH1, characteristic profiles in EXSY spectra confirm that both domains have two interchanging populations in solution that exchange on a timescale of hundreds of milliseconds to a few seconds (**Figure 7A**). By compiling the melting curve midpoints from our work and the published literature, it is apparent that KH domains can cover the entire spectrum of structural stability (**Figure 7B**).

**Fig 7:**
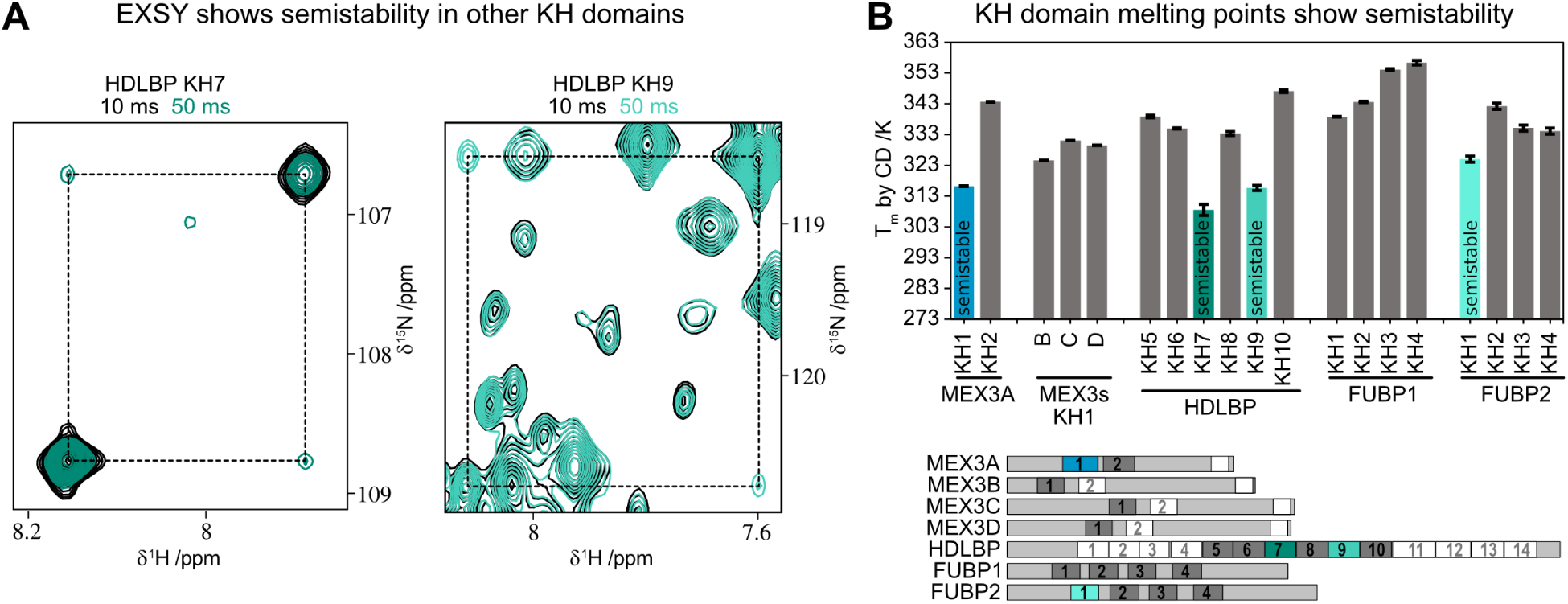
Semi-stability occurs in several multi-KH domain proteins. **(A)** EXSY experiments on HDLBP KH7 and KH9 reveal their semi-stability. **(B)** Melting temperatures of various KH domains reveal semi-stability to be a feature of several unrelated KH domains. Domains judged to be stable in grey, semi-stable domains in color. Untested KH domains in the schematic in white.

To assess whether existing bioinformatics tools could identify additional semi-stable KH domains, we compared the four experimentally confirmed semi-stable domains with a control set of stably folded KH domains for which apo structures are available. Different sequence and structure-based scores of disorder were unable to clearly distinguish between the two groups (**Figure S19**), suggesting that these measures do not reliably identify semi-stability in the domains examined.

We next assessed whether computational stability predictors could capture the effects of the stabilizing substitutions identified through comparison of MEX3A KH1 with its paralogs. Both an AI-driven structure-informed method and a physics-based method predicted relatively small effects for these substitutions compared with alternative mutations at the same positions (**Figure S20**).

These results indicate that the computational methods tested cannot fully predict the experimentally observed intricate folding behavior of these KH domains, nor consistently predict the effects of stabilizing substitutions in MEX3A KH1. Experimental measurements of folding kinetics and stability are required for developing computational approaches, especially in complex cases where multiple populations are present, such as here for the MEX3A KH1 semi-stability.

## Discussion

Only few proteins exhibit a significant population of an unfolded conformation with sufficiently slow folding kinetics to allow detailed investigated [36–38]. Low structural stability can have drastic implications for the functions of proteins, including unique modes of interaction, varied subcellular localization, or loss of function compared to a more stable version of the protein [22, 39]. In this study, we investigated the molecular effects of semi-stability of a KH domain and explored its potential role for the regulation of biological function.

For MEX3A, across mutants a lower unfolding rate correlates a higher affinity to RNA, suggesting that semi-stability is a way to fine tune RNA-affinity of a KH domain. Additionally, certain residues may be accessible for phosphorylation only in the unfolded state of the protein. T162 of MEX3A is one such residue. This phosphorylation of MEX3A KH1 locks the domain in an unfolded state, similar to results published on FUBP2 KH1 [22]. The effect of this phosphorylation goes beyond abolishing RNA binding: 14-3-3 proteins are predicted to bind MEX3A pT162 and 14-3-3 ζ does indeed bind the corresponding phosphorylated peptide (**Figure 8A**)

**Fig 8:**
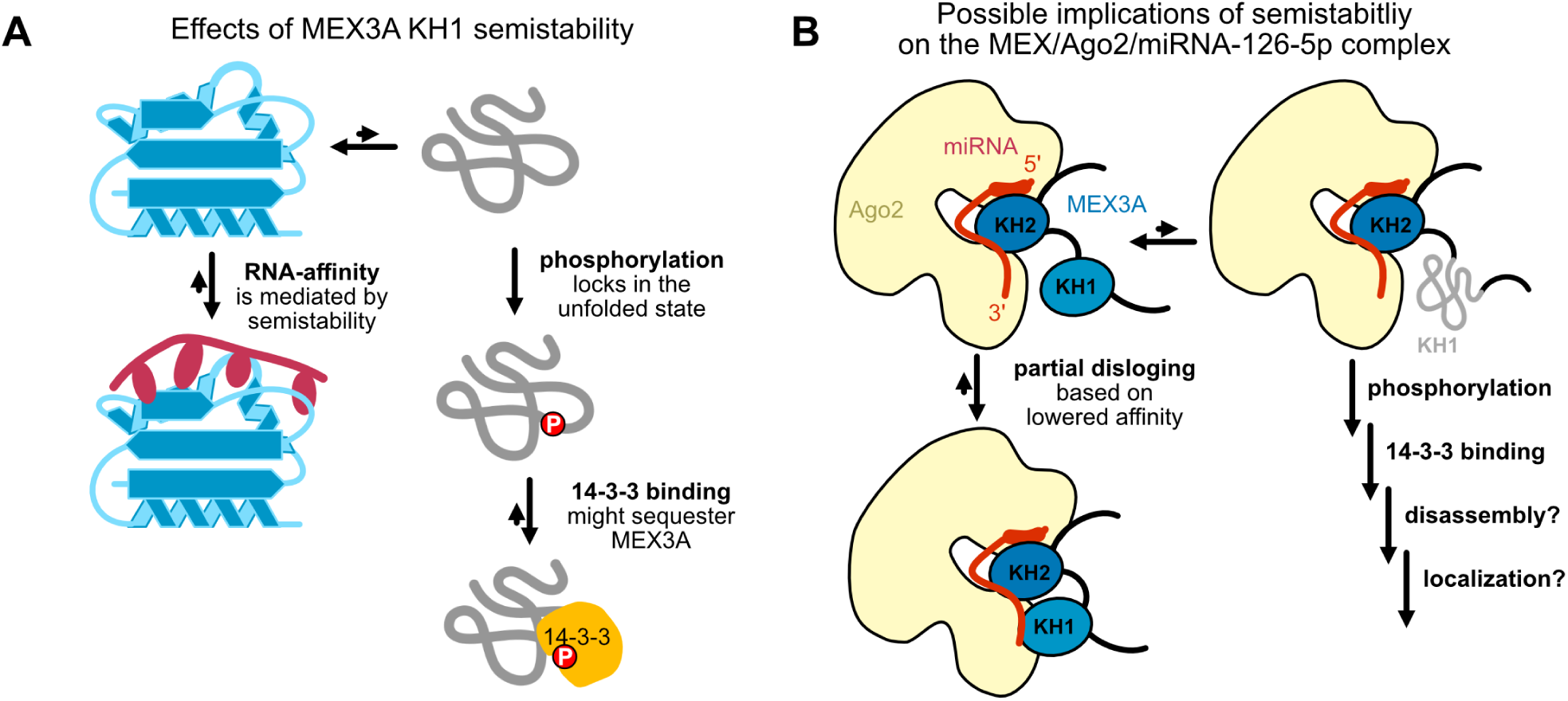
Summary and potential role of KH1 for MEX3A function. **(A)** Semi-stability’s effect on MEX3A‘s function, as elucidated in this study. **(B)** Semi-stability’s proposed influence on the Ago/Mex/miR126-5p complex.

In the context of the complex of MEX3A with Ago2 and miRNA-126-5p, the unfolded fraction of MEX3A KH1 could enable phosphorylation of T162. This would prevent refolding and shift the population of KH1 to a completely unfolded state, which in turn would enable binding of 14-3-3. The unfolded fraction could also contribute to Ago2 binding and stabilize the Ago2/miRNA complex, e.g. to allow for nuclear translocation. Binding of the phosphorylated KH1 domain by 14-3-3 might aid to dissociate MEX3A from the Ago2/miRNA complex (**Figure 8B**). It has been recently suggested that 14-3-3 binding can promote the disassembly of complexes [41]. 14-3-3 proteins could play a role in enabling the anti-apoptotic function of MEX3A by segregating the protein in the nucleus after the disassembly of the MEX3A/miRNA-126-5p/Ago2 complex. Alternatively, phosphorylation might inhibit MEX3A function by sequestering MEX3A in the cytoplasm, so that dephosphorylation would be required for nuclear shuttling.

To understand the causes of MEX3A KH1 semi-stability, we identify unique sequence features that are linked to its semi-stability and differentiate MEX3A from its human paralogs: While the four KH1 domains have identical RNA-binding interfaces, variations in other regions contribute to the semi-stability in MEX3A. Importantly, these sequence variations are evolutionary conserved, consistent with an important functional role for the semi-stability of MEX3A KH1.

Finally, we have identified additional KH domains which exhibit a semi-stability in HDLBP, a multi-KH domain protein. KH domains with high melting curve midpoints are common: MEX3A KH2, HDLBP KH10, as well as FUBP1 KH2-4 even surpass some published thresholds to being considered thermostable [42, 43]. Thus, our data suggest that semi-stability of KH domains may be an evolutionary selected feature that has been acquired to enable additional layers of regulating protein-RNA interactions and molecular functions of multidomain RNA binding proteins.

## Materials and Methods

### Protein expression and purification

All MEX3A constructs (KH1: 129-210 and KH1-2: 125-294) were expressed fused to N-terminal His-tags and SUMO-tags constructs in a pET13 vector in BL21(DE) cells. Addition of 0.5mM IPTG induced the expression that proceeded overnight for about 16 hours at 18-20 °C.

The cells were lysed in 50 mM HEPES, pH 7.5, 500 mM NaCl, 1-3 mM β-mercapto-ethanol, 10 mM imidazole supplemented with commercial protease inhibitor. The cells were lysed using sonification, and the cleared lysate was subjected to IMAC.

The protein was eluted in 50 mM HEPES, pH 7.5, 500 mM NaCl, 1-3 mM β-mercapto-ethanol, 300 mM imidazole. The eluate was supplemented with 0.2 mg SenP2 SUMO-hydrolase and dialyzed overnight against 20 mM HEPES, pH 7.5, 500 mM NaCl, 1-3 mM β-mercapto-ethanol, and 10 mM imidazole. After dialysis, the protein was subjected to reverse IMAC.

In cases where the protein would be used in RNA-binding experiments or crystallization, it was dialyzed or diluted until the buffer contained no more than 50 mM NaCl and then further purified using ion-exchange chromatography using a HiTrap SP column. Buffer A and B contained 10 mM HEPES, pH 7.5 and 0 mM NaCl and 1M NaCl, respectively. Finally, proteins were separated using a HiLoad™ 16/600 Superdex™ 75 pg size-exclusion column.

Constructs of FUBP1 were purified as described previously [9].

All other constructs were purified from a pET11 vector with a TEV-cleavable His6-tag. This includes HDLBP KH5 (430-506), KH6 (504-580), KH7 (576-652), KH8 (649-726), KH9 (722-799), KH10 (797-873) and MEX3B (63-144), MEX3C (229-310), MEX3D (177-257), as well as 14-3-3 ζ (1-233).

### NMR Spectroscopy

NMR experiments were recorded at 900 MHz, 800 MHz or 600 MHz using Bruker Avance III consoles and cryo-TCI probes. Proteins were measured in 20 mM NaP, pH 6.5, 150 mM NaCl, 10 mM DTT, and 5 % D2O.

#### Processing and analysis

The spectra were recorded with TopSpin versions 3.6, 3.7, and 4.0. All collected NMR data were processed with NMRPipe [44]. CCPNMR v3 [45] was used for qualitative analyses, backbone assignments, peak height extraction, and image generation.

Triple resonance experiments were recorded with ~10 % NUS in the case HNCO and HNcaCO, and 25 % in the case of HNCACB and CBCAcoNH. Topspin’s standard implementation of NUS was used [46–48]. The spectra were reconstructed using the SMILE or IST algorithms implemented in NMRPipe [49, 50].

#### Resonance assignment

Generally, resonance assignments were achieved using standard triple resonance experiments: HNCACB, CBCAcoNH, HNCO, HNcaCO, as well as a ^15^N-edited NOESY experiment.

For the MEX3A(129-210) construct representing KH1, only the wild type was assigned with triple resonance and EXSY experiments at 298 K. Transfer to other temperatures was achieved with a series of ^1^H,^15^N-HSQC experiments, spaced by no more than 3 °C. The transferred assignments were corrected and validated using a ^15^N-edited NOESY experiment, as well as an EXSY experiment, which correlates the peaks of the folded and the unfolded state.

The assignment of MEX3A(129-210) mutants was achieved by transferring the assignments of the wild type and then correcting and validating strongly perturbed peaks using EXSY and, if necessary, a ^15^N-edited NOESY experiment.

The assignments of MEX3A(108-210) T140S T188A, which is the basis for the structural model, were created by NMRtist without human intervention.

The reference values to calculate secondary chemical shifts of C^α^ and C^β^ atoms were taken from Wishart et al. [51].

#### Peak height comparison over temperature

To qualitatively compare the proportions of folded and unfolded state over a temperature range, peaks heights of residues spread across the sequence of the KH1 domain. Residues 132, 142, 152, 162, 172, 184, 192 and 202 were compared for MEX3A(129-210) at pH 6.5 (**Figure 1D**) and for MEX3A(125-292) at pH 5.5 (**Figure S4B**). Averages of the ratios for single residues are displayed with standard deviation as error bars.

#### HSQC titrations

Titration of miRNA-126-5p(15-19) to MEX3A constructs was conducted on a 600 MHz spectrometer at 298 K. CSPs were calculated using:

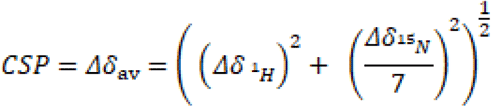

Titrations of MEX3A KH1 mutants with RNA were analyzed using NMR TITAN [21], using a conformational selection model:

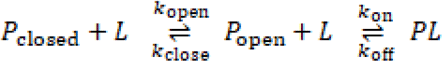

Which, in this specific case, is:

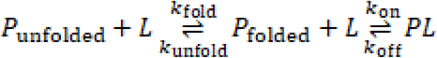

With these parameters given: P as protein concentration, L as ligand concentration, *k*_close/fold_ as the folding rate and *k_open_*_/unfold_ as the unfolding rate. With the dissociation constant (*K*_d_) and the dissociation rate (*k*_off_) derived from the fit.

The parameters for the two folded and unfolded state in the absence of ligand were fitted in a first step based purely on the apo spectrum. The folding and unfolding rates were set based on the results of EXSY experiments measured at 298 K on a 600 MHz spectrometer. The parameters for the bound state, as well as the *K*_d_ and the *k*_off_ were fitted in a second step using all spectra of the titration. Errors were calculated using 100 iterations of bootstrap error analysis. Of the 50µM MEX3A(129-210) WT, 20 residues were tracked and used for fitting over nine 9 titration points ranging from 0 to 1000 µM RNA. Of the 50 µM MEX3A(129-210) D185A mutant, 23 residues were tracked and used for fitting over 9 titration points ranging from 0 to 800 µM RNA. Of the 50 µM MEX3A(129-210) T140S T188A mutant, 18 residues were tracked and used for fitting over 9 titration points ranging from 0 to 400 µM RNA.

The titration of MEX3A derived peptides to 14-3-3 ζ was conducted on 200 µM perdeuterated 14-3-3 protein at 298 K on a 900 MHz spectrometer using ^15^N TROSY experiments. The phosphorylated and unphosphorylated peptides were acquired commercially from Peptide Specialty Laboratories GmbH (Heidelberg, Germany). The synthesized sequences are KALRAK[pT]NTYI and KALRAKTNTYI. They were resuspended to a stock concentration of 10 mM and their pH was adjusted to ~6.5. The phosphorylated peptide was titrated in 10 steps from 0 to 1200µM, the non-phosphorylated peptide was titrated in 7 steps from 0 to 800 µM.

The spectra were analyzed with NMR TITAN [21] using a two-state binding model. Errors were calculated using 100 iterations of bootstrap error analysis.

#### {^1^H}-^15^N heteronuclear NOE

The pulse sequence underlying this experiment is noef3gpwg3d from the Bruker library.

{^1^H}-^15^N heteronuclear NOE (hetNOE) experiments of MEX3A were recorded at 900 MHz on a 1.1 mM sample of MEX3A(129-210) WT. The temperature was set to 298K. The recovery delay was set to 6 seconds.

Intensity ratios were calculated from peak volumes, which were extracted using PeakFit [52]. Indicated error bars were calculated based on the noise level of both spectra, using Gaussian error propagation. The intensity ratios were not disentangled to account for the impact of folding and unfolding over the course of the experiment, as suggested by Farrow et al. [53].

#### ^15^N *R*_2_-relaxation

The ^15^N R_2_-relaxation measurements used fixed-duration CPMG loops, water suppression with Watergate, and water flip-back. At 900 MHz, the loop had a duration of 16.108 ms. In 16 increments, loop iterations ranged from 1 to 20.

Peak volumes were extracted using PeakFit [52]The exponential decay function was fitted to the data using an in-house Python implementation. Average value and standard deviations of the fitted parameters were based on Monte Carlo simulations (N = 1000) [54].

#### CLEANEX

CELANEX-PM experiments were measured on a 1.2 GHz spectrometer. Interscan delay was set to 2 seconds. Delays were set to 1, 2, 5, 10, 20, 50, 70, 100, 120, and 200 ms. The pulse sequence used is fhsqccxf3gpph in the Bruker library [55]. The reference pulse sequence is fhsqcf3gpph [56].The analysis was based on peak height rather than globally fitted peak volumes.

The chemical exchange rates were extracted using an initial slope fit using 1, 2, 5, and 10 ms time points [55]. To account for the slow return of water protons’ magnetization to equilibrium, the approximation from the original publication was used: The pulse program was cropped to only include the water magnetization and then measured with 64 dummy scans and a pre-scan delay of either 2 seconds, the same as the CLEANEX experiment, or 60 seconds to allow for full relaxation of the water protons. The ratio of water proton intensity between these two experiments was used as a correction factor. At 298 K and 1.2 GHz, this correction factor was 1.514, and at 310 K, it was 1.485. Either fit was validated with Monte Carlo simulations (N = 200). The fitted values were discounted if the extracted water exchange values were not significant, as judged by the Monte Carlo-derived value minus standard deviation including the value 0, or if the fitted value correlated with another value by more than 0.75.

#### EXSY

EXSY experiments were conducted on a 900 MHz Bruker Avance III spectrometer at 310 K, using 1.4 to 1.6 mM protein, or on a 600 MHz Bruker Avance III spectrometer at 298 K using 1.1 mM protein. The recovery delay was set to 7 s. The mixing times ranged from 10 ms to 2.6 s. The intensities of all four peaks per residue were extracted by globally fitting the pseudo-3D data using the PeakFit implementation in Python [52].

Because the high exchange of amide protons with the solvent water at 310 K (**Figure S14**), only three residues with low solvent exchange, well resolved peaks over all mutants were chosen (138, 145, 146) and their peaks were used to derive their respective populations of folded and unfolded states, their respective *R*_1_ values and exchange rate, using the formulae derived by Farrow et al. [17] and an in-house Python script, which were created with support from ChatGPT 3.5.

At 298 K, the spectra were recorded at 900 MHz for MEX3A WT. The exchange rates were fit for every residue and averaged for the reported global value. Peaks that overlapped too heavily were manually removed from the analysis. Peaks that were shifted by the PeakFit implementation beyond the boundaries of the manually assigned peak were also excluded from this analysis. Only those residues were used for regression analyses for which three or four peaks were well resolved.

To compare the WT protein with the stabilized mutant T140S/T188A and the destabilized D185A, EXSY experiments were recorded at 600 MHz using 6 delays ranging from 10 ms to 1.4 s. To derive comparative values, peak heights extracted with CCPN v3 were used, rather than volumes extracted using PeakFit. The peak heights were used on data from single residues, fitting the same equations as above using an in-house python script, which were created with support from ChatGPT 3.5. The rates derived for single residues were averaged and the averaged rates were normalized to those of the WT protein. Data from these residues were used where available: 138, 144, 149, 167, 184, 188.

#### Chemical Exchange Saturation Transfer (CEST)

CEST experiments of MEX3A(129-210) were recorded on a 1.2 GHz spectrometer using B1 fields of 15 Hz and 50 Hz with the exchange delay set to 600 ms. The pulse sequence hsqc_cest_etf3gpsitc3d from the Bruker library was used [19]. The concentration of protein in the sample was 1.6 mM. The buffer was the standard NMR buffer described above. The data was recorded at 310 K, 36.85 °C. Peak volumes were extracted using PeakFit [52]. Peaks with unresolvable overlaps were excluded from the analysis. The analysis and fits were conducted using ChemEx version 2026.9.2 [57]. Only peaks of the folded state were analyzed. Residues for which the ^15^N resonance of the two states was not well separated were not included. The relaxation parameters of the two states were not fixed to be identical. Global fits of the exchange rates and populations were applied.

The 33 residues used for fitting were (**Data S1, S2**):

132, 133, 134, 140, 142, 143, 144, 145, 146, 148, 149, 152, 156, 160, 162, 164, 165, 167, 171, 172, 178, 179, 181, 182, 184, 186, 188, 189, 190, 191, 192, 195, 206.

#### H^N^ temperature coefficient analysis

^1^H,^15^N-HSQCs were measured on a 900 MHz spectrometer.

The differences in amide proton frequency between the starting temperature of 288 K and the measured temperature were correlated. The regression analysis was carried out in Python using an in-house script, which created with support from ChatGPT. As no DSS or other reference compound was used in this sample, the calculated temperature coefficients were adjusted by subtracting the temperature coefficient of water (11.9 ppb / K) as established by Wishart et al. [58].

The cutoffs used are described in detail by Cierpicki and Otlewski [30], as well as Andersen [31].

#### Structure calculation of MEX3A(108-210) T140S T188A

The structural model of MEX3A(108-210) T140S T188A was calculated using NMRtist [59]. An array of experiments was recorded on two uniformly ^1^H/^15^N/^13^C–labelled samples, processed, and uploaded. Otherwise, only the amino acid sequence was provided to the program.

The experiments included: ^1^H,^15^N-HSQC (based on Mhnhsqcwgs_dec2), ^15^N-edited NOESY, HNCACB (based on hncacbgpwg3d), CBCAcoNH (based on cbcaconhgpwg3d), HNCO (based on hncogpwg3d), HNcaCO (based on hncacogpwg3d), and C-evolved HcccoNH (based on hccconhgpwg3d3) in water. An additional H-evolved HcccoNH (based on hccconhgpwg3d2) was recorded in water, as well as a ^1^H^13^C-HMQC, and ^1^H^13^C-edited NOESY [60] in 100% D_2_O.

The two samples used had a final concentration of 780 µM and 1.6 mM, respectively. They were recorded in 20 mM NaP, pH 6.5, 150 mM NaCl, 5 mM DTT, and 5 % D_2_O. To transfer the sample into 100% D_2_O, a sample in protonated buffer was lyophilized and taken up in 100% D2O.

The experiments were recorded at 298 K on a 900 MHz spectrometer.

### In-vitro phosphorylation of MEX3A(129-210)

MEX3A(129-210) WT protein, uniformly labelled with ^15^N and labelled with ^13^C on the methyl groups of methionine, leucine, isoleucine, and valine, was used. After recording of a reference ^15^N SOFAST-HMQC experiment at 600MHz, 50 µM of the protein was buffer exchanged into the reaction buffer proposed by the supplier (20 mM MOPS pH 7.0, 1 mM EDTA, 1% Glycerol, 0.001% Brij, 0.2 mg/ml BSA, 20 mM Magnesium Acetate, 10 mM ATP, 5% D2O, 5 mM DTT).

20 µl (10 µg) of the commercial NUAK1 (also called Ark5) (GTIN 04053252678288) kinase solution was added and incubated at 30 °C for 48 hours, while the phosphorylation status was observed by ^15^N-SOFAST-HMQC measurements and ^13^C-HMQC measurements.

After the reaction was judged to be complete, based on the ^13^C HMQC spectra, the sample was buffer exchanged to standard NMR buffer (20 mM NaP, pH 6.5, 150 mM NaCl, 5 mM DTT, 5% D_2_O) and measured again.

### CD melting point analysis

For the melting point analysis using circular dichroism, the proteins of interest were buffer exchanged by repeated concentration and dilution until at least 99.5 % of the buffer was exchanged. 200 µl of a 50 or 100 µM solution of the proteins was then loaded into a quartz cuvette and measured using a J-1500 CD spectrometer. The temperature increased from 5 or 10 to 95/100 °C, increasing by 1.5 °C per minute. Over the course of the experiment, the CD was measured at either 222 nm or 220 nm to evaluate the folding state of the α-helices contained in the protein.

The resulting data was fit by minimizing this formula for a single unfolding event, using an in-house script:

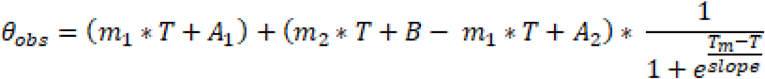

With m_1_*T+A_1_ describing a baseline of the pre-transition state, defined by a slope m_1_, the temperature T, and the intercept value A. m_2_*T+A_2_ similarly describes the baseline for the post-transition state. 1/(1+e^((T_m_-T)/(slope))) describes a logistical function to fit the transition midpoint where T_m_ is equal to T, and the slope is maximal.

Melting curves with two domains were analyzed using:

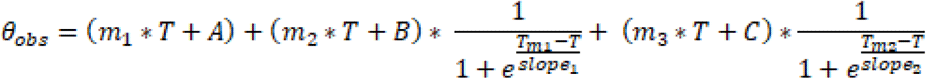

Reported errors correspond to the fitting error of a single experiment.

### Bioinformatics analyses

#### Dataset construction and phylogenetic analysis of MEX3 KH1 domains

The KH1-domain sequences of human MEX3A, MEX3B, MEX3C and MEX3D were used as BLASTp queries against UniProtKB reference proteomes, with an E-value threshold of 0.001 [61]. Retrieved homologous sequences were filtered to remove redundant and low-confidence entries, yielding a curated dataset of 34 MEX3 KH1 sequences. A homologous KH-domain sequence from Drosophila (UniProt accession Q9V4F3) was additionally included as an outgroup.

Sequences were aligned using MAFFT [62]. The resulting multiple sequence alignment was used to calculate pairwise sequence identities and infer a maximum-likelihood phylogeny using IQ-TREE 3.1.2 [63]. The best-fitting amino-acid substitution model was selected using ModelFinder (setting –m set to MFP), and branch support was assessed using 10,000 ultrafast bootstrap replicates (setting -B set to 10000). The tree was rooted using the Drosophila sequence and visualized and annotated using phylo.io [64].

#### Computational assessment of KH-domain semi-stability

To assess whether computational predictions could distinguish semi-stable KH domains from folded controls, we manually curated the Protein Data Bank (PDB) for KH domains with experimental structures determined in the absence of RNA. This yielded 20 PDB entries representing distinct KH domains: HDLBP KH1 (2CTE), KH5 (1VIG), KH8 (2CTJ), KH12 (2CTK), KH13 (2CTL) and KH14 (2CTM); FMR1 KH1 (2FMR); FUBP1 KH1 (4LIJ), KH3 (6Y2C) and KH4 (6Y24); mouse FUBP1 KH2 (1X4M); FUBP2 KH2 (2OPV), KH3 (2HH3) and KH4 (2HH2); KHDR1 KH (7ZAC); KHDR3 KH (5EL3); mouse TDRKH KH2 (1WE8); FXR1 KH1 (2CPQ); Caenorhabditis elegans MINA-1 KH3 (6FBL); and MEX3D KH2 (2DGR). These domains were used as a reference set presumed to adopt a well-folded conformation, even in the absence of RNA (apo state).

For the corresponding domain sequences, we retrieved per-residue disorder consensus scores from MobiDB [65], based on eight underlying disorder predictions. We also retrieved per-residue AlphaFold2 predicted local distance difference test (pLDDT) scores through MobiDB. These scores quantify local structural prediction confidence and were examined as an indirect indicator of disorder [66].

The reference set was supplemented with MEX3B, MEX3C and MEX3D KH1, which were shown to be folded in this study but lack experimental apo structures. The semi-stable domains MEX3A KH1, HDLBP KH7 and HDLBP KH9, identified experimentally in this study, were included for comparison. Domain sequences were aligned using MAFFT, and the per-residue scores were mapped onto the alignment to compare corresponding positions across domains.

#### Prediction of mutational effects on MEX3A KH1 stability

We assessed the predicted effects of single amino-acid substitutions on MEX3A KH1 stability using ThermoMPNN (ThermoMPNN-I google colab notebook default settings [67]), a structure-informed neural-network predictor, and Rosetta, an energy-based molecular modelling method (PyRosetta v4 default settings, [68]). Predicted stability changes were examined across the domain, with particular attention to the experimentally validated stabilizing substitutions identified through comparison with MEX3A KH1 paralogs. Negative ΔΔG values indicate predicted stabilization, whereas positive values indicate predicted destabilization. ThermoMPNN predictions are reported in kcal mol⁻¹ and Rosetta scores in Rosetta energy units (REU).

## Acknowledgments

We thank Gerd Gemmecker, BNMRZ, for help with NMR experiments, and Sebastian Metzler, BNMRZ, for discussions.

## Author contributions

Conceptualization: P.M., C.C., D.S., C.W., W.V., M.S.

Methodology: P.M., C.C., W.V., M.S.

Investigation: P.M., C.C., P.Z, C.H.

Visualization: P.M., C.C.

Supervision: W.V., M.S.

Writing—original draft: P.M., C.C., W.V., M.S.

Writing—review & editing: P.M., C.C., S.A., W.V., M.S.

## Data, code, and materials availability

Structural coordinates and restraints for the calculation of the MEX3A(108-210) T140S T188M protein are available upon request and are being deposited at the PDB and BMRB with the accession codes to be made available soon.

NMR chemical shifts for the MEX3A(129-210) and MEX3A(125-294) constructs are available upon request and are being deposited at the BMRB (54089, 54100).

