## supporting information for "Semi-stability and unfolding of a KH domain modulate RNA binding and functions of MEX3A"

**This PDF file includes:**

Figs. S1 to S20
Tables S1 to S2
Data S1 to S2

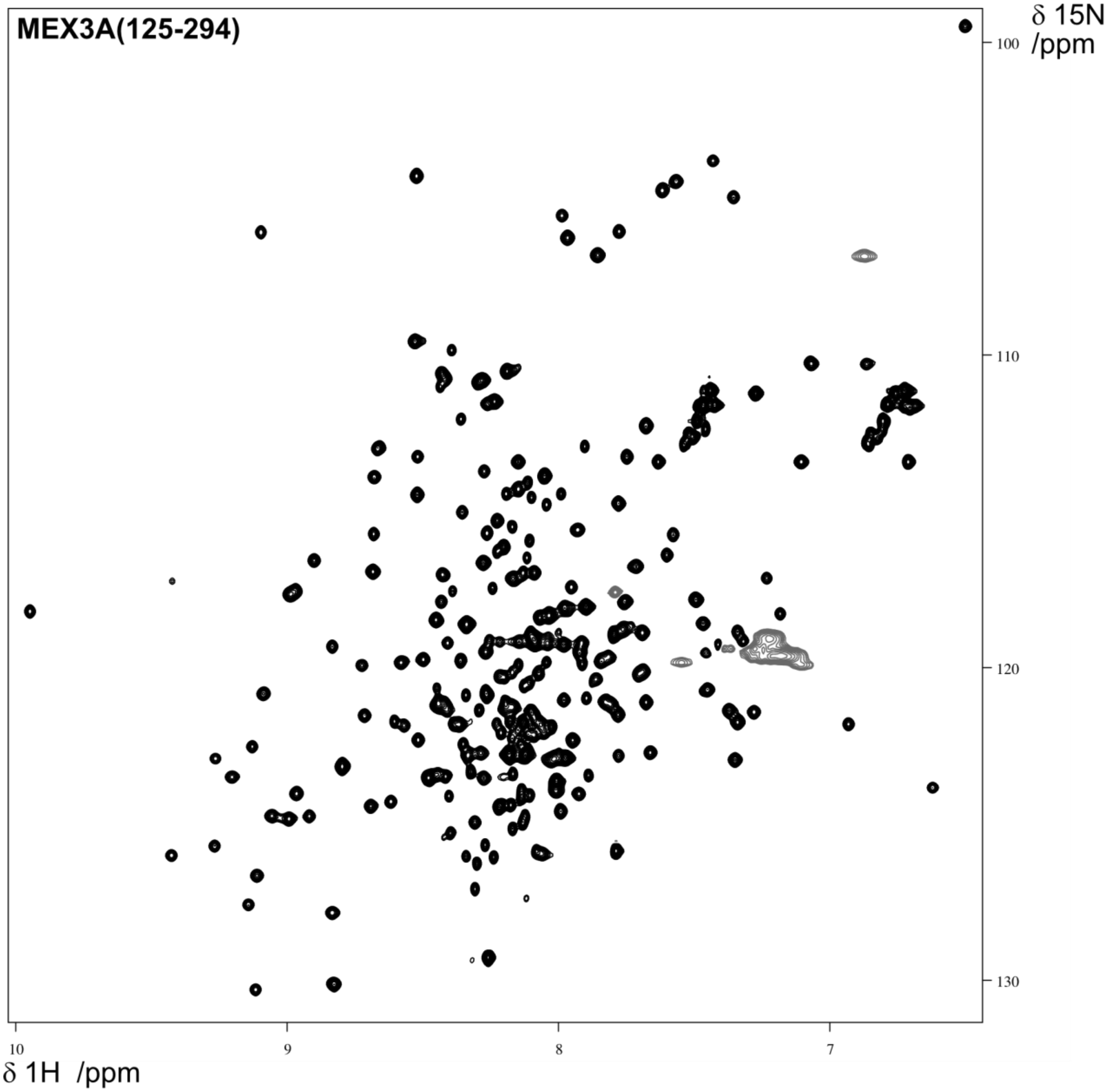


**Fig. S1. HSQC of MEX3A(125-294)**

HSQC experiment of the KH1/2 construct recorded at 298 K.

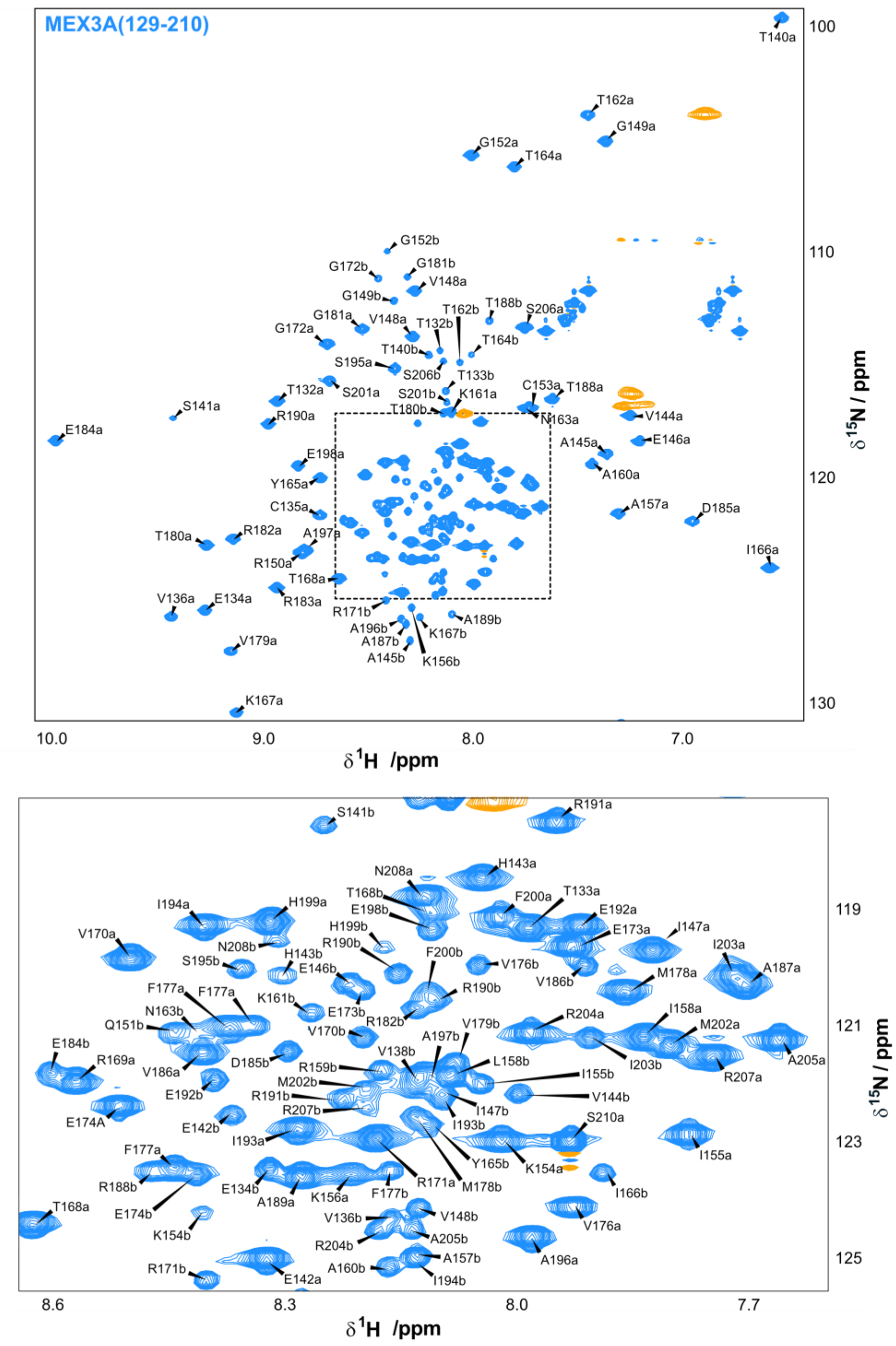


**Fig. S2. Amide backbone assignment of MEX3A(129-210) WT**

A residues assignment in the folded state is suffixed with a lower case ‘a’, while the same residue in the unfolded state is suffixed with a lower case ‘b’. Lower intensities of the unfolded state peaks are apparent, as is the limited dispersion of those peaks.

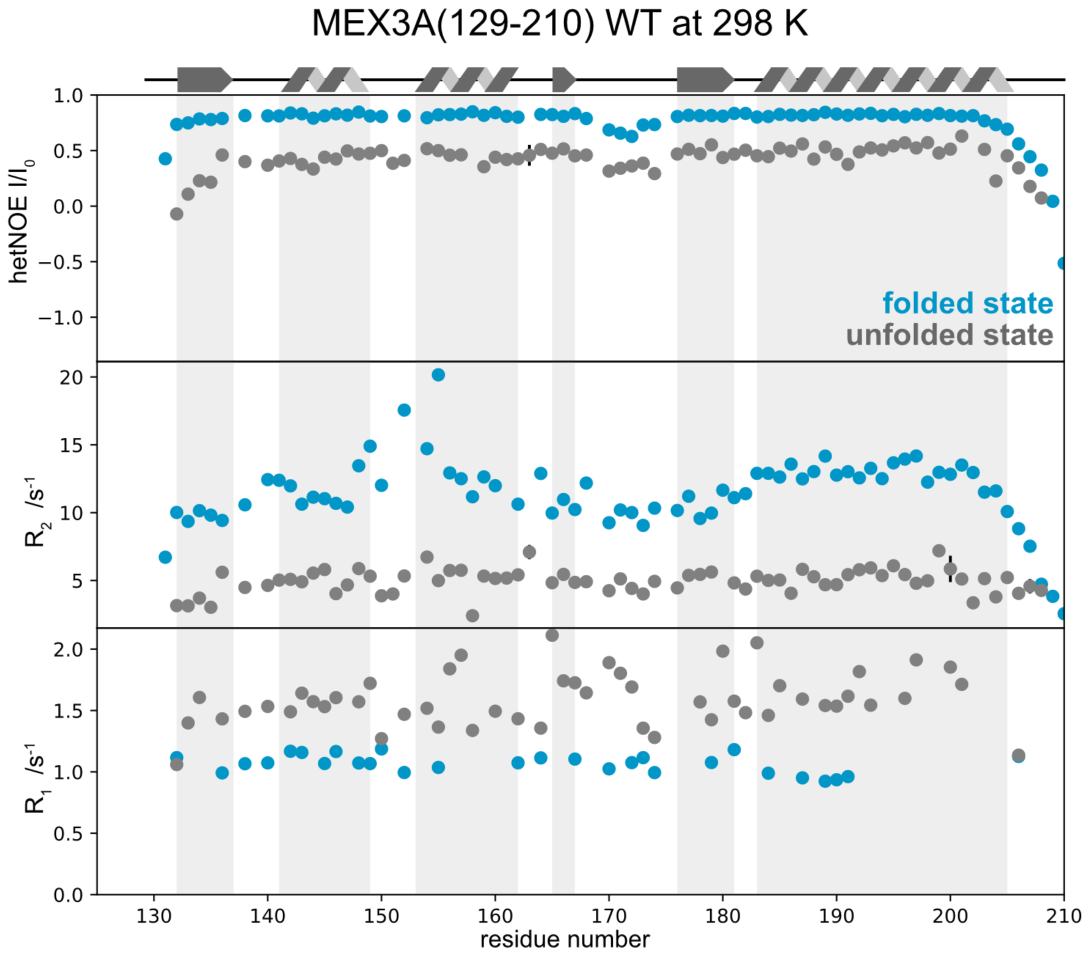


**Fig. S3. {$^1$H}-$^{15}$N NOE (hetNOE), $^{15}$N $R_2$ and $^{15}$N $R_1$ of MEX3A(129-210) WT**

MEX3A(129-210) at 298K at 900 MHz. Values are reported for the folded (blue) and unfolded state (grey). $R_1$-values were derived from an EXSY experiment. Error bars for the {$^1$H}-$^{15}$N NOE are based on the errors of the peak volumes as reported by PeakFit. Error bars for $R_2$ and $R_1$ are based on Monte Carlo simulations, which were in turn based on errors of the peak volumes as reported by PeakFit.

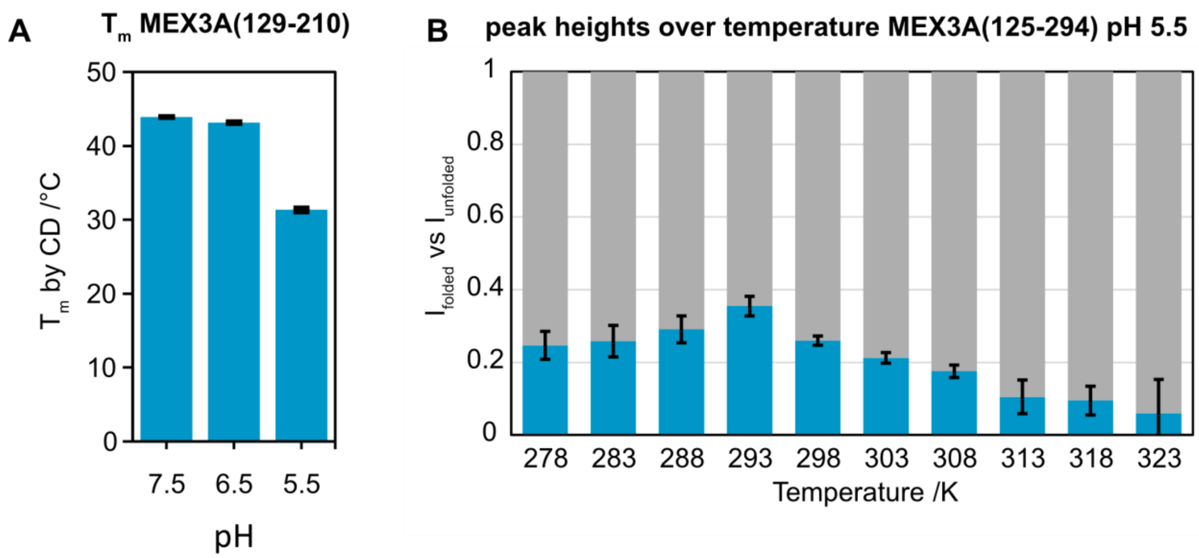


**Fig. S4. MEX3A KH1 folding equilibrium is sensitive to pH and temperature.**

**(A)** Melting temperature measured by CD spectroscopy: 317.1 K/316.3 K/304.5 K for pH 7.5/6.5/5.5.

**(B)** Peak height from an HSQC of MEX3A(129-210) at pH 5.5 compared over different temperatures. At this pH the protein adopts the folded state the most at 293 K. The cold denaturation is apparent below this temperature. Error bars are the standard deviation of the folding ratio of all measured residues. The errors are generally lower than at 6.5, likely due to lowered amide proton exchange with the solvent.

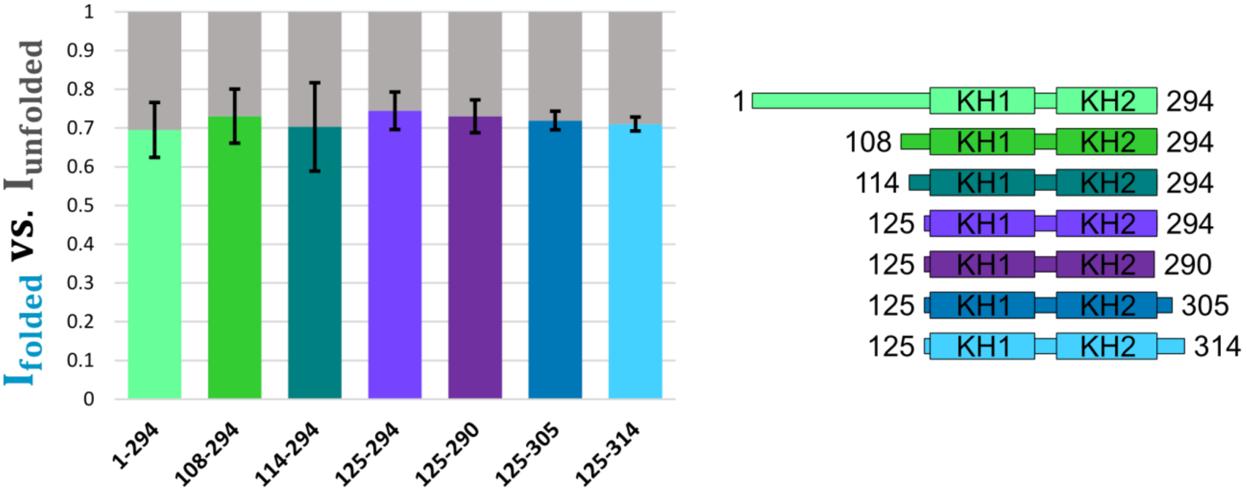


**Fig. S5. Semi-stability is not an artifact of ill-chosen construct boundaries**

Peak height comparison of the folded and unfolded state of different MEX3A KH domain construct. These range from including the entire unstructured N-terminus to reaching 20 residues past the KH2 boundaries in the C-terminal direction. Constructs that further elongated at the C-terminus are unstable, express at low yields and have increasingly crowded HSQC spectra, making this analysis less tenable.

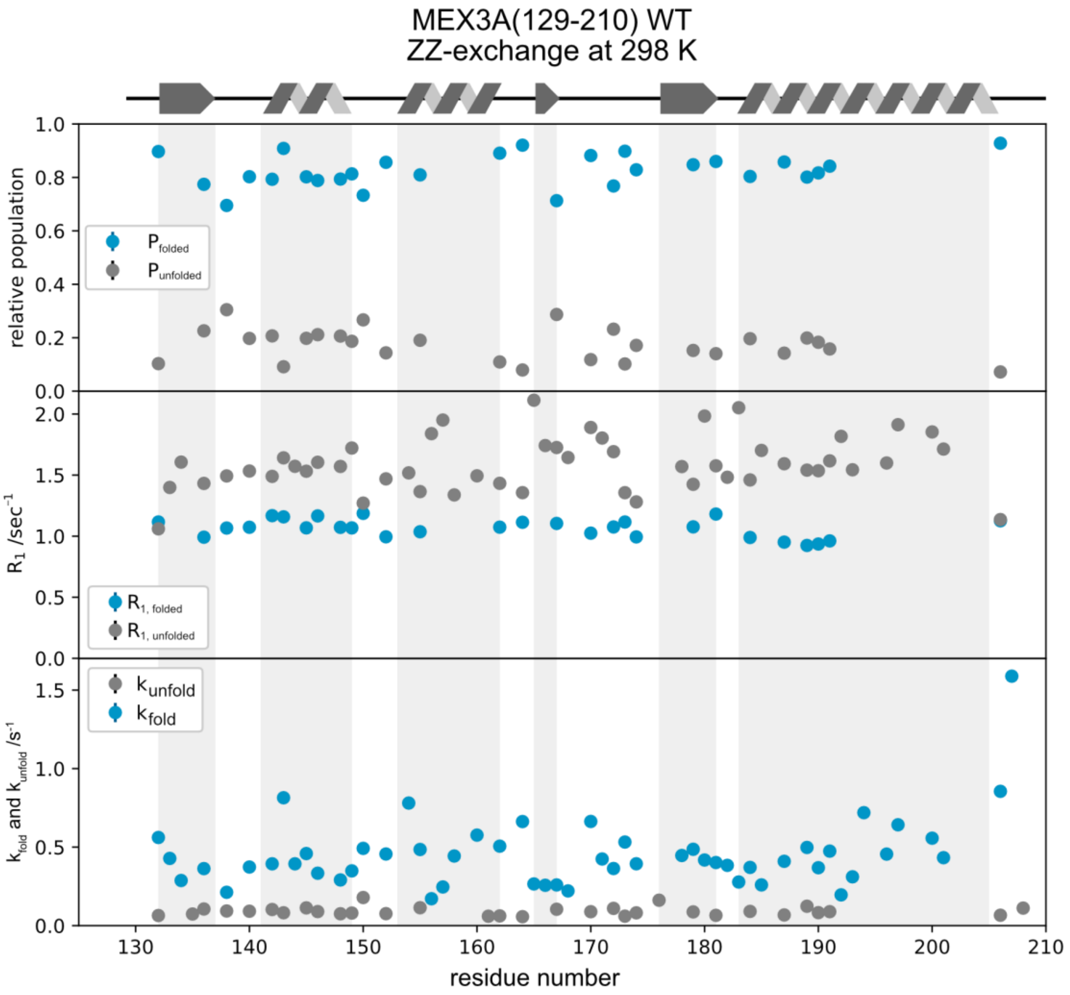


**Fig. S6. EXSY derived values of MEX3A(129-210)**

All significant values resulting from the single-residue fit of EXSY data of MEX3A(129-210) at 298K at 900 MHz.

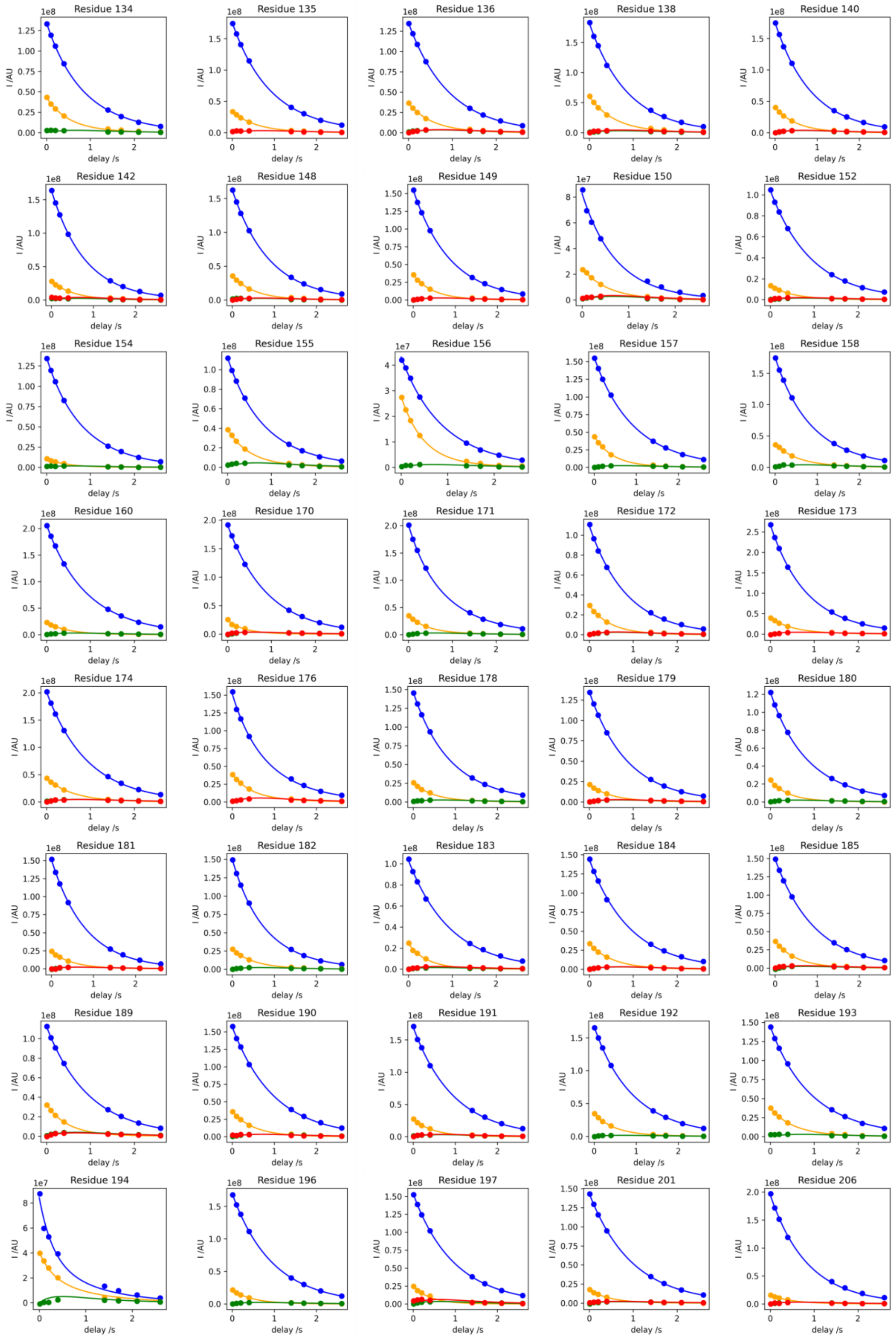
Residue 134
Residue 135
Residue 136
Residue 138
Residue 140
Residue 142
Residue 148
Residue 149
Residue 150
Residue 152
Residue 154
Residue 155
Residue 156
Residue 157
Residue 158
Residue 160
Residue 170
Residue 171
Residue 172
Residue 173
Residue 174
Residue 176
Residue 178
Residue 179
Residue 180
Residue 181
Residue 182
Residue 183
Residue 184
Residue 185
Residue 189
Residue 190
Residue 191
Residue 192
Residue 193
Residue 194
Residue 196
Residue 197
Residue 201
Residue 206
I /AU
delay /s

**Fig. S7. Single residue EXSY derived values of MEX3A(129-210) at 298 K**

Intensity in arbitrary units on the y-axis. Delay in seconds on the x-axis. Dots represent data, curves represent fits: Folded state (blue), unfolded state (yellow) and exchange peaks (red and green).

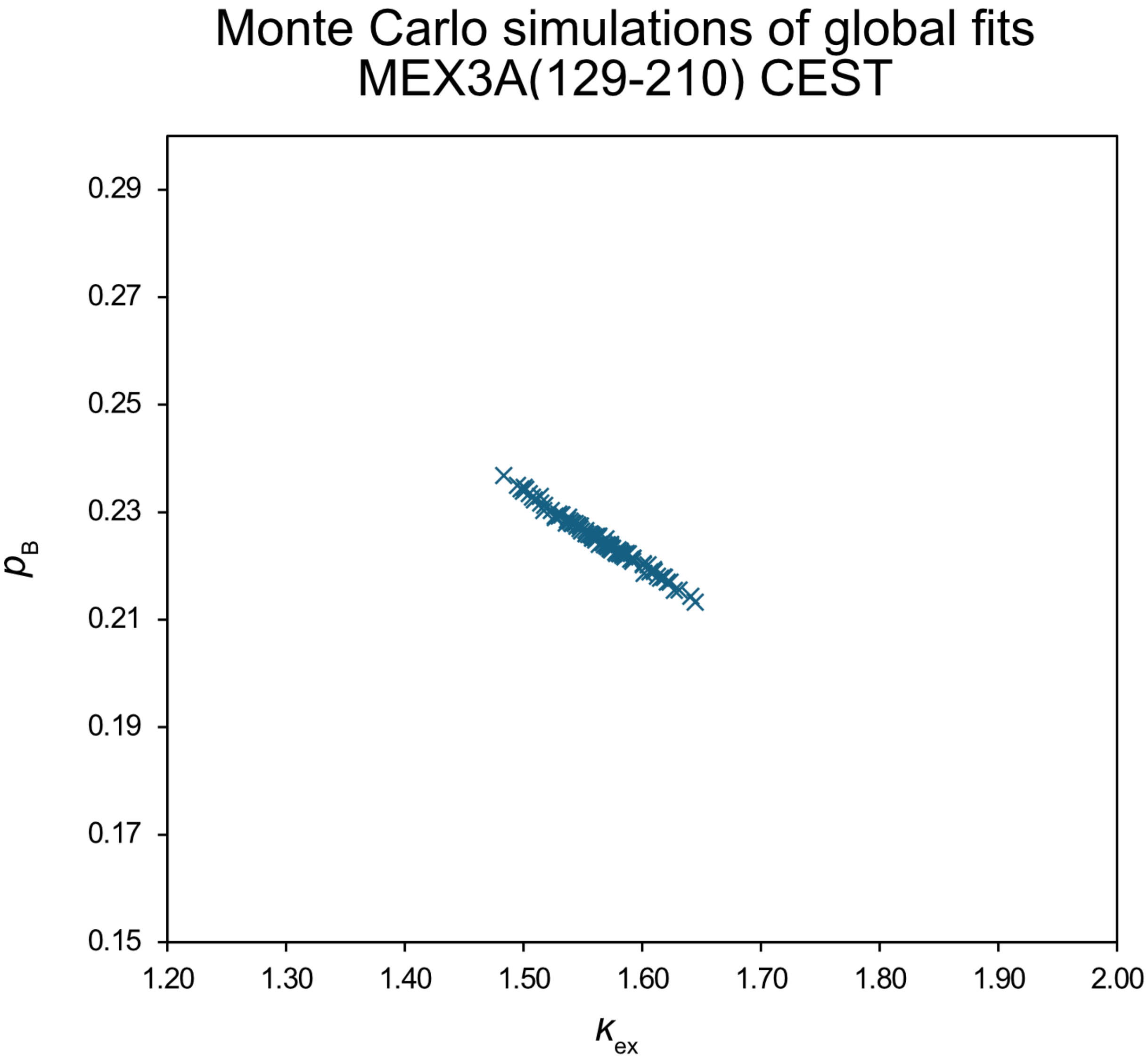


**Fig. S8. Chemical Exchange Saturation Transfer Monte Carlo calculations**

100 Monte Carlo simulations were used based on the CHEMEX scatter errors, rather than the spectra's signal to noise.

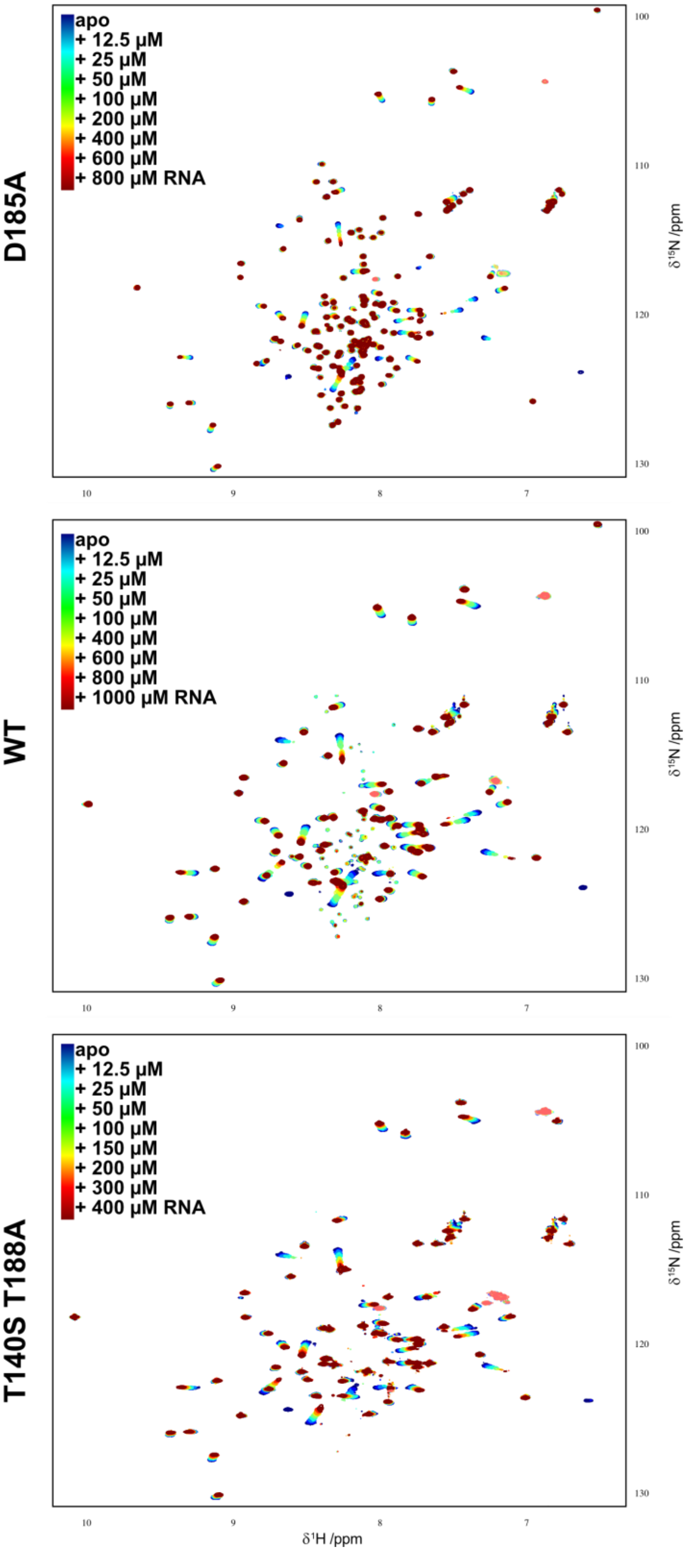


**Fig. S9. Full HSQC spectra of the titration of MEX3A KH1 with miRNA-126-5p(15-19) (GUACG)**

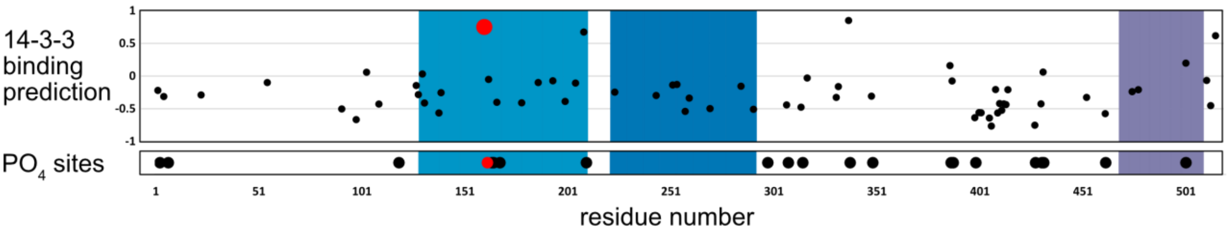


**Fig. S10. Prediction of 14-3-3 binding to and phosphorylation of MEX3A KH1**

The binding prediction was conducted for every threonine and serine in MEX3A's sequence using 14-3-3-Pred with higher values indicating a higher likelihood of a 14-3-3 binding site around the potential phosphorylation site. Phosphorylations represent every site annotated for MEX3A in EPSD, including those from ambiguously designed peptides. T162 is highlighted in red. Background color indicates protein domains: KH1 (light blue), KH2 (dark blue) and the RING-type Zinc finger (purple).

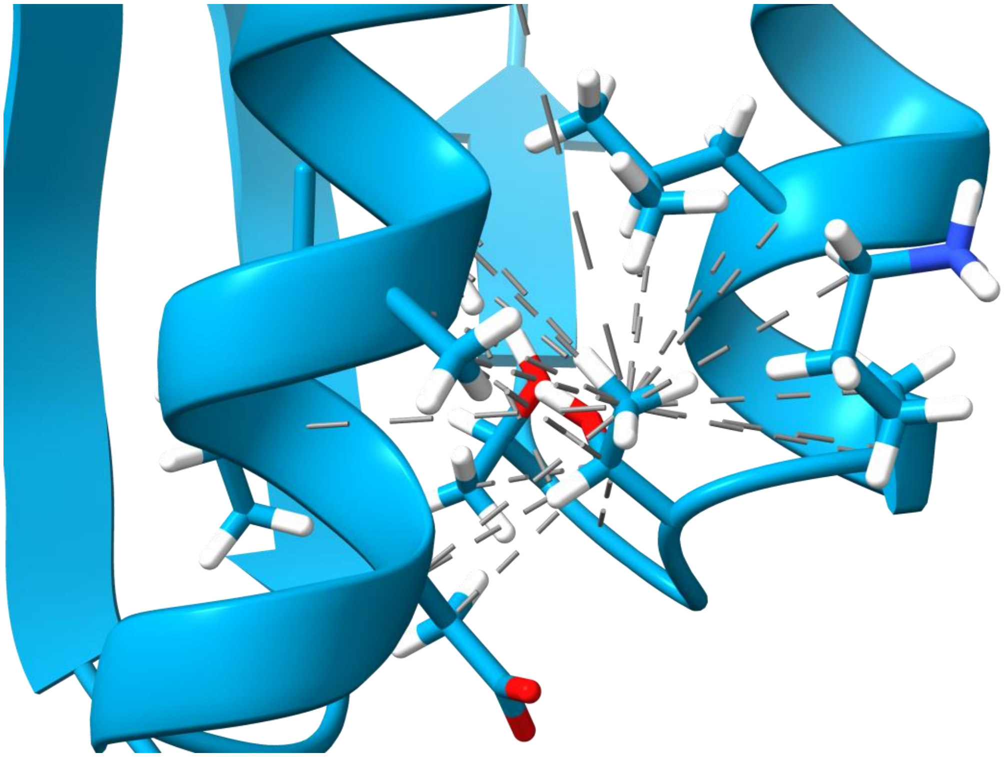

**Fig. S11. NOE restraints on the T162 side chain**

Restraints from the NMRtist automatic structure calculation that involve the side chain of T162. Connections are displayed from and to heteroatoms to also display restraints linked to equivalent protons.

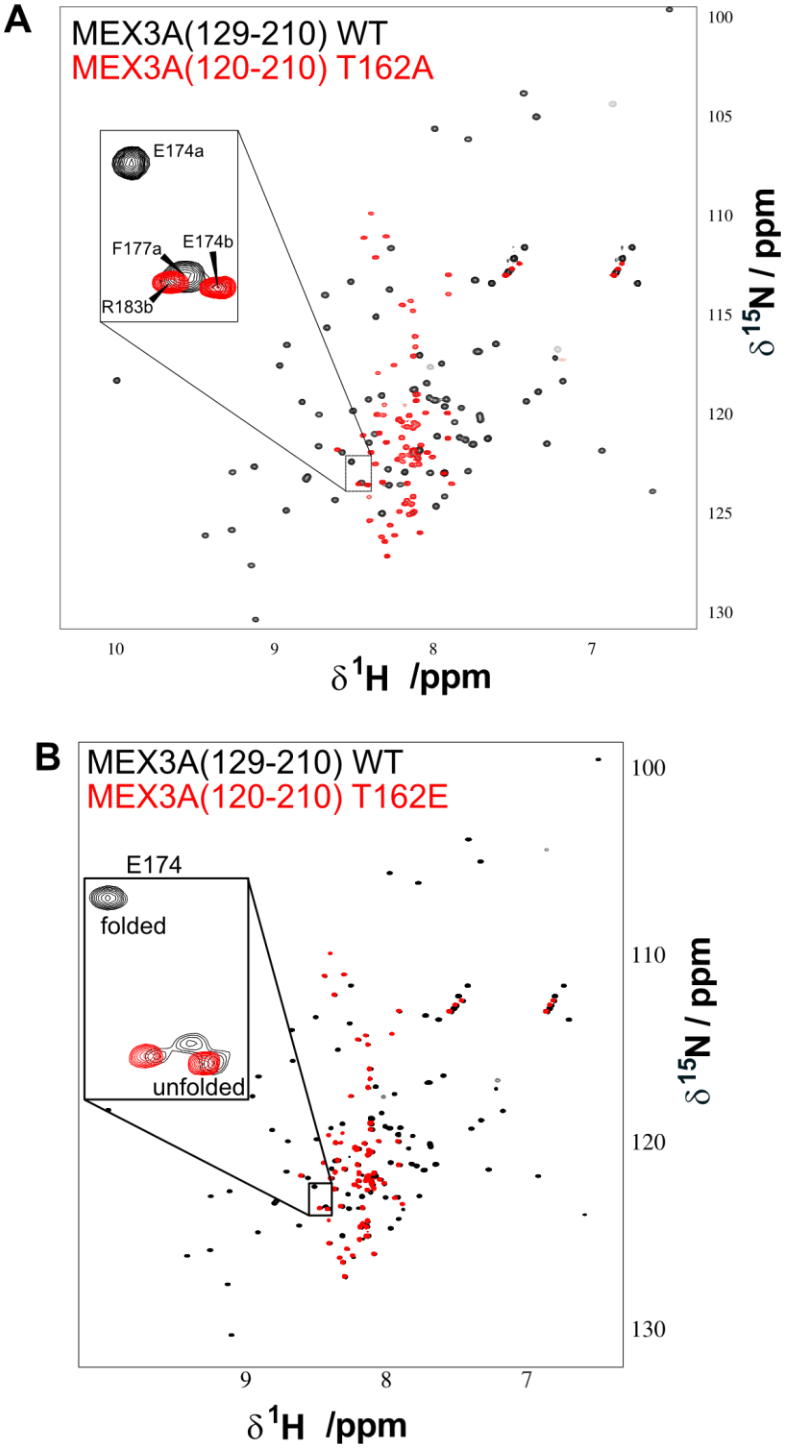


**Fig. S12. T162 of MEX3A is structurally important to the domain**

HSQC NMR experiments of MEX3A(129-210), the KH1 domain, shows two peak sets for the WT protein, but only the unfolded state for the two mutants T162A **(A)** and T162E **(B)**.

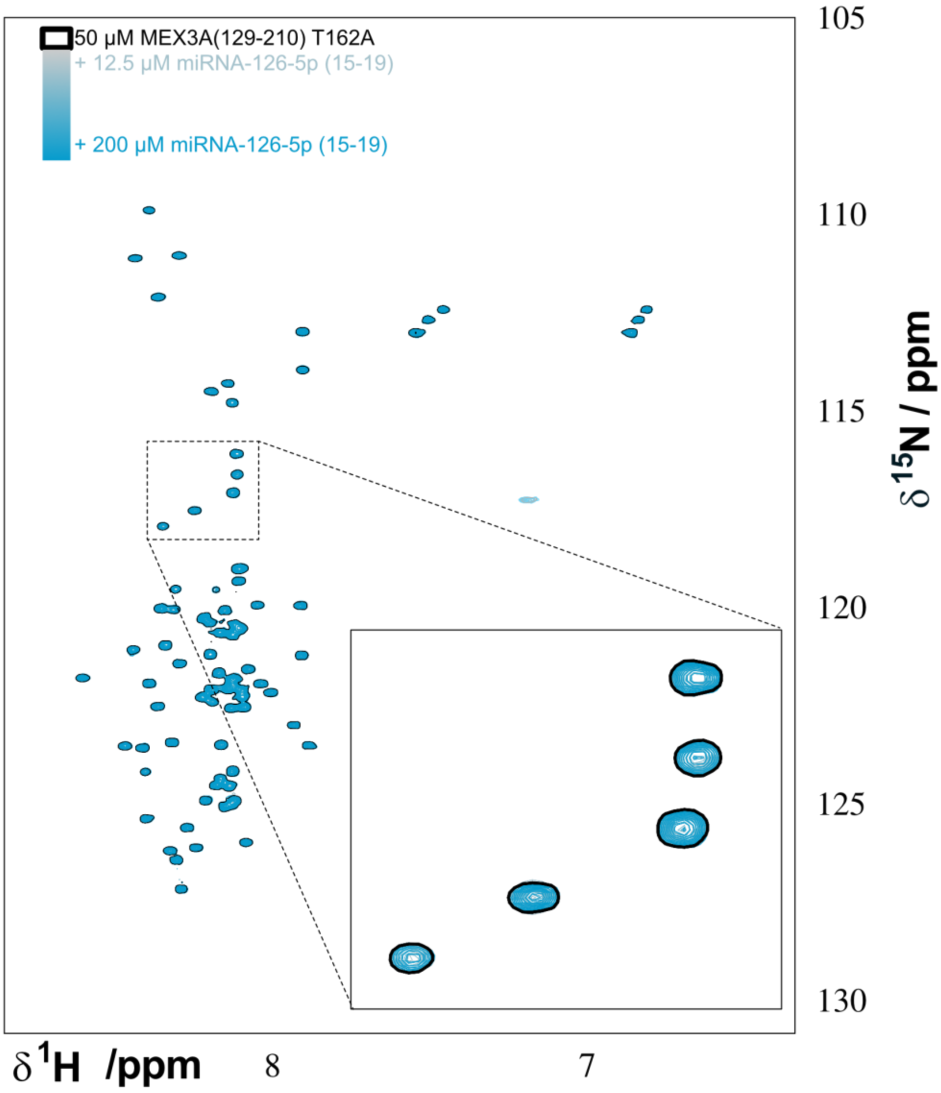


**Fig. S13. MEX3A KH1's unfolded state is not RNA-binding competent.**

The HSQC of MEX3A(129-210) T162A, a fully disordered mutant, does not show peak broadening or chemical shift perturbation upon titration with miRNA-126-5p(15-19) (GUACG).

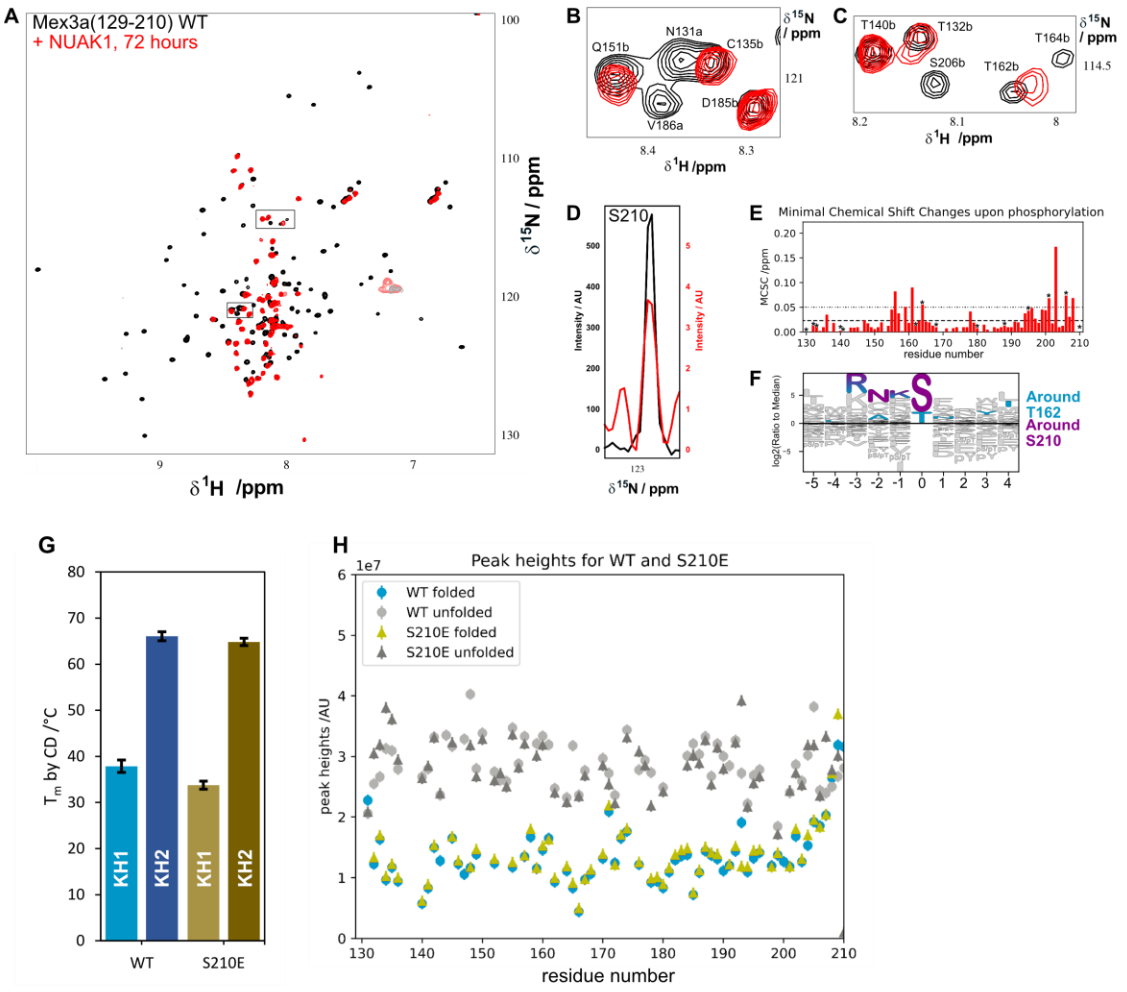

**Fig. S13.**

**NUAK1 phosphorylates two sites on MEX3A KH1. Only one influences stability**

**(A)** MEX3A can be phosphorylated by NUAK1, which unfolds the protein.
**(B)** Zoom-ins showing the fold abolished in the phosphorylated state.
**(C)** Zoom-in on T162b. How much the peak shifts upon phosphorylation could only be determined upon further resonance assignment. The new peak appearing close the T162b in the phosphorylated state likely represents a different residue and the proximity of the peaks is due to the poor dispersion of the peaks in the disordered state.
**(D)** 1D slice of S210 in the apo state and the phosphorylated state, for which the intensity was increased by a factor of 100. This suggests that S210 was phosphorylated.
**(E)** Minimal Chemical Shift Changes upon phosphorylation along the sequence of MEX3A(129-210). The pattern is consistent with the phosphorylation of S210 and T162.
**(F)** Recognition sequence of NUAK1, colored purple for the sequence leading up to S210 and light blue for the sequence surrounding T162.
**(G)** The phosphomimetic S210E has little effect on the stability of MEX3A KH1, as judged by CD melting point analysis.
**(H)** The HQCS peak hights of the folded and unfolded state are not meaningfully different between S210E and WT. The unfolding observed upon phosphorylation with NUAK1 is thus likely due to the phosphorylation of T162, not S210.

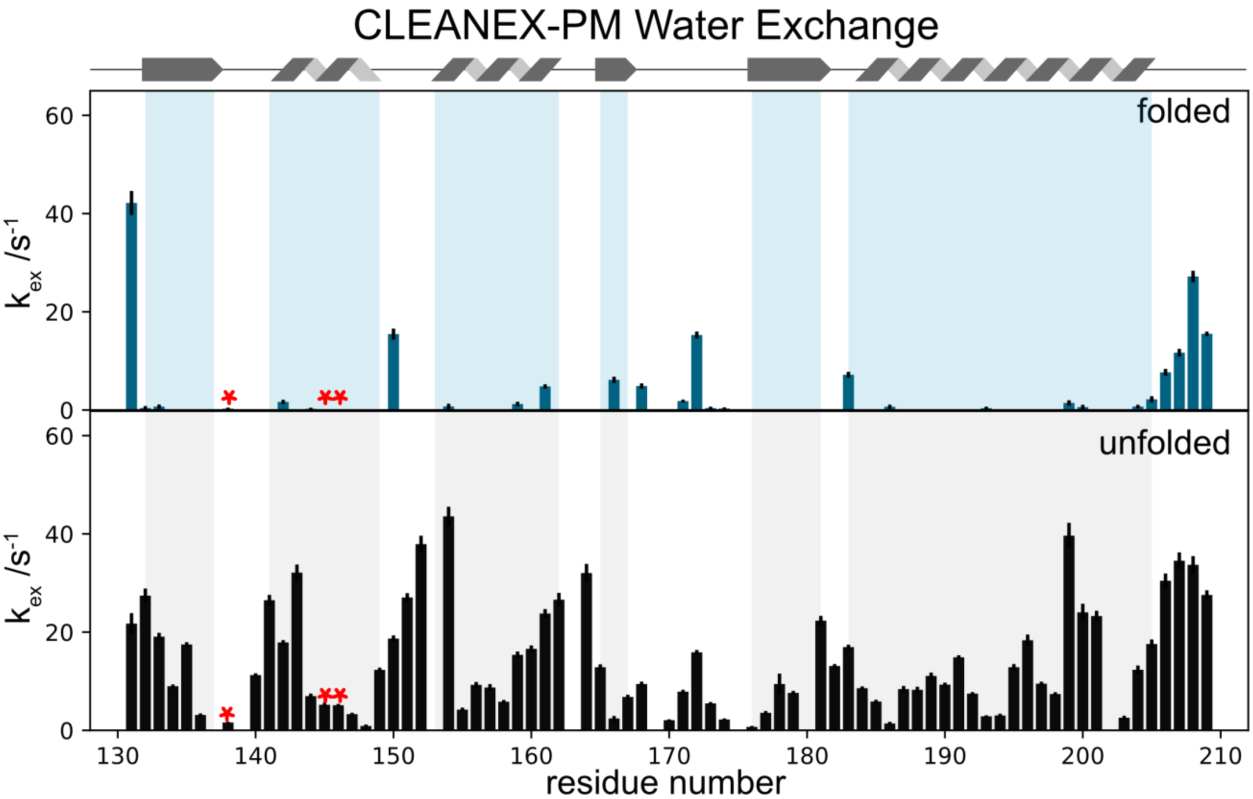


**Fig. S14. Water exchange rates of amide protons of MEX3A(129-210) in the folded and unfolded state**

Amide proton solvent exchange measured at 310 K on a 1.2 GHz spectrometer and analyzed using an initial slope fit using 1, 2, 5 and 10 ms delays. Errors from 200 Monte Carlo Simulations. Red stars indicate residues used to quantify EXSY experiments at 310 K.

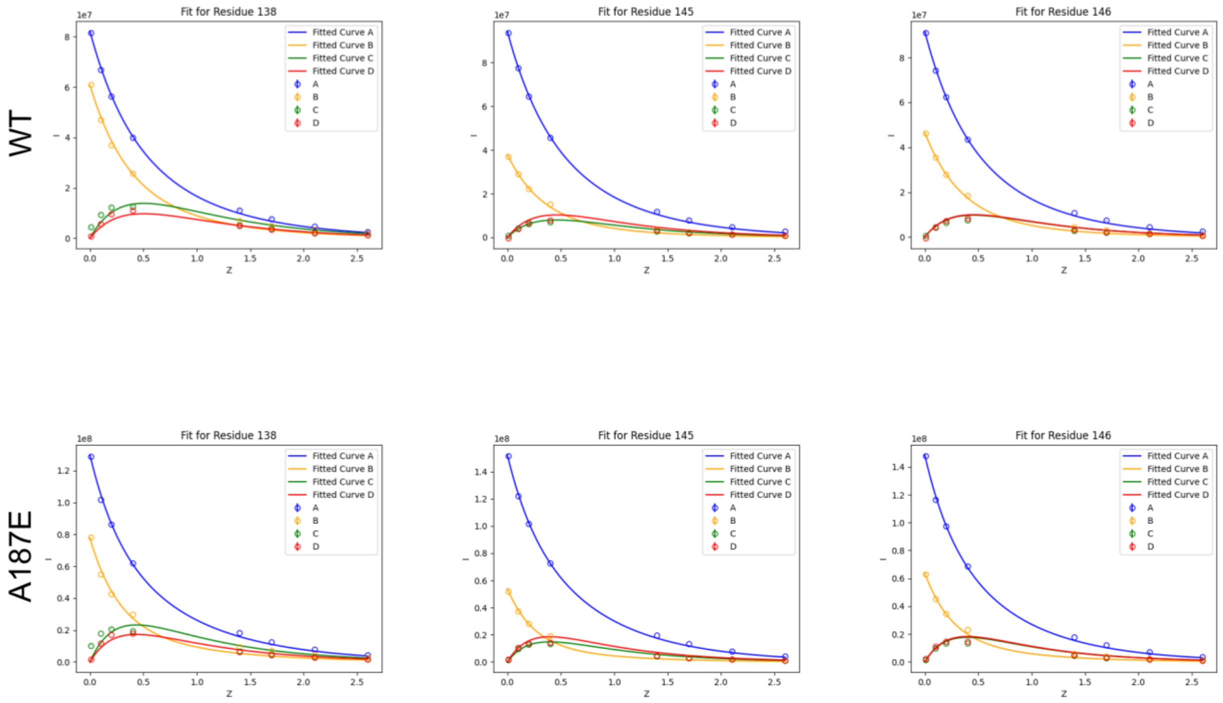


**Fig. S15. MEX3A(129-210) WT and A187E EXSY for select residues at 310K**

Dots represent data, curves represent fits: Folded state (blue), unfolded state (yellow) and exchange peaks (red and green).

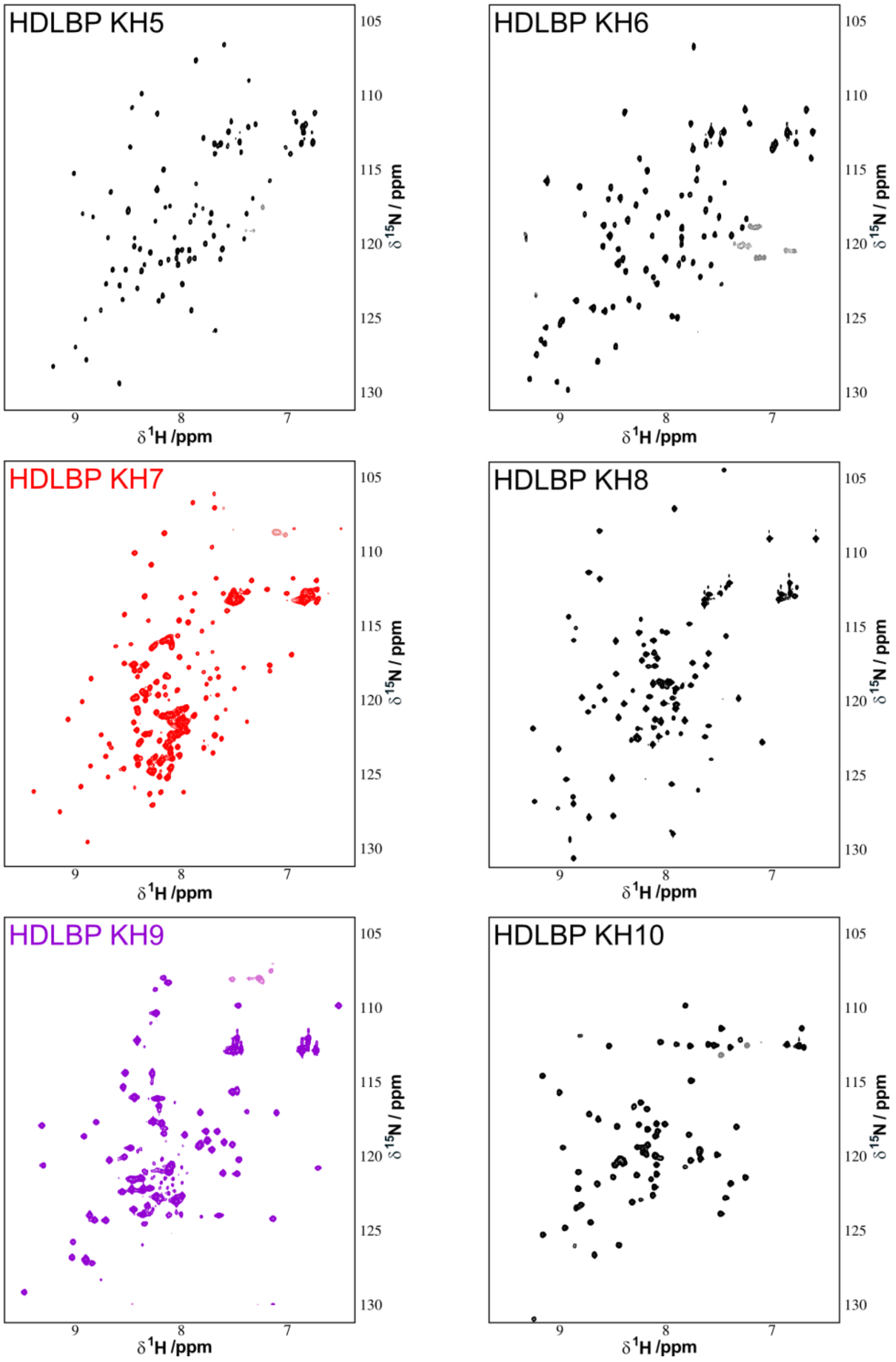


**Fig. S17. HSQCs of 6 KH domains from HDLBP indicate two as semi-stable**

KH5, KH6, KH7 and KH10 were recorded at 298 K in 20 mM NaP pH 6.6, 500 mM on a 600 MHz spectrometer. KH8 was recorded at 298 K in 20 mM HEPES pH 7.2, 500 mM on a 1.2 GHz spectrometer. KH9 was recorded at 298 K in 20 mM NaP pH 6.6, 500 mM on a 1.2 GHz spectrometer.

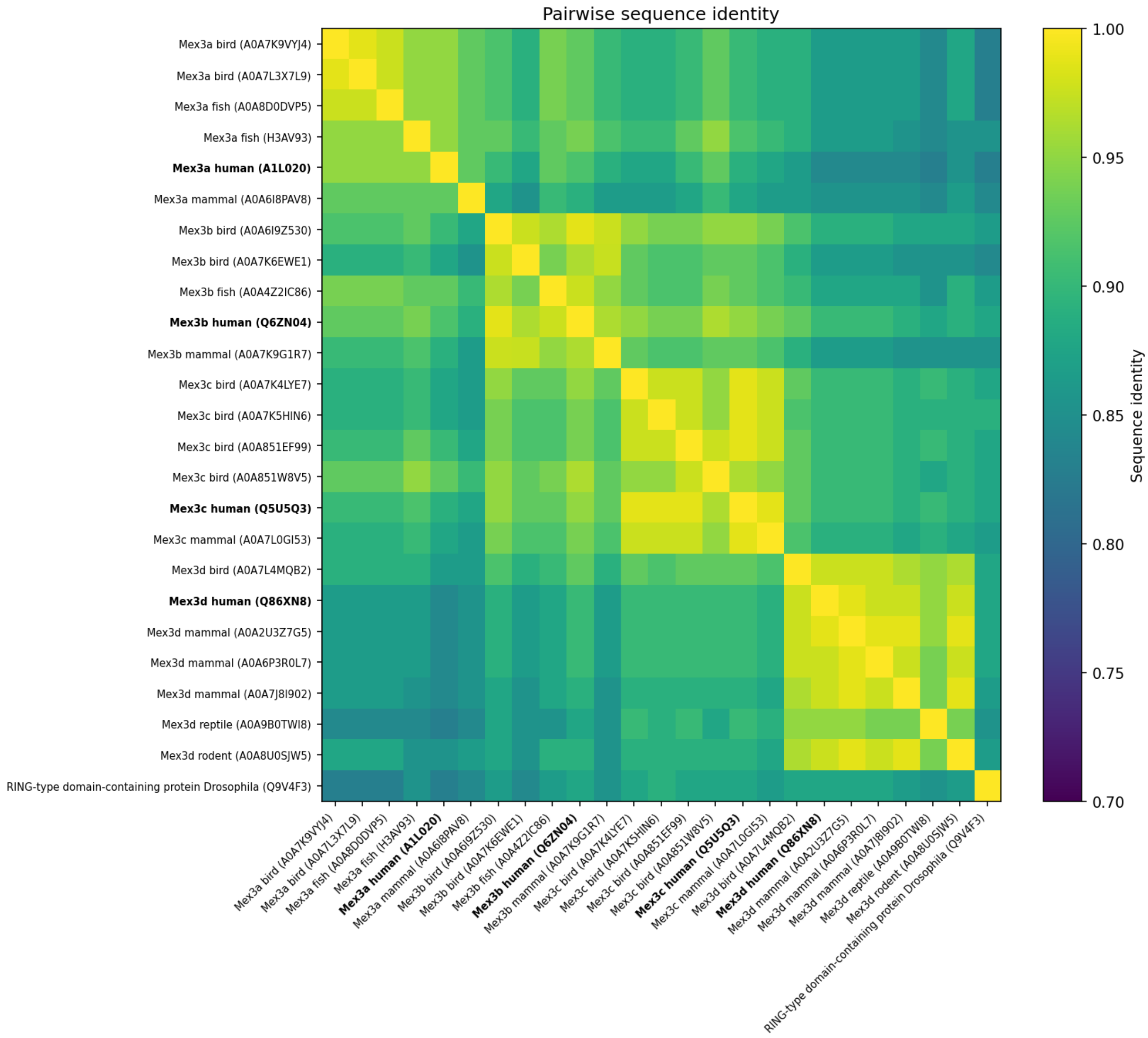


**Fig. S18. Pairwise sequence identity among selected MEX3 KH1 homologs.**

The heatmap shows pairwise amino-acid sequence identities calculated from a MAFFT alignment of selected KH1 sequences identified by BLASTp searches. The four human MEX3 paralogs are highlighted in bold. KH1 sequences show high conservation both within and between MEX3 paralog groups. The Drosophila homolog Q9V4F3, used as an outgroup for phylogenetic analysis, generally shares lower sequence identity with the vertebrate sequences

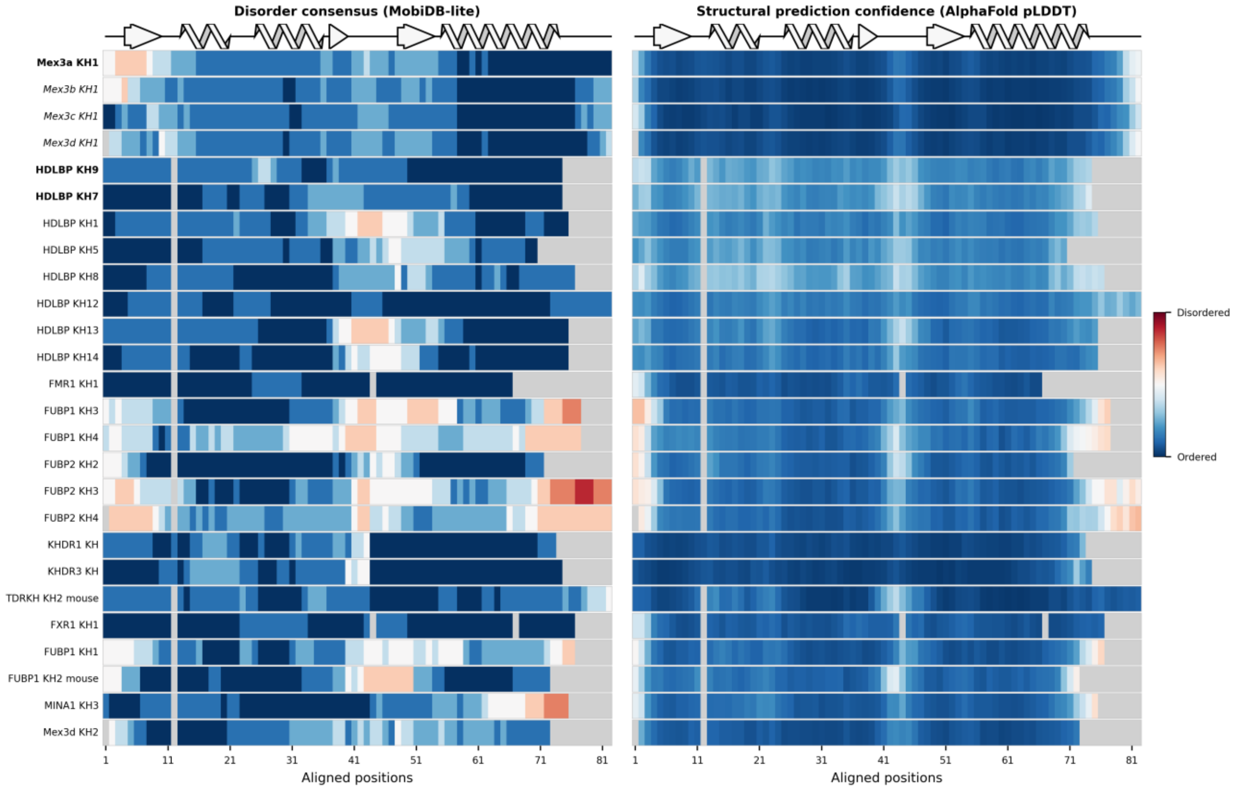


**Fig. S19. Comparison of features associated with well-folded and semi-stable KH domains.** Heatmaps show per-residue disorder consensus scores retrieved from MobiDB(left), and AlphaFold2 pLDDT scores (right). MobiDB-lite disorder consensus scores represent the fraction of eight underlying predictors that classify each residue as disordered. Scores range from 0 (all predictors classify the residue as ordered; blue) to 1 (all classify it as disordered; red). Higher pLDDT scores indicate greater confidence in the predicted local structure; lower scores can be associated with disorder. The reference set comprises KH domains with experimental structures determined in the absence of RNA, supplemented with MEX3B, MEX3C and MEX3D KH1, which were shown to be folded in this study but lack experimental apo structures. Experimentally identified semi-stable domains are labelled in bold, and the additional MEX3A **K**H1 paralogs are labelled in italics. Scores are mapped onto a MAFFT sequence alignment, with positions referenced to MEX3A KH1. Grey cells indicate alignment gaps.

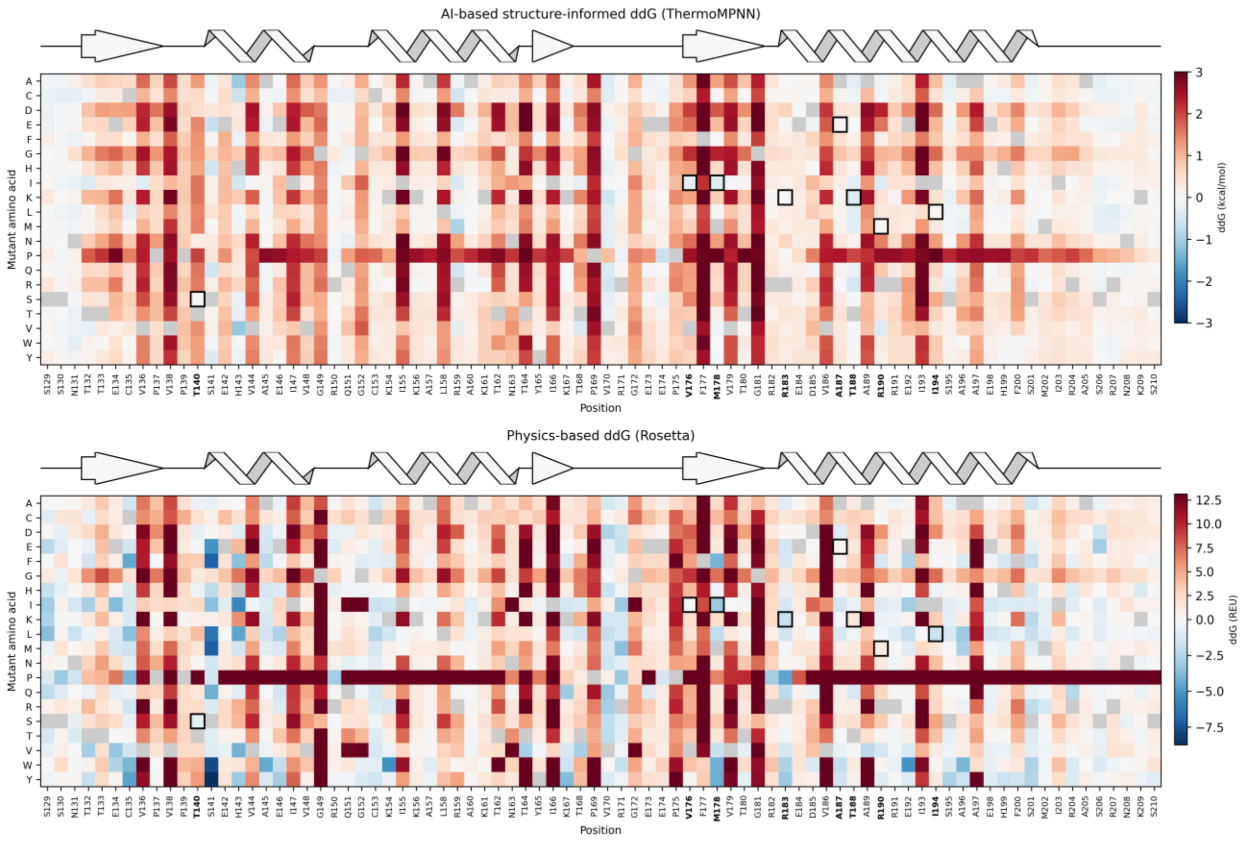


**Fig. S20. Predicted effects of single amino-acid substitutions on MEX3A KH1 stability.**

Heatmaps show predicted stability changes for single amino-acid substitutions across MEX3A KH1 using ThermoMPNN (top) and Rosetta (bottom). Columns indicate residue positions and wild-type amino acids, and rows indicate substituted amino acids. Negative values indicate predicted stabilization (blue), whereas positive values indicate predicted destabilization (red). Black outlines mark experimentally validated stabilizing substitutions.

**Table S1.**

NMR structure and refinement statistics for the NMRtist calculated 20-structure ensemble of MEX3A(108-210) T140S T188A.

| | |
|---|---|
| **NMR constraints** | |
| Total NOE | 1517 |
| Intraresidue | 423 |
| Interresidue | |
| Sequential (\|i-j\| = 1) | 391 |
| Medium range (1 < \|i-j\| <5) | 239 |
| Long range ((\|i-j\| ≥5) | 464 |
| Number of torsion angle restraints | 156 |
| **Structure statistics** | |
| Violations (mean and SD) | |
| Distance restraint violations > 0.2 Å | 7 ± 2 |
| Maximal distance restraint violation | 0.17 ± 0.04 Å |
| Angle restraint violations | 5 ± 0 |
| Maximal angle restraint violation | 21.54 ± 0.22 ° |
| Ramachandran plot | |
| Residues most favored | 87.60% |
| Residues additionally allowed | 12.10% |
| Residues in generously allowed regions | 0.20% |
| Residues in disallowed regions | 0.10% |
| Average pairwise r.m.s. Devation (20 structures) (Å) | 0.8 Å |

**Table S2. Inter residue NOE restraints involving the sidechain of MEX3A T162**

| sequence code | residue name | atom name | sequence code | residue name | atom name | upper limit |
|---|---|---|---|---|---|---|
| 162 | THR | HB | 189 | ALA | HB% | 4.37 |
| 162 | THR | HG2% | 189 | ALA | HA | 3.29 |
| 162 | THR | HB | 164 | THR | HG2% | 3.43 |
| 162 | THR | HB | 185 | ASP | HA | 3.85 |
| 162 | THR | HB | 185 | ASP | HBy | 4.24 |
| 162 | THR | HG2% | 189 | ALA | HB% | 3.12 |
| 161 | LYS | HBy | 162 | THR | HG2% | 3.94 |
| 158 | LEU | HDy% | 162 | THR | HG2% | 3.9 |
| 162 | THR | HB | 185 | ASP | HBx | 4.56 |
| 162 | THR | HB | 179 | VAL | HGx% | 5.5 |
| 162 | THR | HG2% | 185 | ASP | HA | 4.05 |
| 162 | THR | HG2% | 192 | GLU | HBx | 4.75 |
| 161 | LYS | HBx | 162 | THR | HG2% | 4.53 |
| 162 | THR | HG2% | 188 | ALA | HB% | 3.11 |
| 162 | THR | HB | 188 | ALA | HB% | 3.94 |
| 158 | LEU | HG | 162 | THR | HG2% | 3.25 |
| 158 | LEU | HA | 162 | THR | HG2% | 3.94 |
| 162 | THR | HG2% | 186 | VAL | HA | 5.5 |
| 161 | LYS | HA | 162 | THR | HG2% | 5.08 |
| 162 | THR | HG2% | 188 | ALA | HA | 5.5 |
| 158 | LEU | HDx% | 162 | THR | HG2% | 3.9 |
| 162 | THR | HG2% | 189 | ALA | H | 3.37 |
| 161 | LYS | H | 162 | THR | H | 2.73 |
| 162 | THR | HG2% | 164 | THR | H | 4.81 |
| 161 | LYS | H | 162 | THR | HG2% | 4.2 |
| 162 | THR | HG2% | 192 | GLU | H | 5.5 |
| 158 | LEU | HD% | 162 | THR | HG2% | 2.38 |
| 161 | LYS | HE% | 162 | THR | HG2% | 3.39 |

**Data S1. (separate file)**
MEX3A-129-210-WT-CEST-15hz_A.pdf
Data, fitted curve and residual for every residue of the folded state used in the CEST experiment at $B_1$ of 15Hz

**Data S2. (separate file)**
MEX3A-129-210-WT-CEST-50Hz_A.pdf
Data, fitted curve and residual for every residue of the folded state used in the CEST experiment at $B_1$ of 50Hz
