## Supplemental Data 1 for "Semi-stability and unfolding of a KH domain modulate RNA binding and functions of MEX3A"

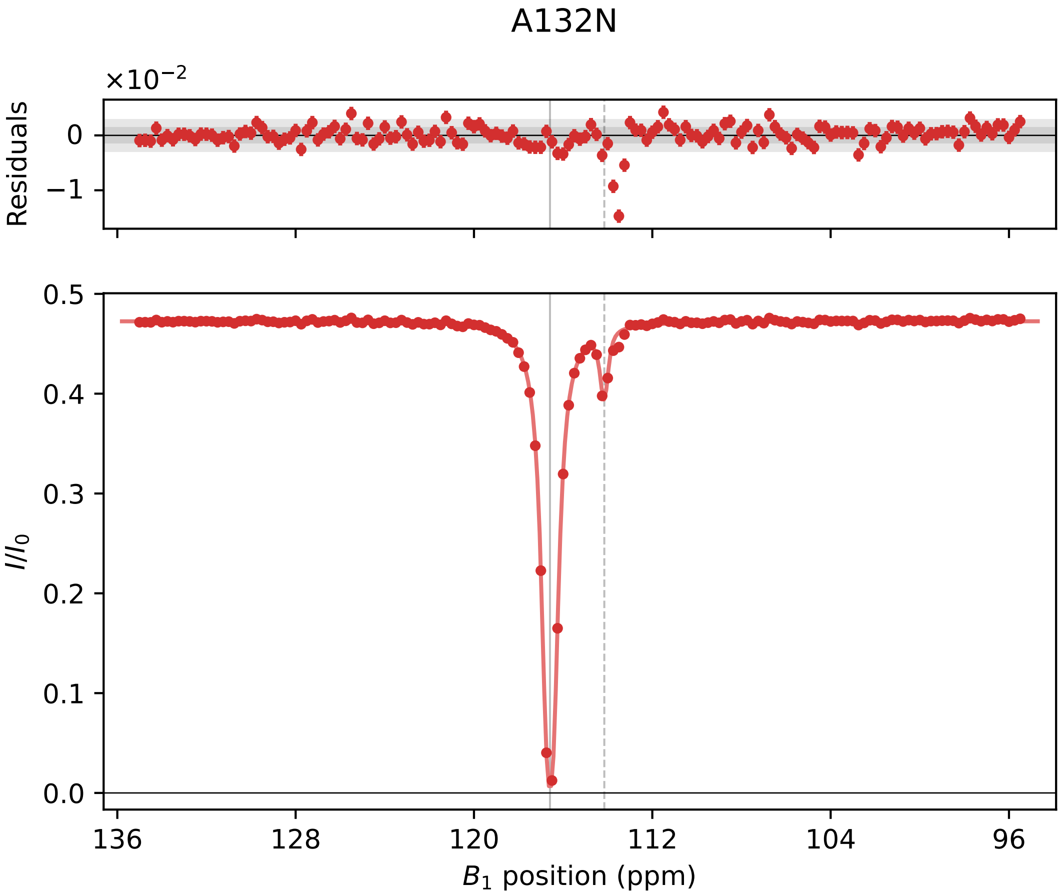

A132N
×10−2
Residuals
0
−1
I/I0
0.5
0.4
0.3
0.2
0.1
0.0
136
128
120
112
104
96
B1 position (ppm)

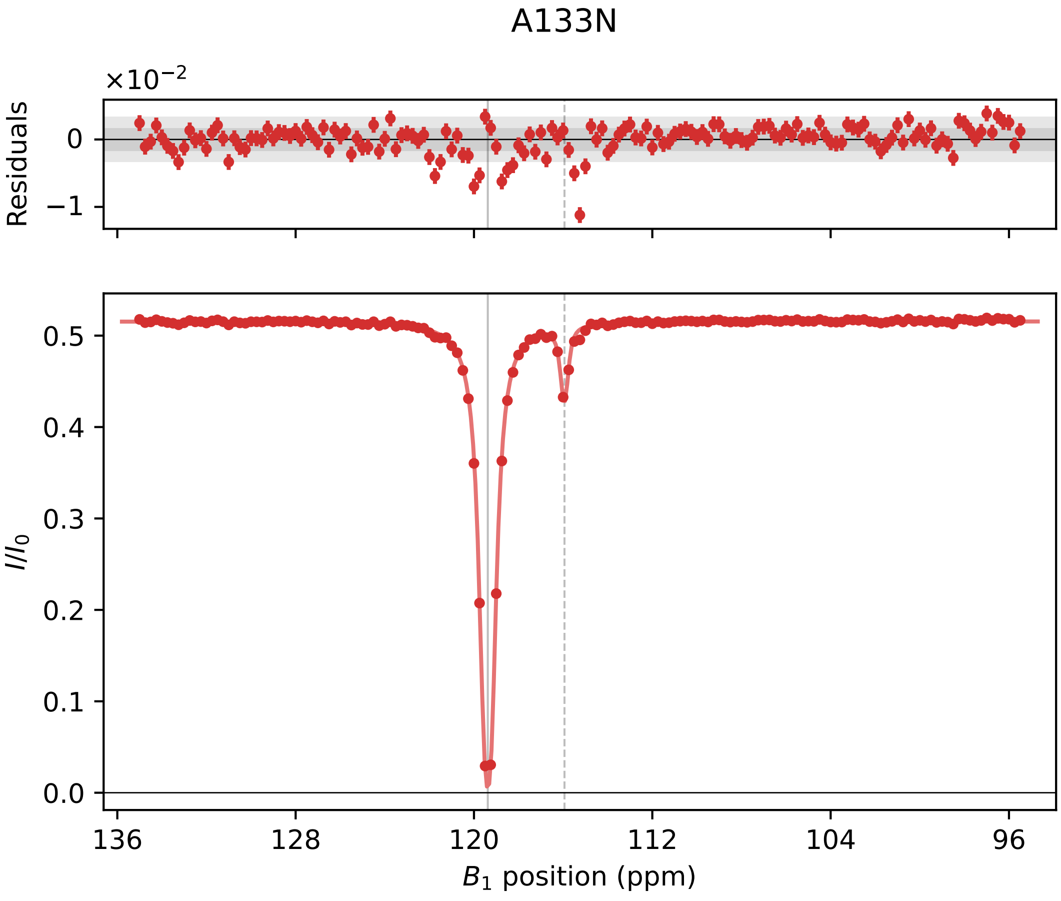

A133N
×10⁻²
Residuals
0
−1
I/I₀
0.5
0.4
0.3
0.2
0.1
0.0
136
128
120
112
104
96
B₁ position (ppm)

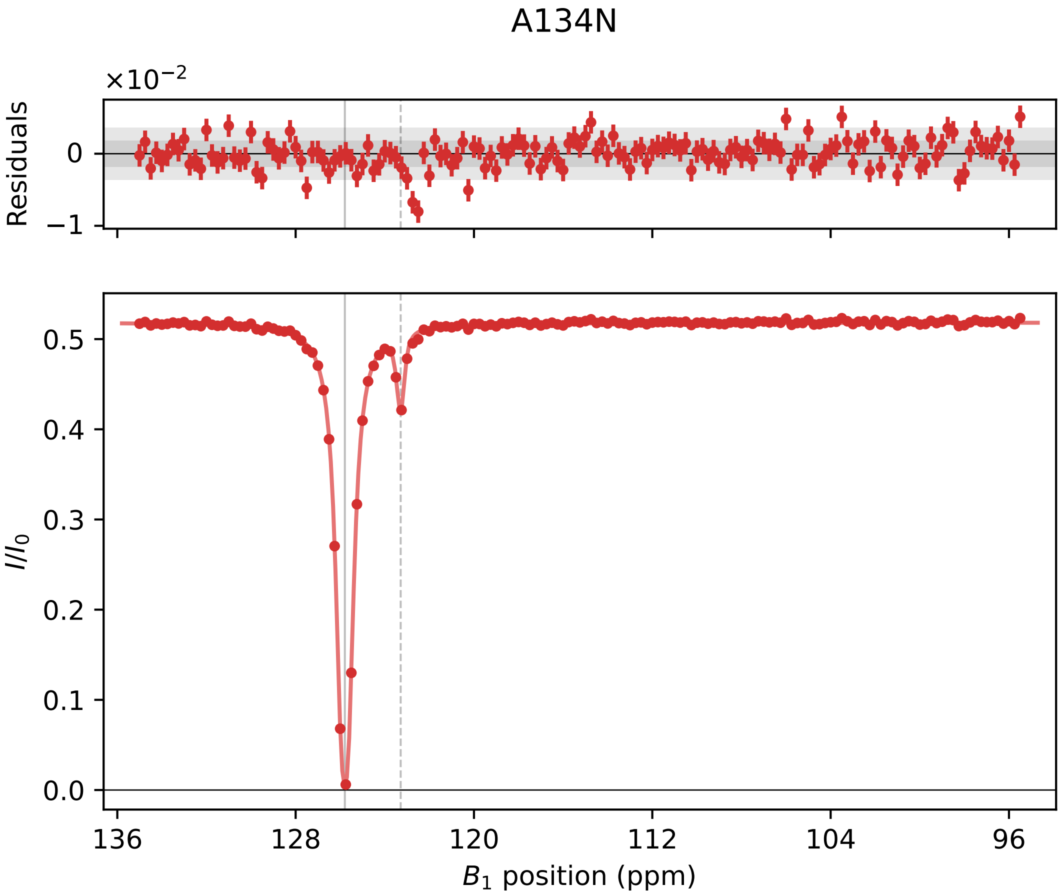

A134N
×10⁻²
Residuals
0
−1
I/I0
0.5
0.4
0.3
0.2
0.1
0.0
136
128
120
112
104
96
B1 position (ppm)

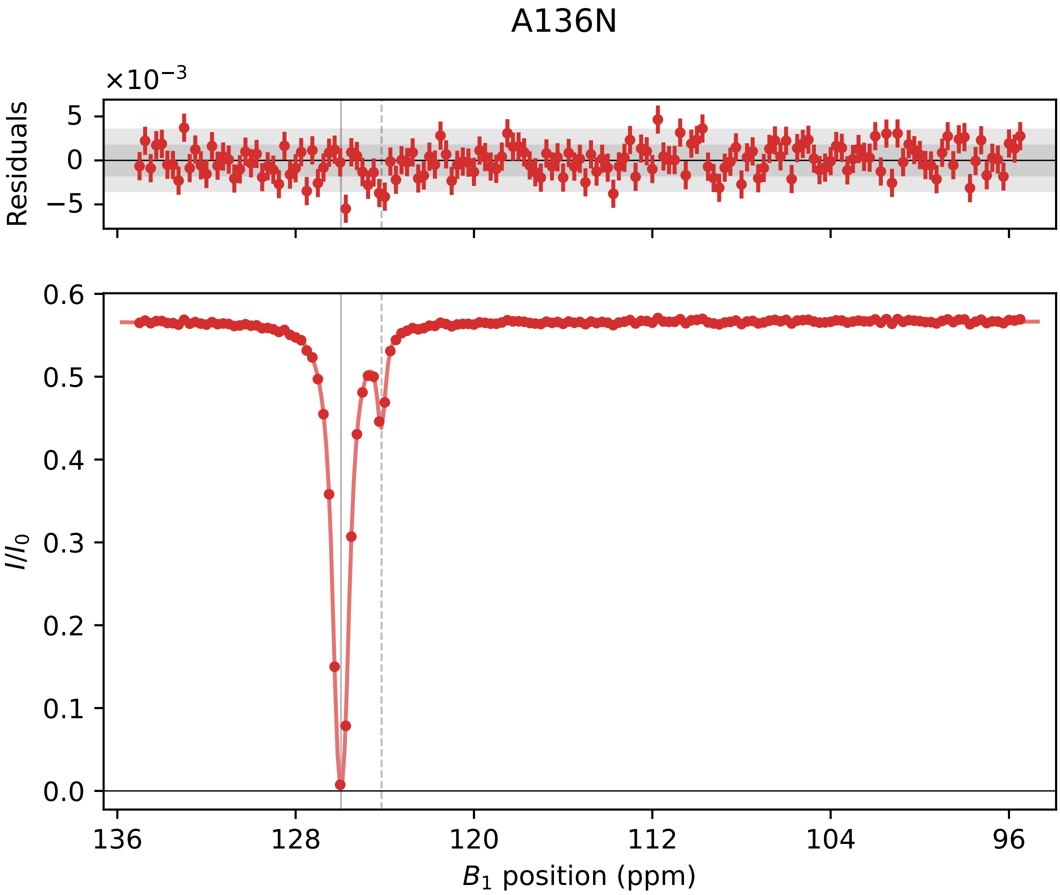

A136N
×10⁻³
Residuals
5
0
−5
0.6
0.5
0.4
0.3
0.2
0.1
0.0
$I/I_0$
136
128
120
112
104
96
$B_1$ position (ppm)

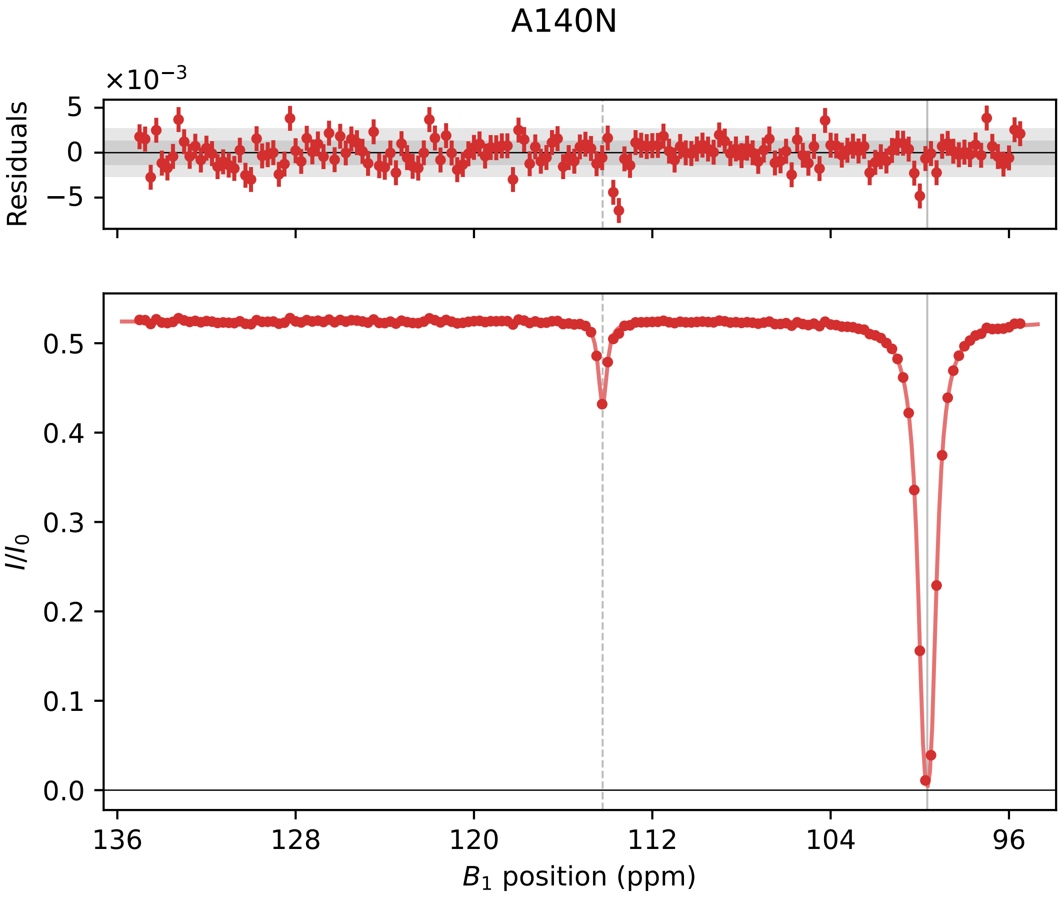

A140N
×10⁻³
Residuals
5
0
−5
0.5
0.4
0.3
0.2
0.1
0.0
I/I₀
136
128
120
112
104
96
B₁ position (ppm)

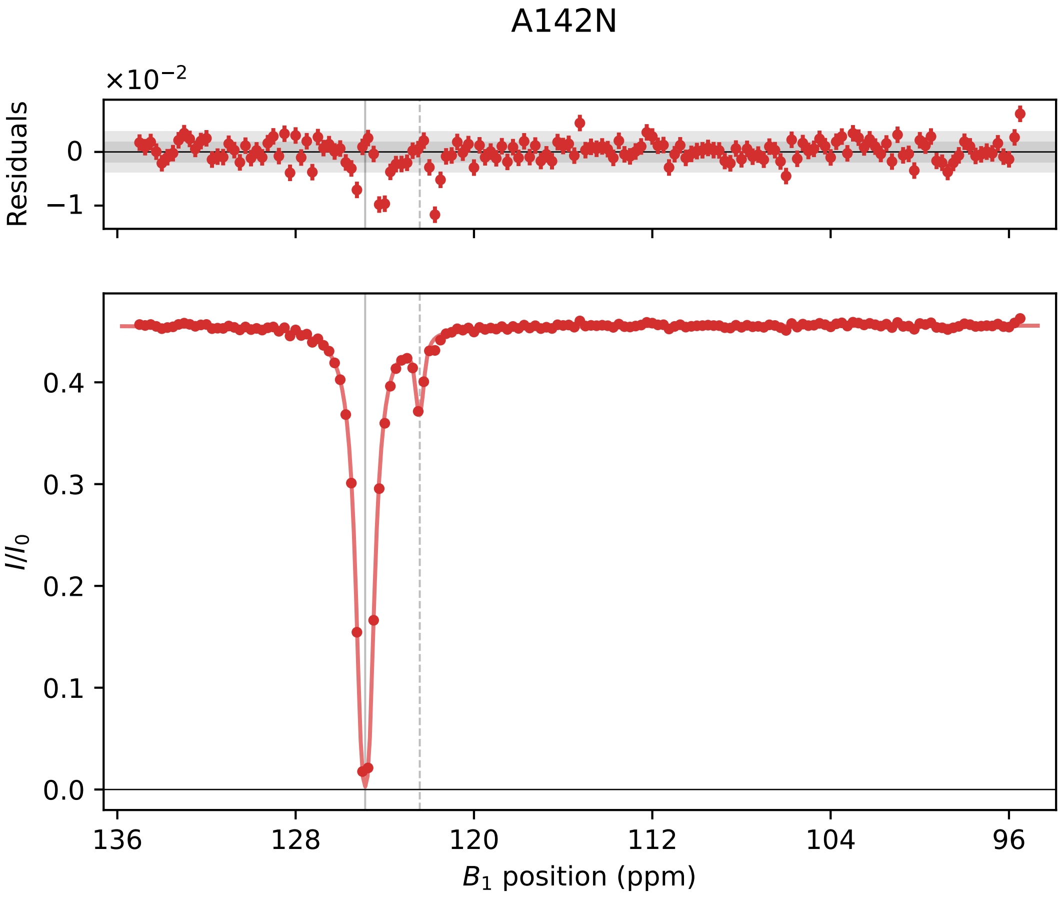

A142N
×10^−2
Residuals
0
−1
I/I0
0.4
0.3
0.2
0.1
0.0
136
128
120
112
104
96
B1 position (ppm)

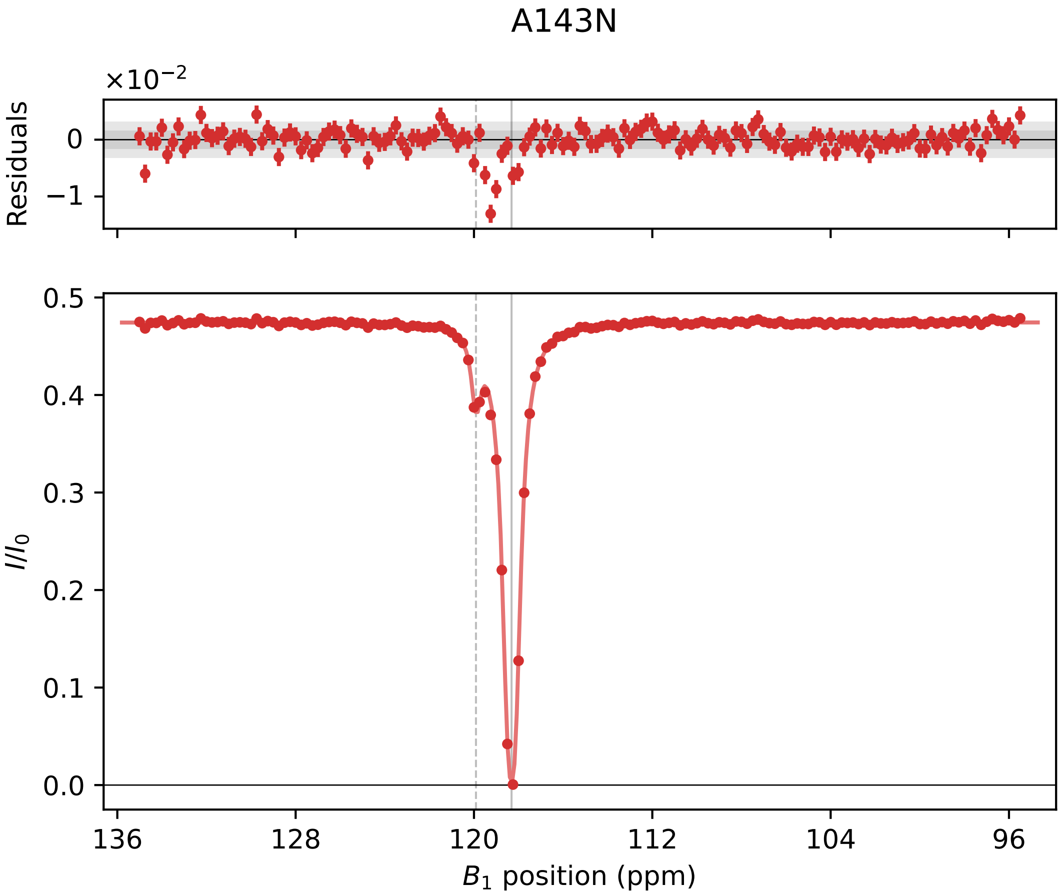

A143N
$\times10^{-2}$
Residuals
0
−1
$I/I_0$
0.5
0.4
0.3
0.2
0.1
0.0
136
128
120
112
104
96
$B_1$ position (ppm)

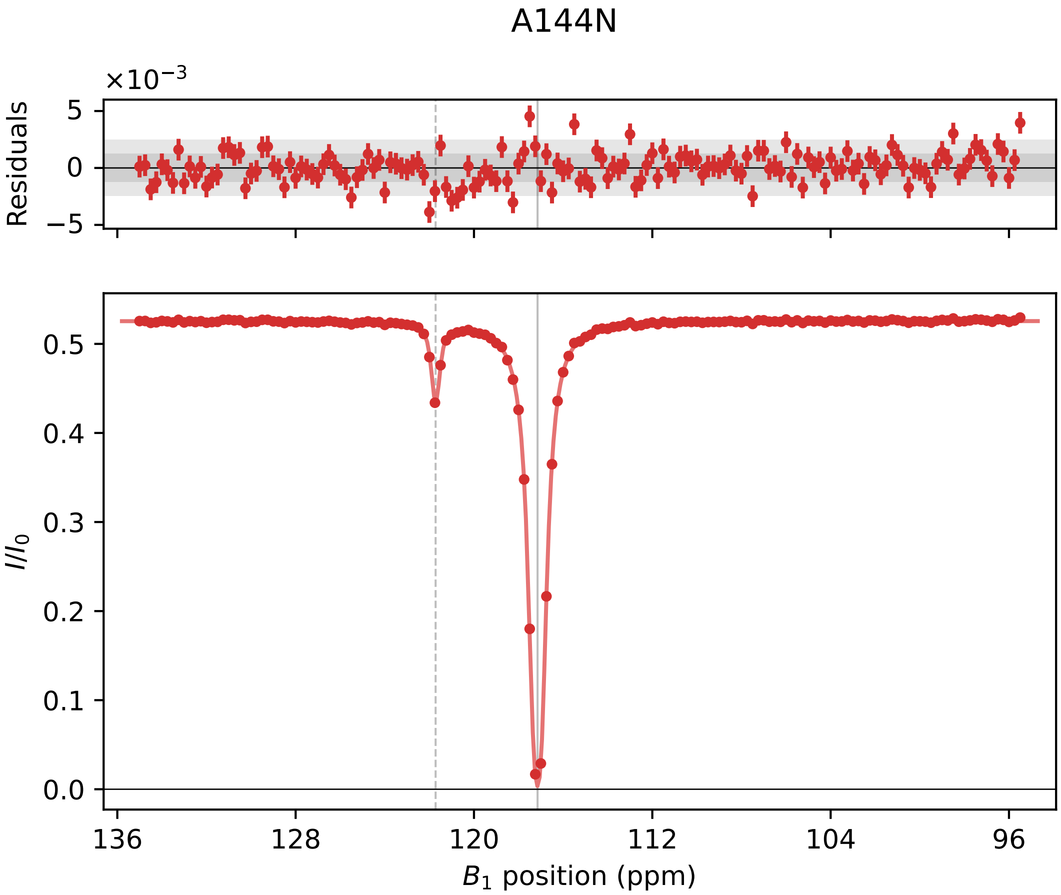

A144N
×10⁻³
Residuals
5
0
−5
0.5
0.4
0.3
0.2
0.1
0.0
$I/I_0$
136
128
120
112
104
96
$B_1$ position (ppm)

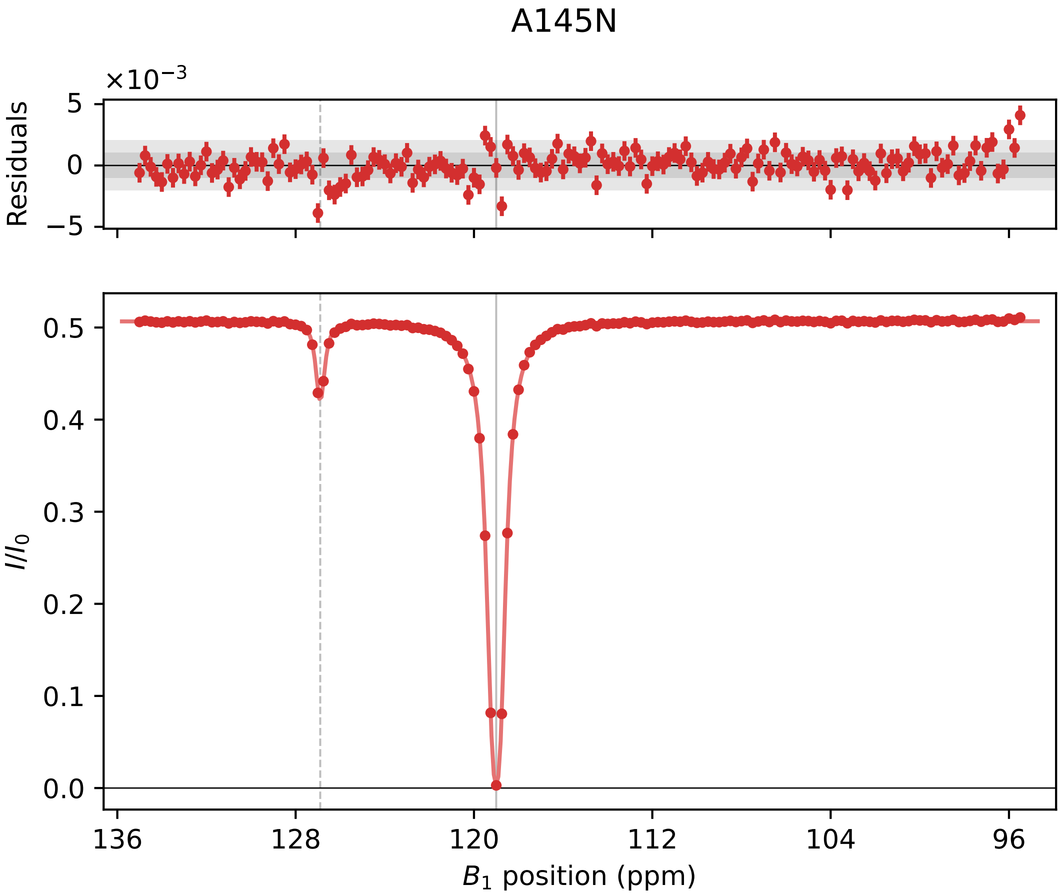
A145N
×10⁻³
Residuals
5
0
−5
I/I₀
0.5
0.4
0.3
0.2
0.1
0.0
136
128
120
112
104
96
B₁ position (ppm)

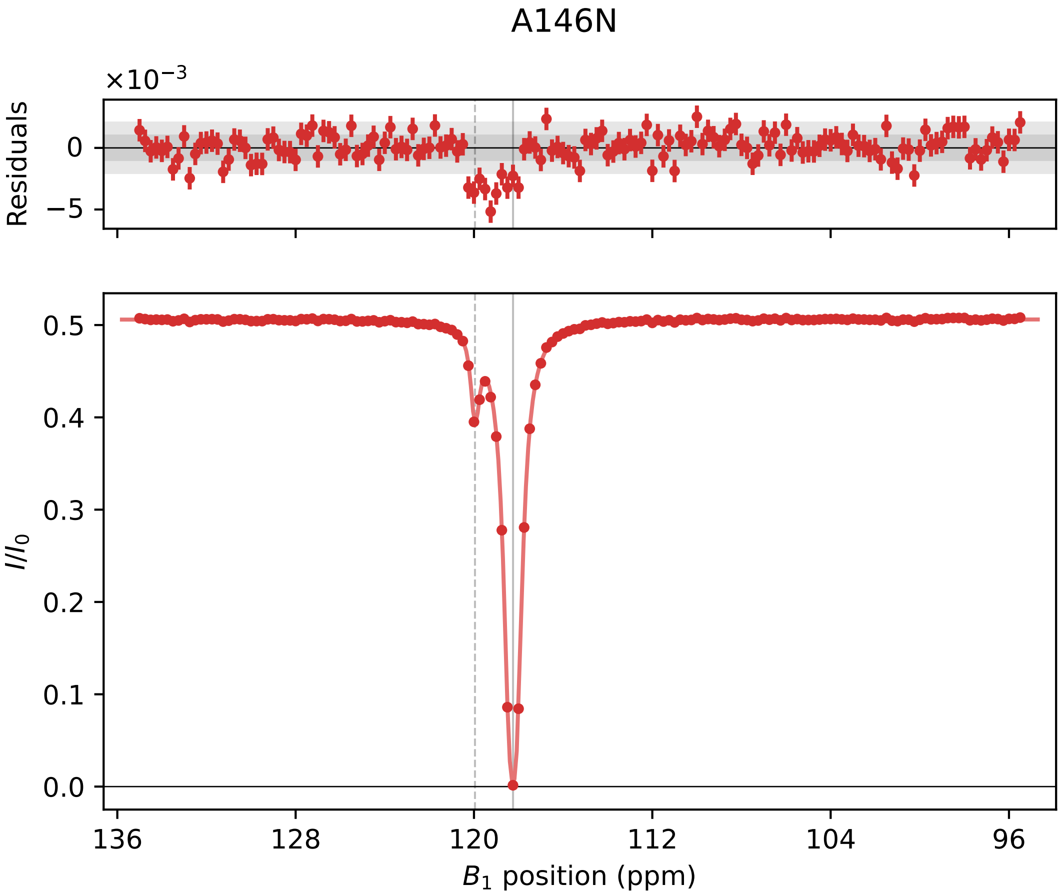

A146N
×10^−3
Residuals
0
−5
0.5
0.4
0.3
0.2
0.1
0.0
I/I0
136
128
120
112
104
96
B1 position (ppm)

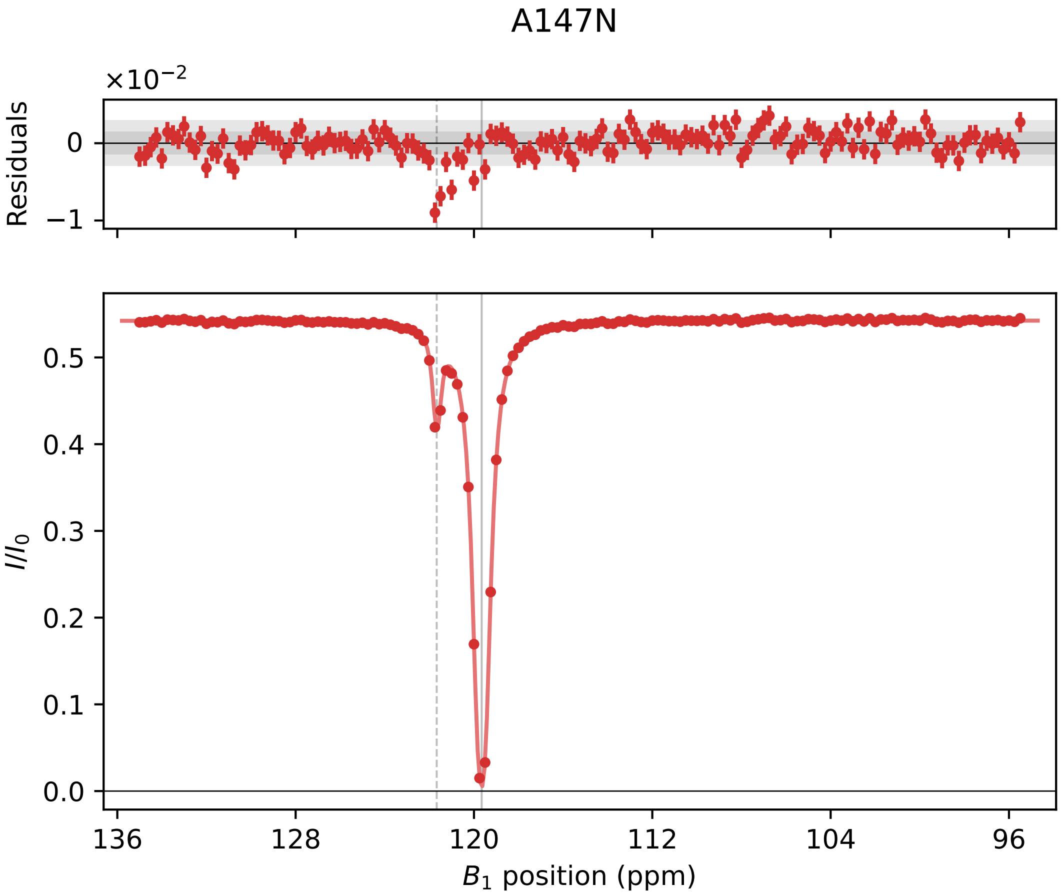

A147N
×10^−2
Residuals
0
−1
I/I_0
0.5
0.4
0.3
0.2
0.1
0.0
136
128
120
112
104
96
B_1 position (ppm)

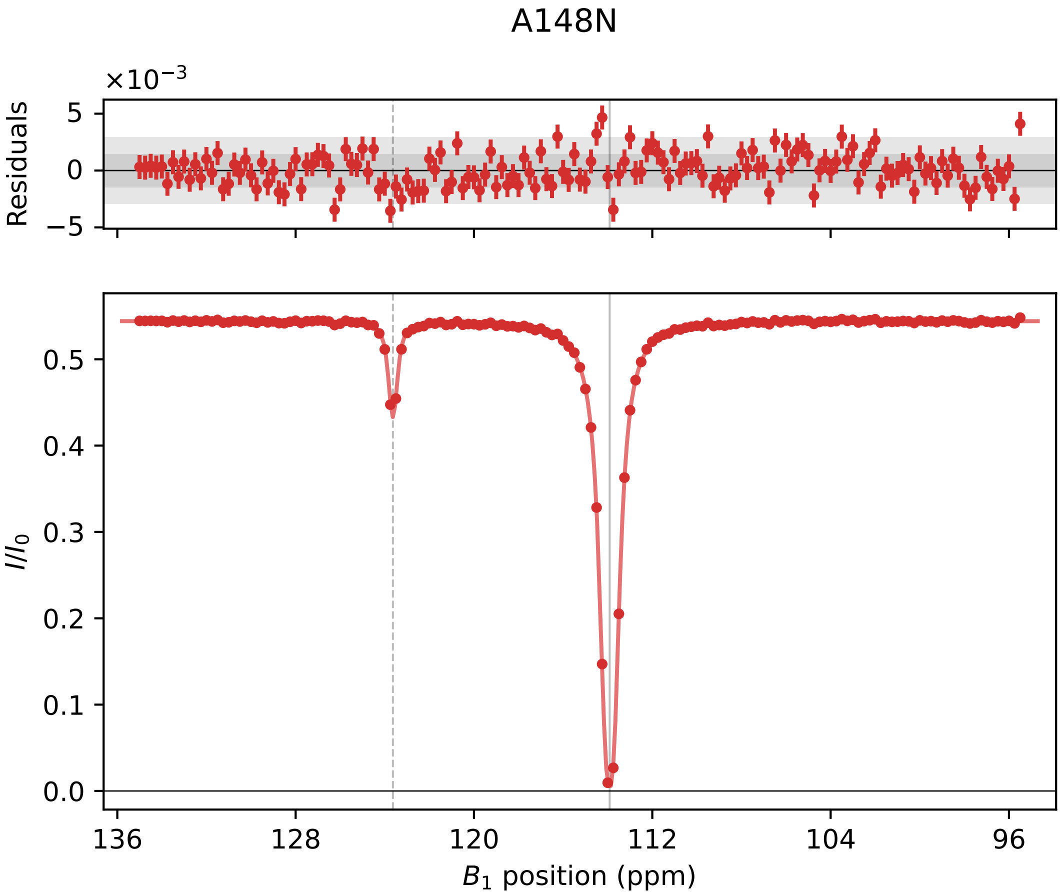
A148N
×10⁻³
Residuals
5
0
−5
I/I0
0.5
0.4
0.3
0.2
0.1
0.0
136
128
120
112
104
96
B1 position (ppm)

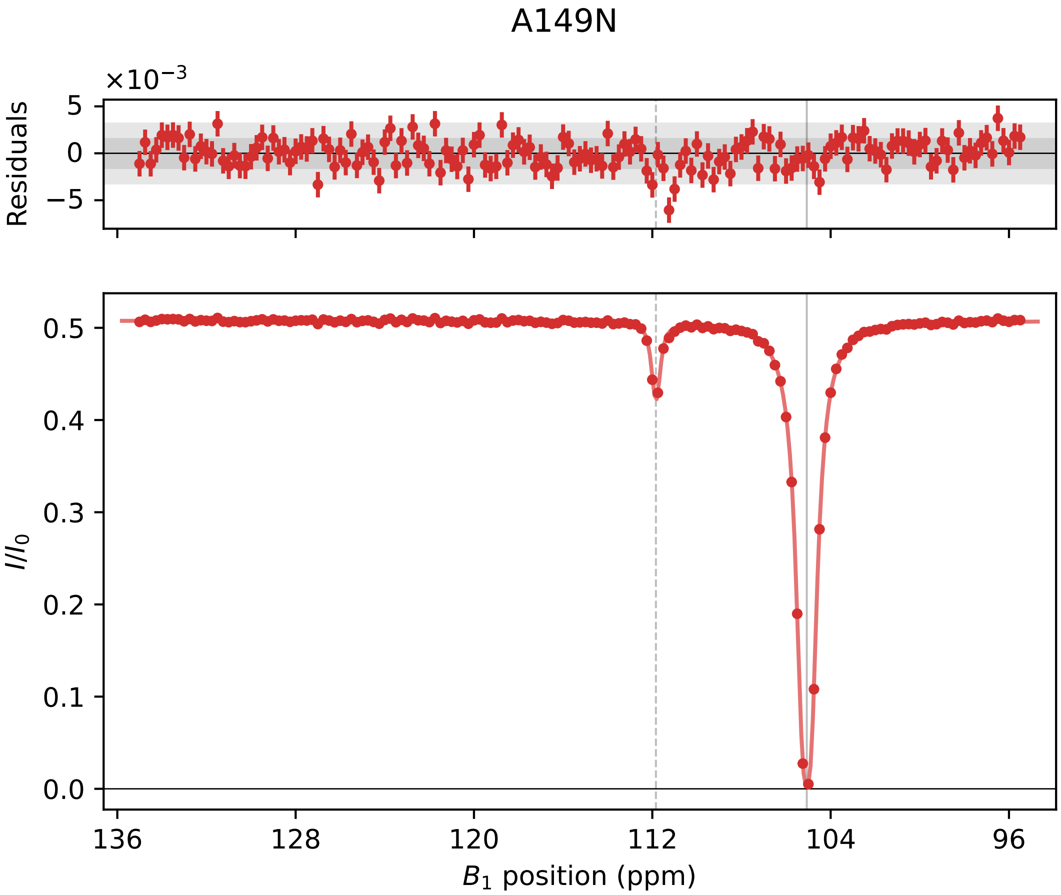
A149N
×10^−3
Residuals
5
0
−5
0.5
0.4
0.3
0.2
0.1
0.0
I/I_0
136
128
120
112
104
96
B_1 position (ppm)

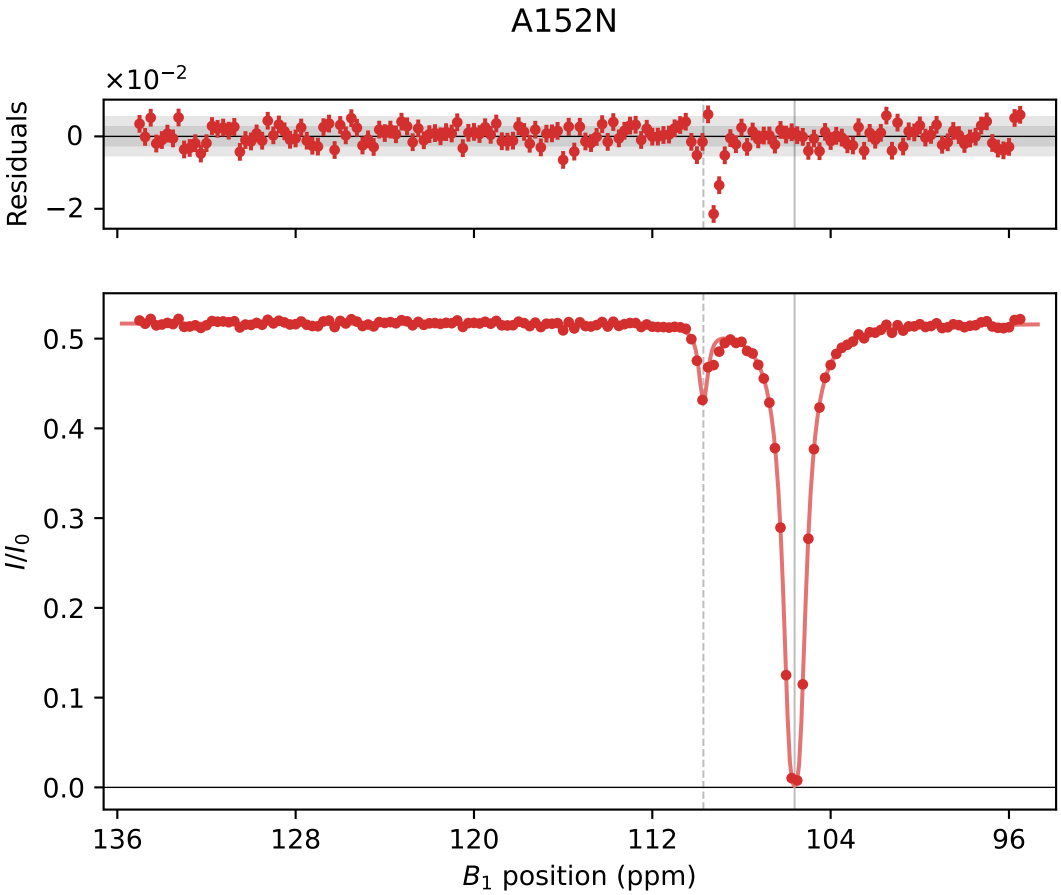
A152N
×10⁻²
Residuals
0
−2
I/I₀
0.5
0.4
0.3
0.2
0.1
0.0
136
128
120
112
104
96
B₁ position (ppm)

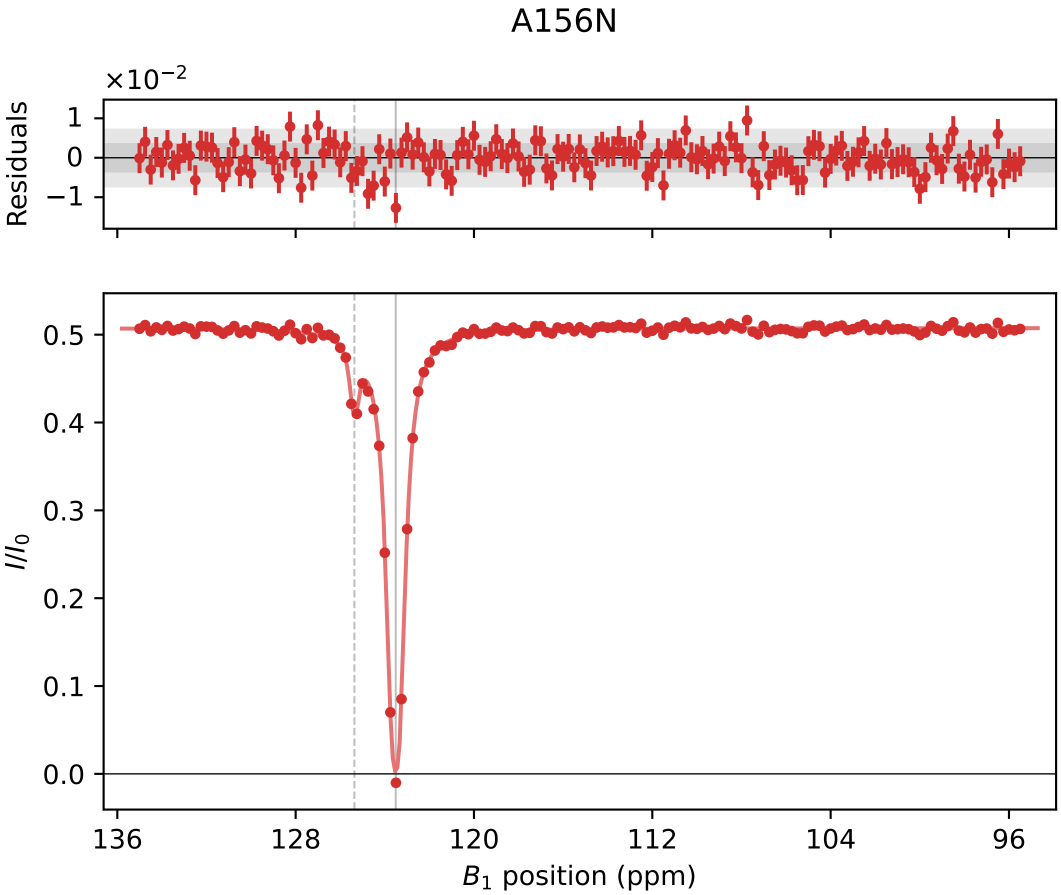

A156N
×10^−2
Residuals
1
0
−1
0.5
0.4
0.3
0.2
0.1
0.0
I/I0
136
128
120
112
104
96
B1 position (ppm)

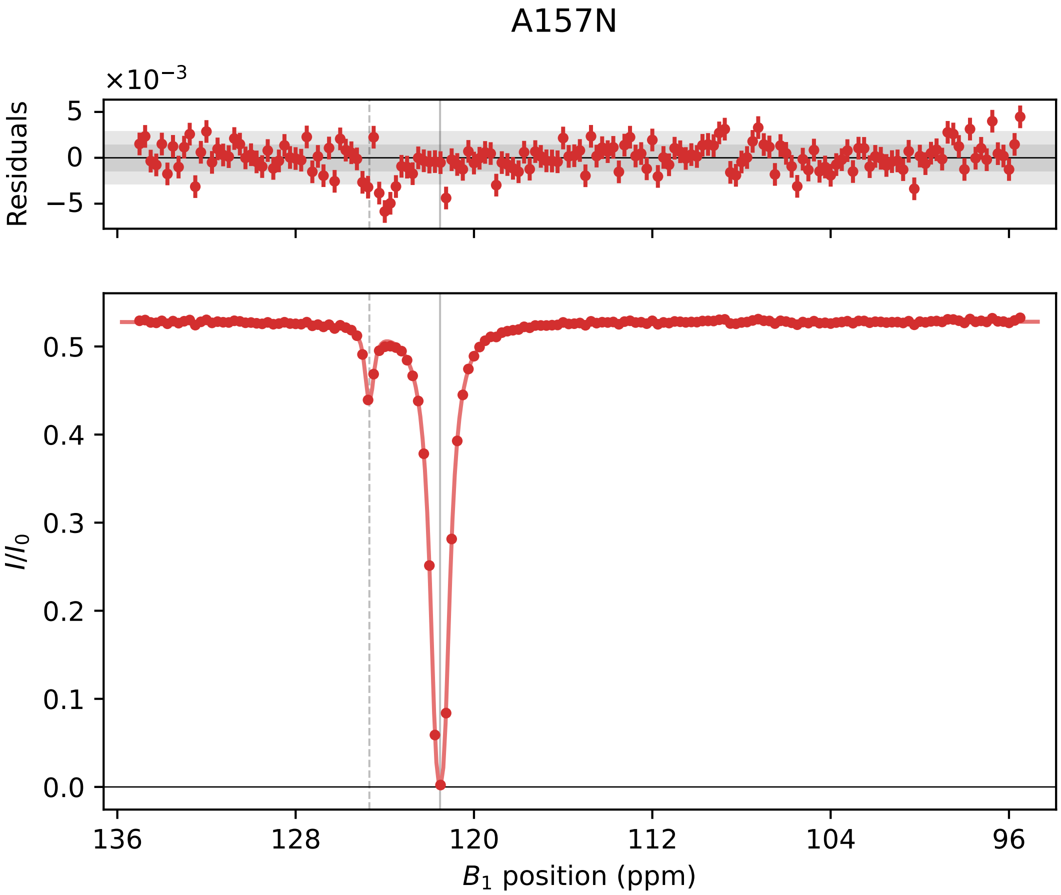
A157N
×10⁻³
Residuals
5
0
−5
I/I₀
0.5
0.4
0.3
0.2
0.1
0.0
136
128
120
112
104
96
B₁ position (ppm)

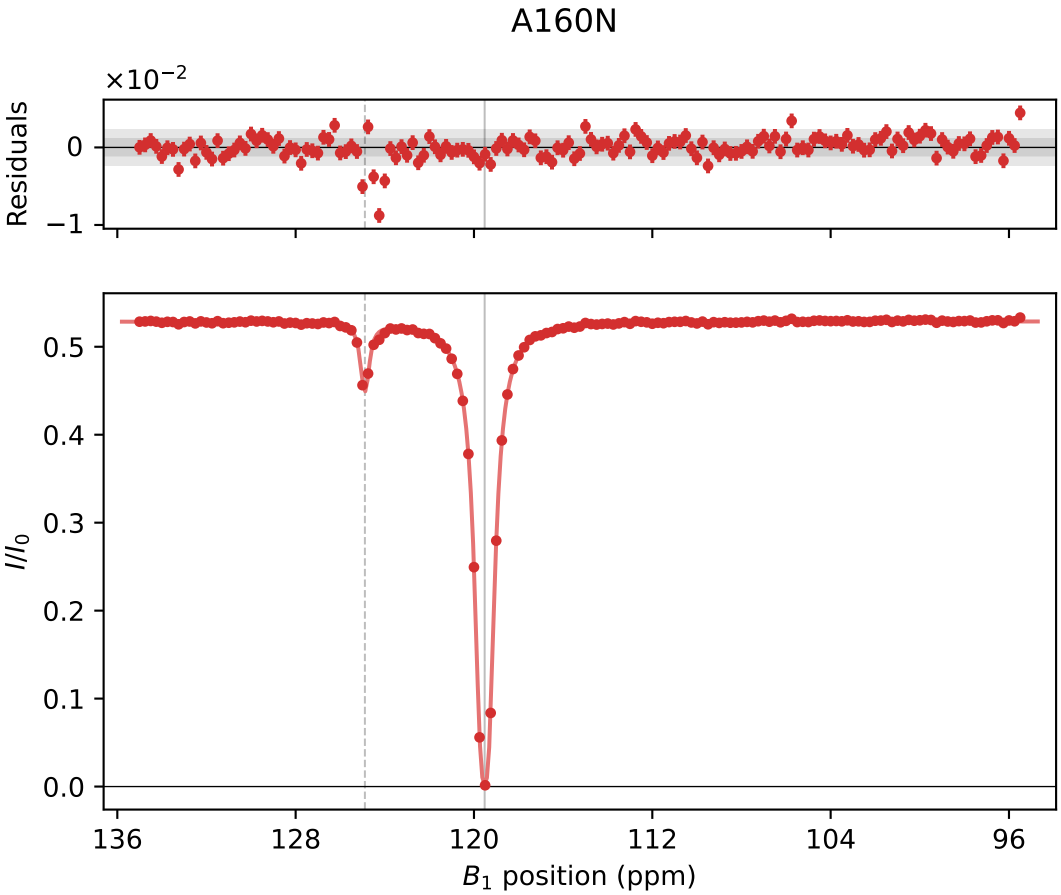

A160N
×10⁻²
Residuals
0
−1
I/I0
0.5
0.4
0.3
0.2
0.1
0.0
136
128
120
112
104
96
B1 position (ppm)

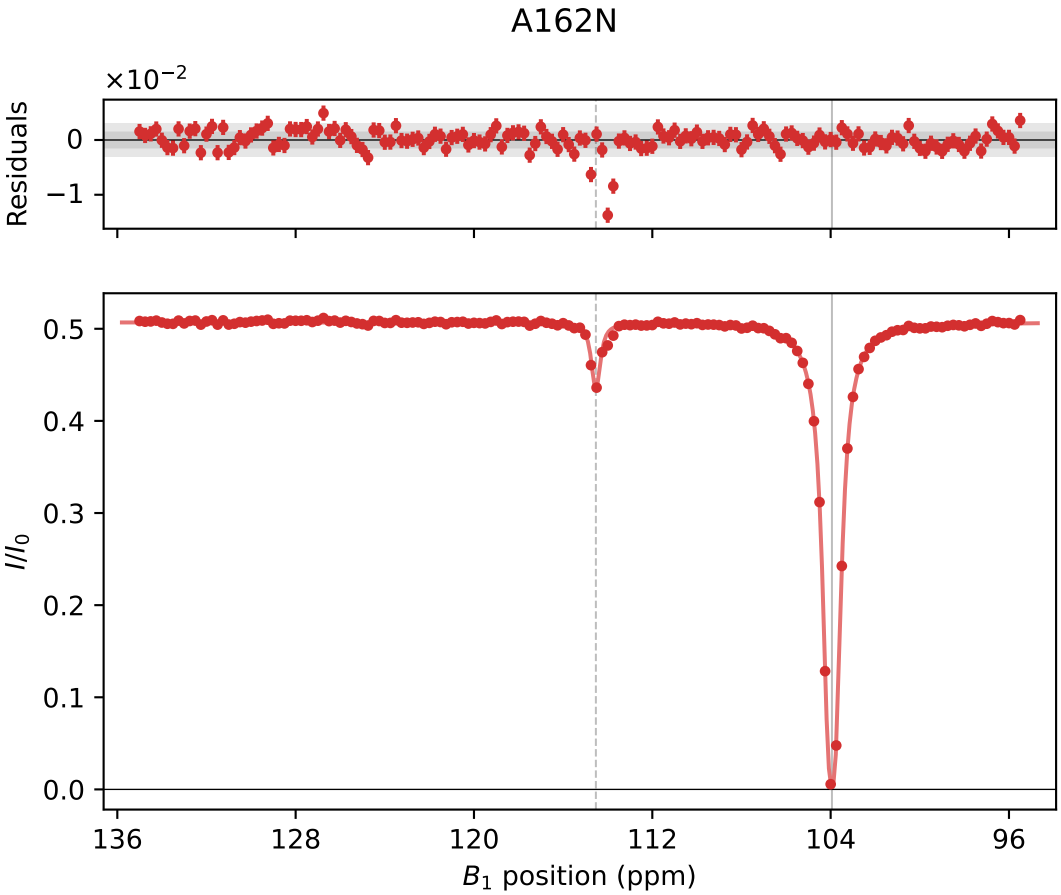

A162N
×10−2
Residuals
0
−1
I/I0
0.5
0.4
0.3
0.2
0.1
0.0
136
128
120
112
104
96
B1 position (ppm)

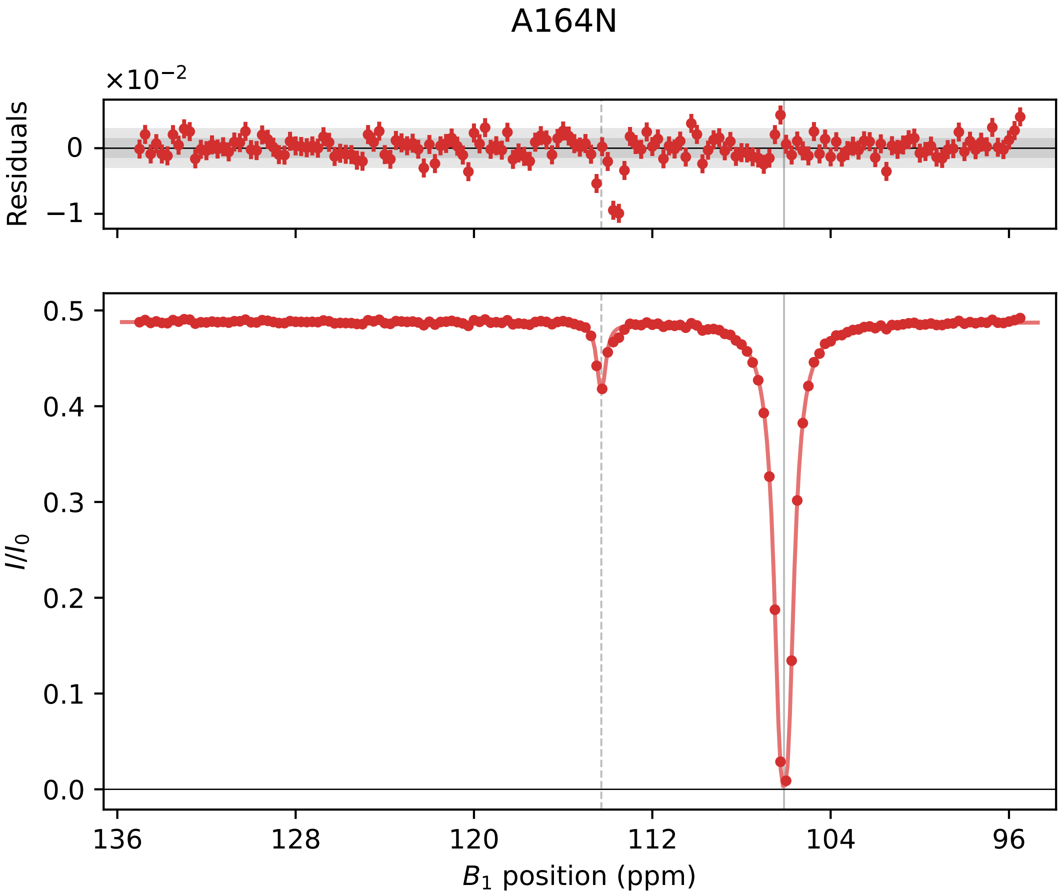

A164N
×10^−2
Residuals
0
−1
I/I_0
0.5
0.4
0.3
0.2
0.1
0.0
136
128
120
112
104
96
B_1 position (ppm)

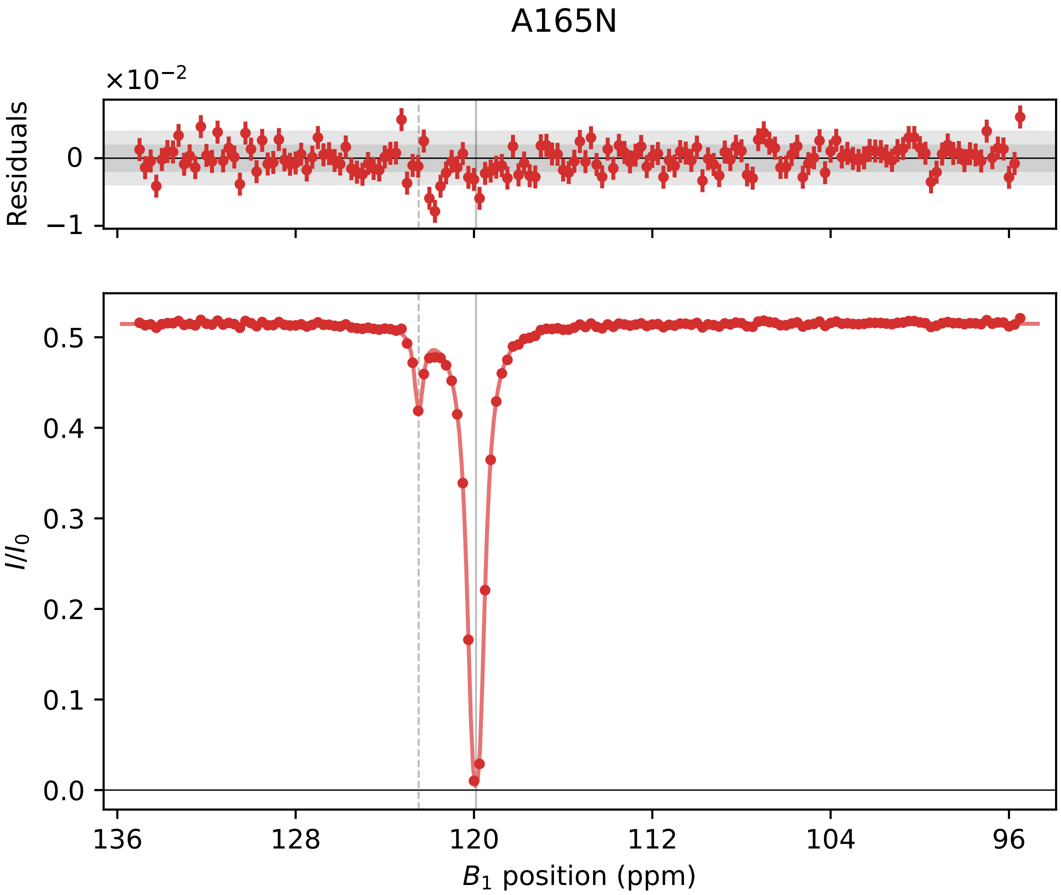

A165N
×10^−2
Residuals
0
−1
I/I_0
0.5
0.4
0.3
0.2
0.1
0.0
136
128
120
112
104
96
B_1 position (ppm)

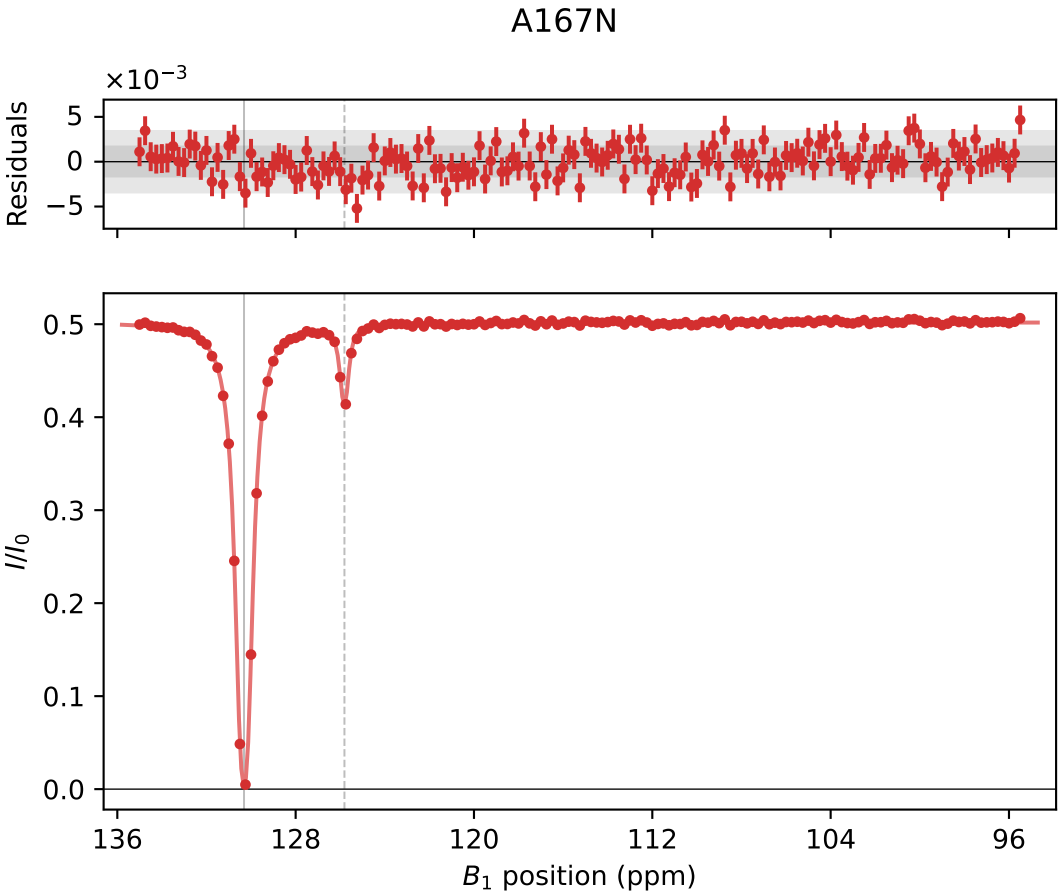
A167N
×10^−3
Residuals
5
0
−5
0.5
0.4
0.3
0.2
0.1
0.0
I/I0
136
128
120
112
104
96
B1 position (ppm)

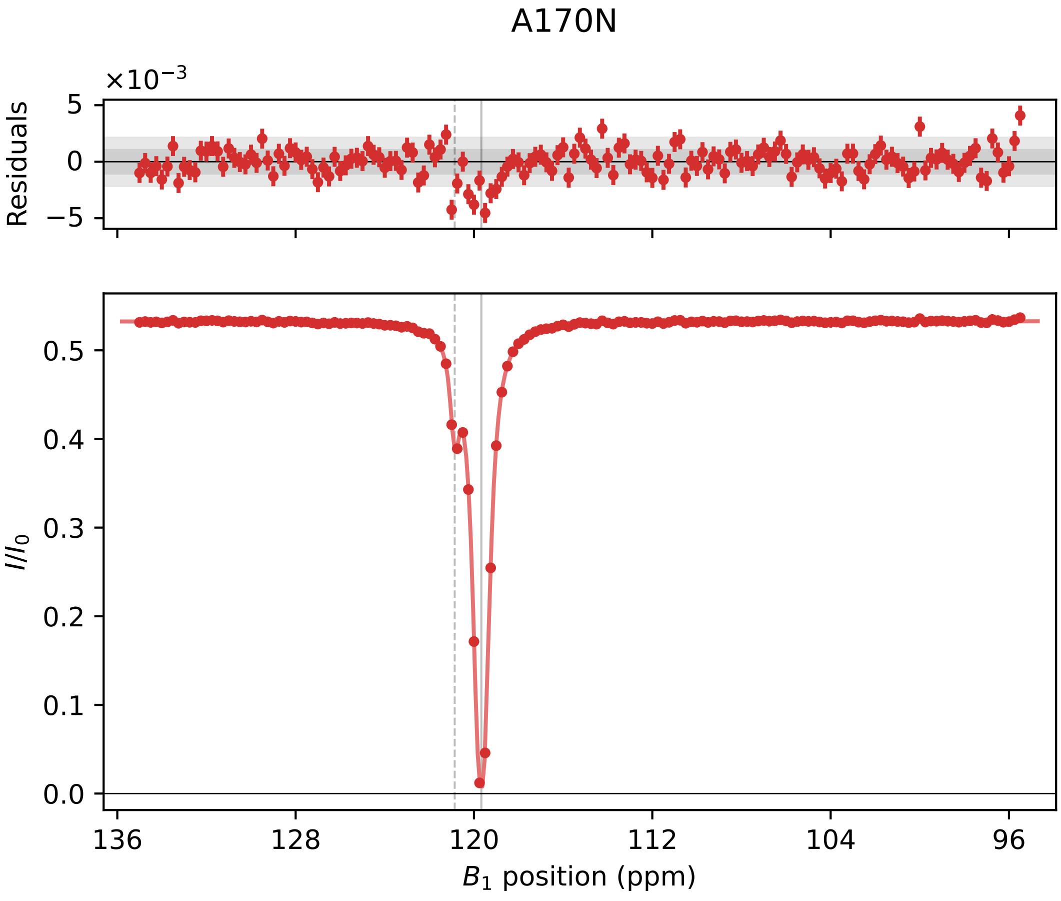

A170N
×10⁻³
Residuals
5
0
−5
I/I0
0.5
0.4
0.3
0.2
0.1
0.0
136
128
120
112
104
96
B1 position (ppm)

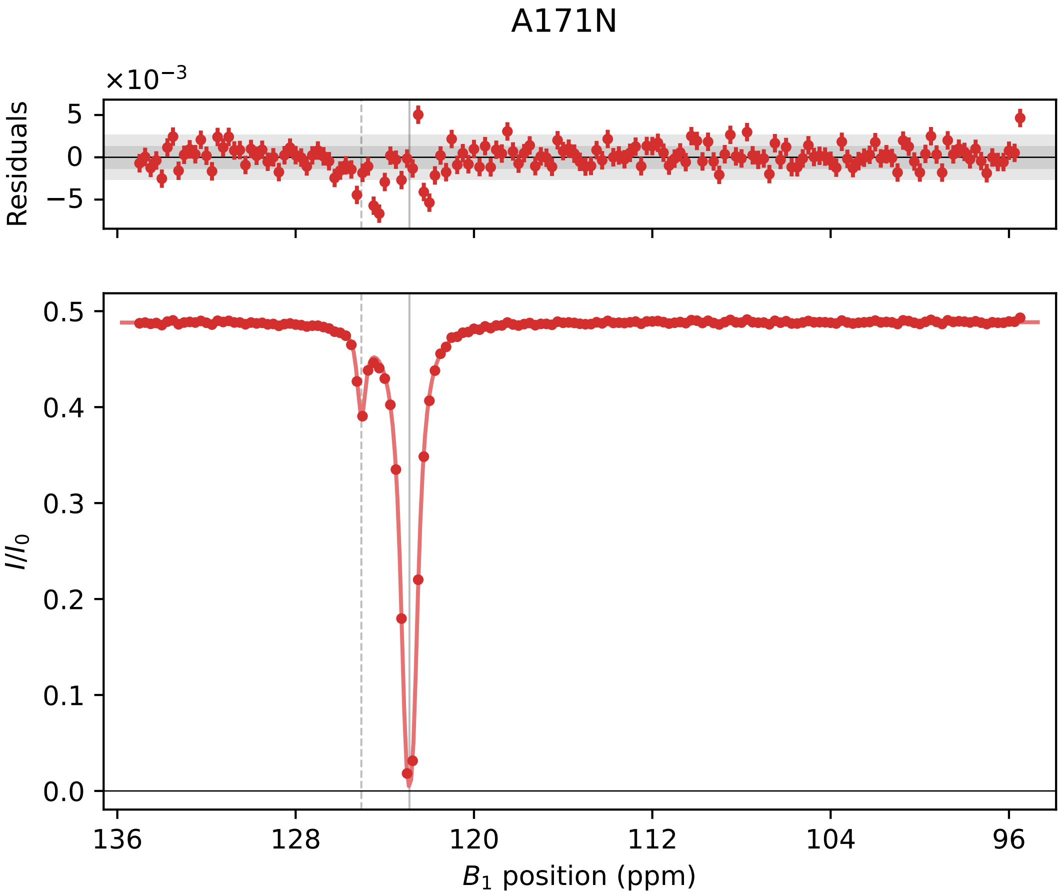

A171N
×10⁻³
Residuals
5
0
−5
0.5
0.4
0.3
0.2
0.1
0.0
I/I0
136
128
120
112
104
96
B1 position (ppm)

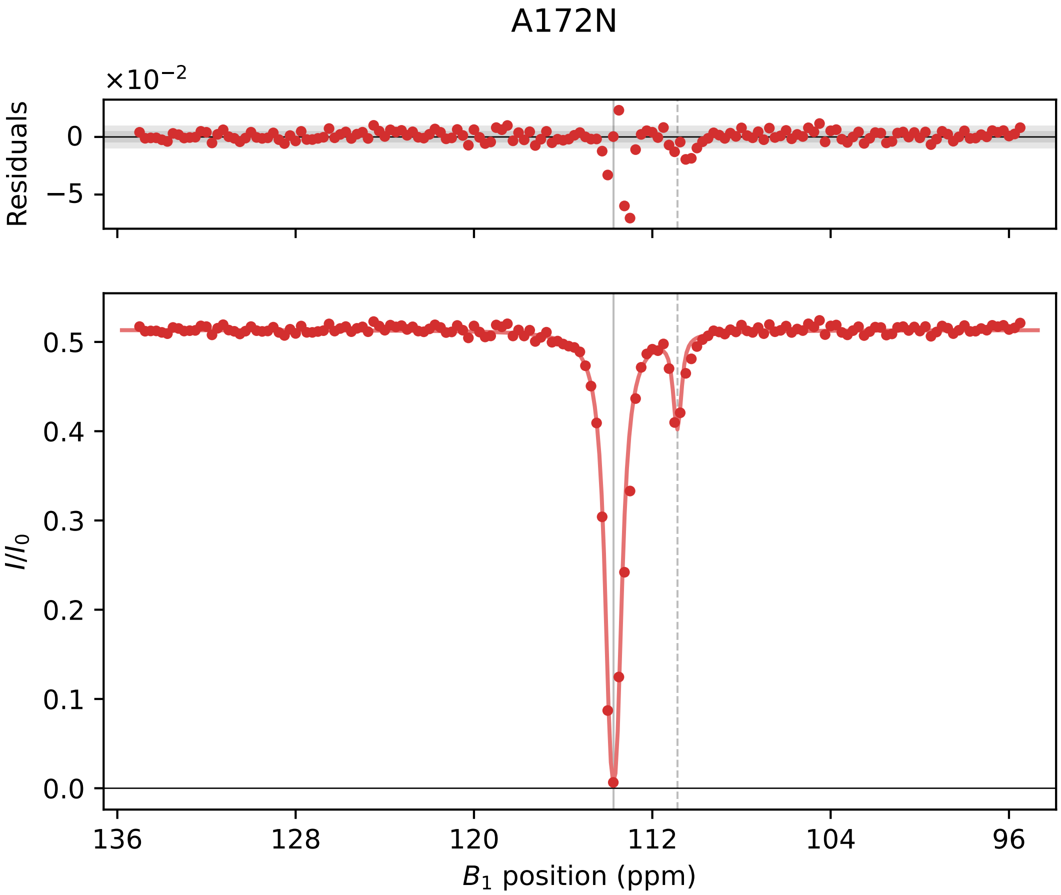
A172N
×10⁻²
Residuals
0
−5
I/I0
0.5
0.4
0.3
0.2
0.1
0.0
136
128
120
112
104
96
B1 position (ppm)

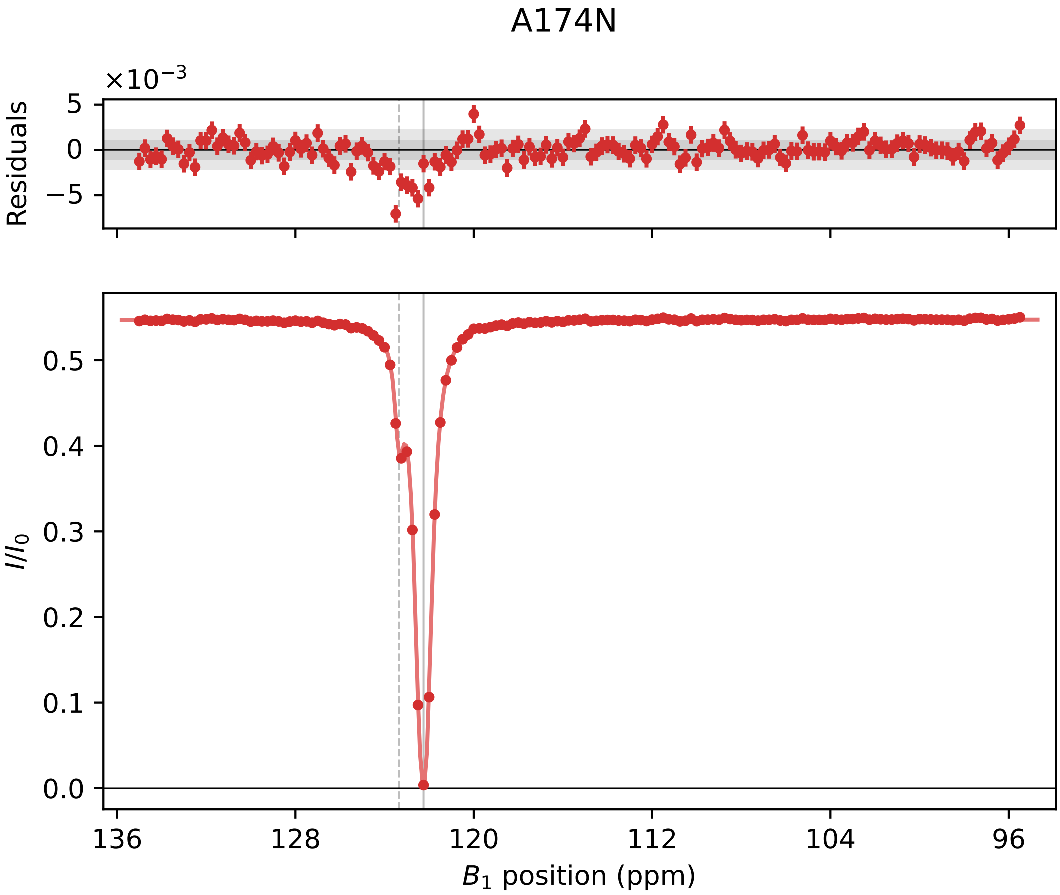

A174N
×10⁻³
Residuals
5
0
−5
I/I0
0.5
0.4
0.3
0.2
0.1
0.0
136
128
120
112
104
96
B1 position (ppm)

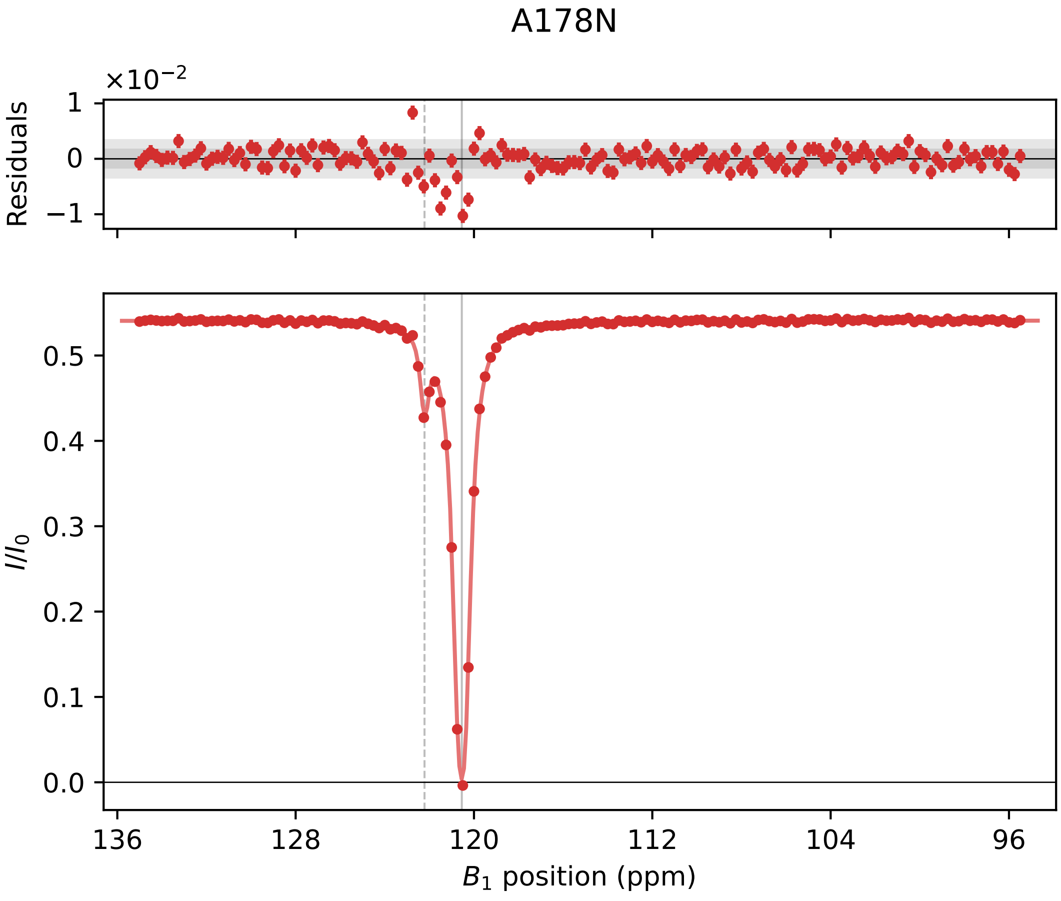

A178N
×10−2
Residuals
1
0
−1
I/I0
0.5
0.4
0.3
0.2
0.1
0.0
136
128
120
112
104
96
B1 position (ppm)

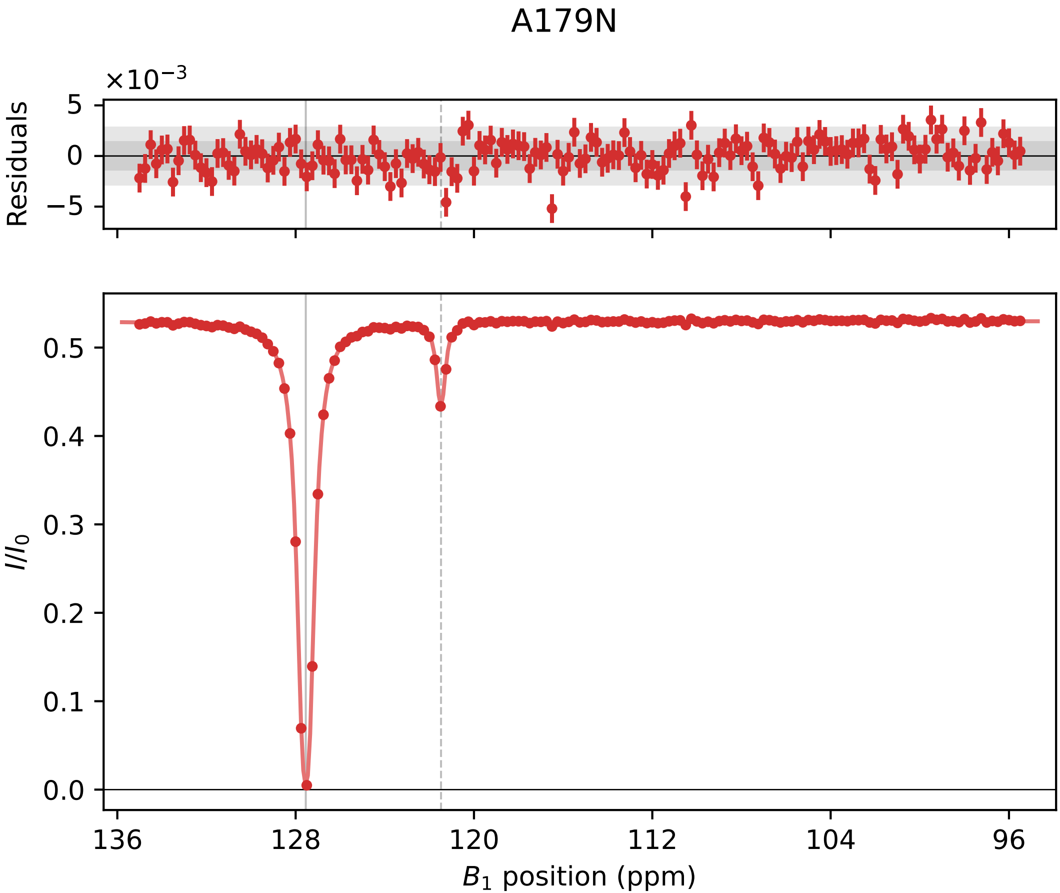
A179N
×10⁻³
Residuals
5
0
−5
I/I₀
0.5
0.4
0.3
0.2
0.1
0.0
136
128
120
112
104
96
B₁ position (ppm)

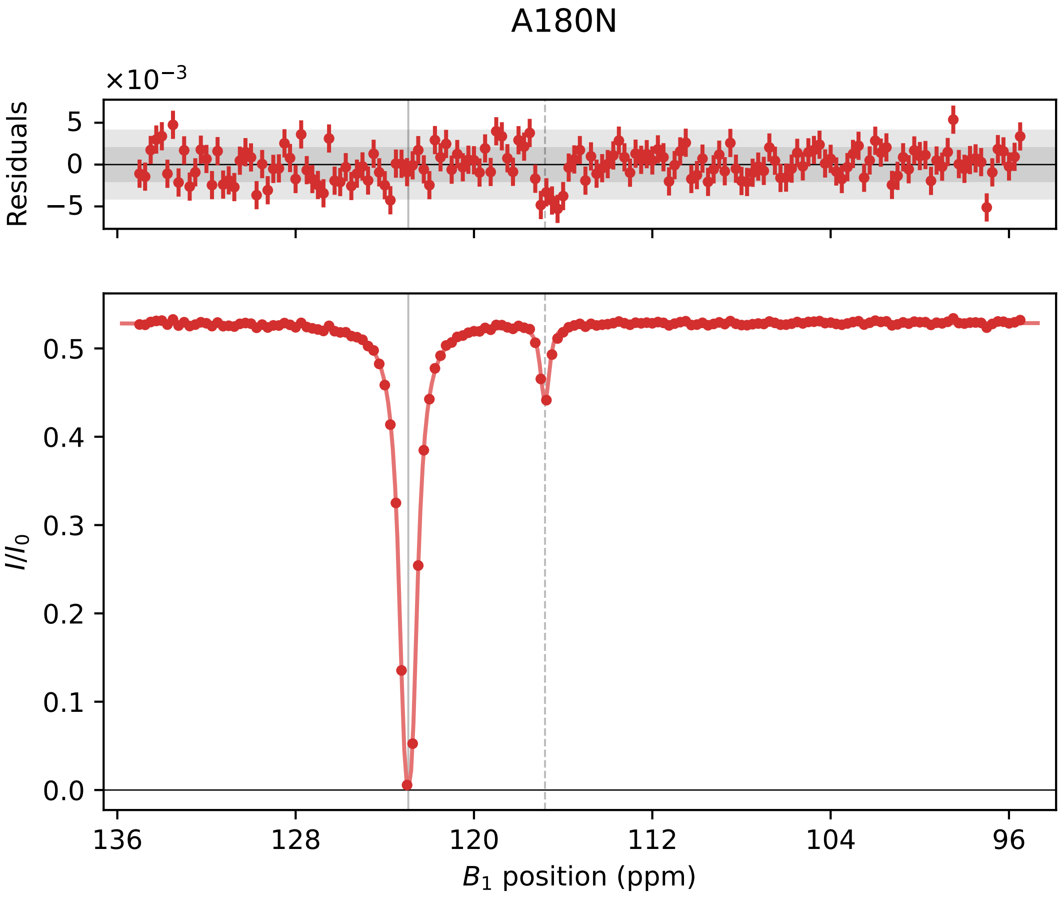

A180N
×10^−3
Residuals
5
0
−5
0.5
0.4
0.3
0.2
0.1
0.0
I/I_0
136
128
120
112
104
96
B_1 position (ppm)

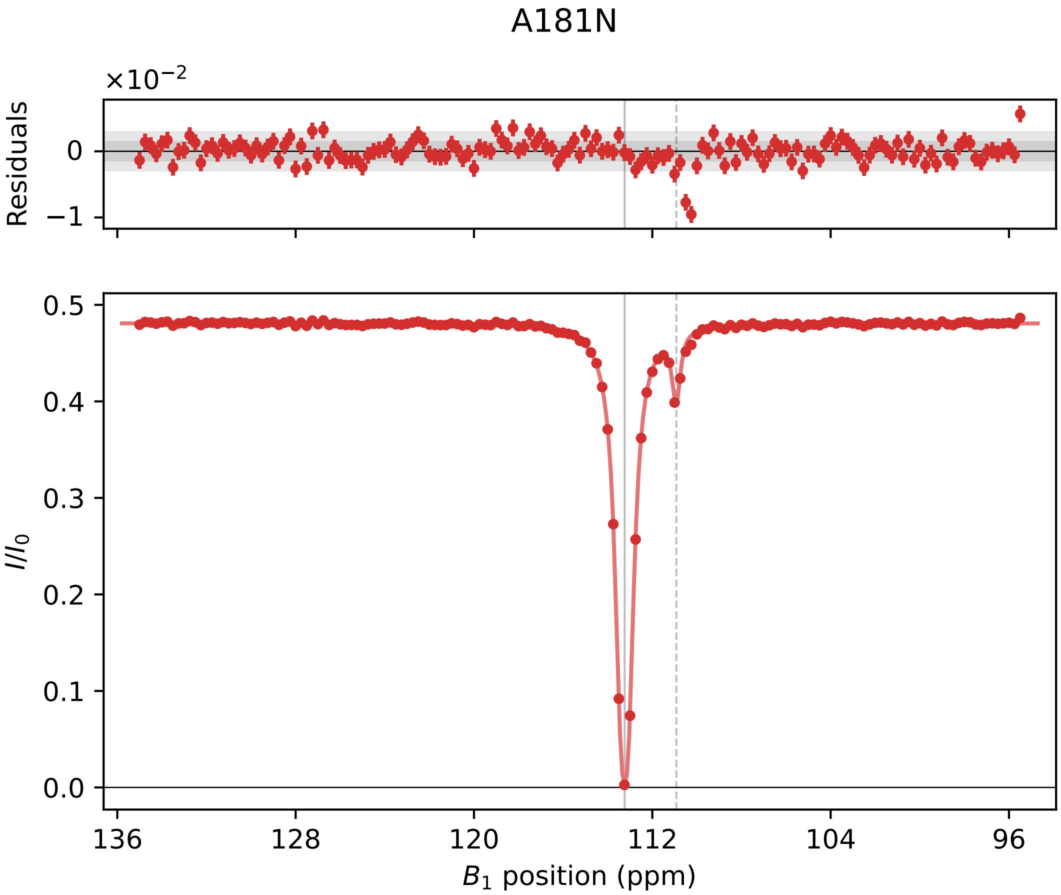

A181N
×10⁻²
Residuals
0
−1
I/I0
0.5
0.4
0.3
0.2
0.1
0.0
136
128
120
112
104
96
B1 position (ppm)

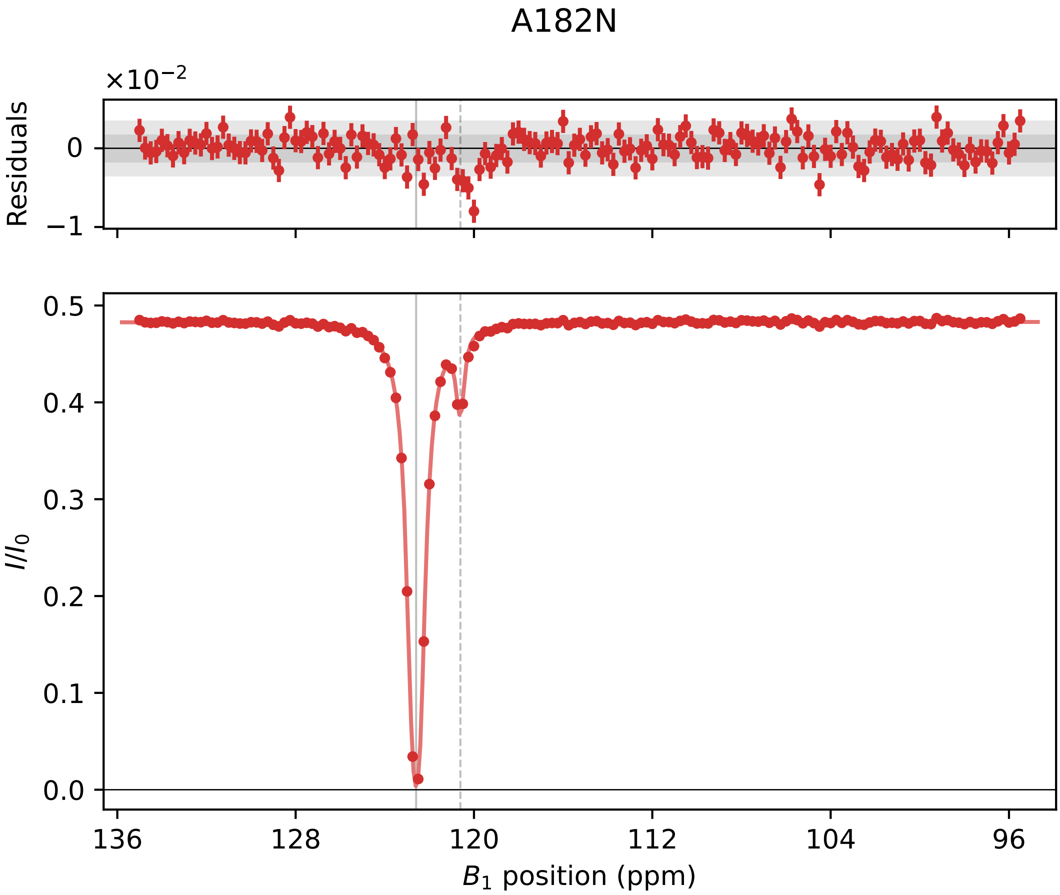

A182N
×10−2
Residuals
0
−1
I/I0
0.5
0.4
0.3
0.2
0.1
0.0
136
128
120
112
104
96
B1 position (ppm)

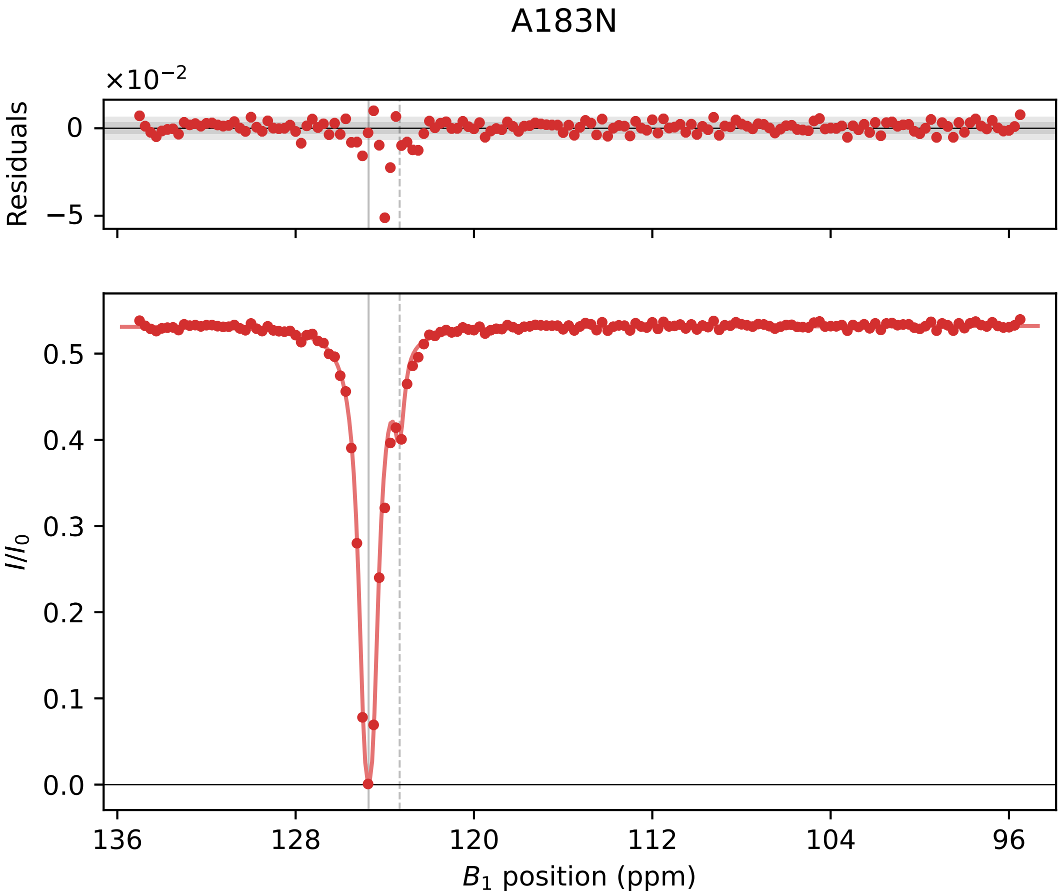

A183N
×10^−2
Residuals
0
−5
I/I0
0.0
0.1
0.2
0.3
0.4
0.5
136
128
120
112
104
96
B1 position (ppm)

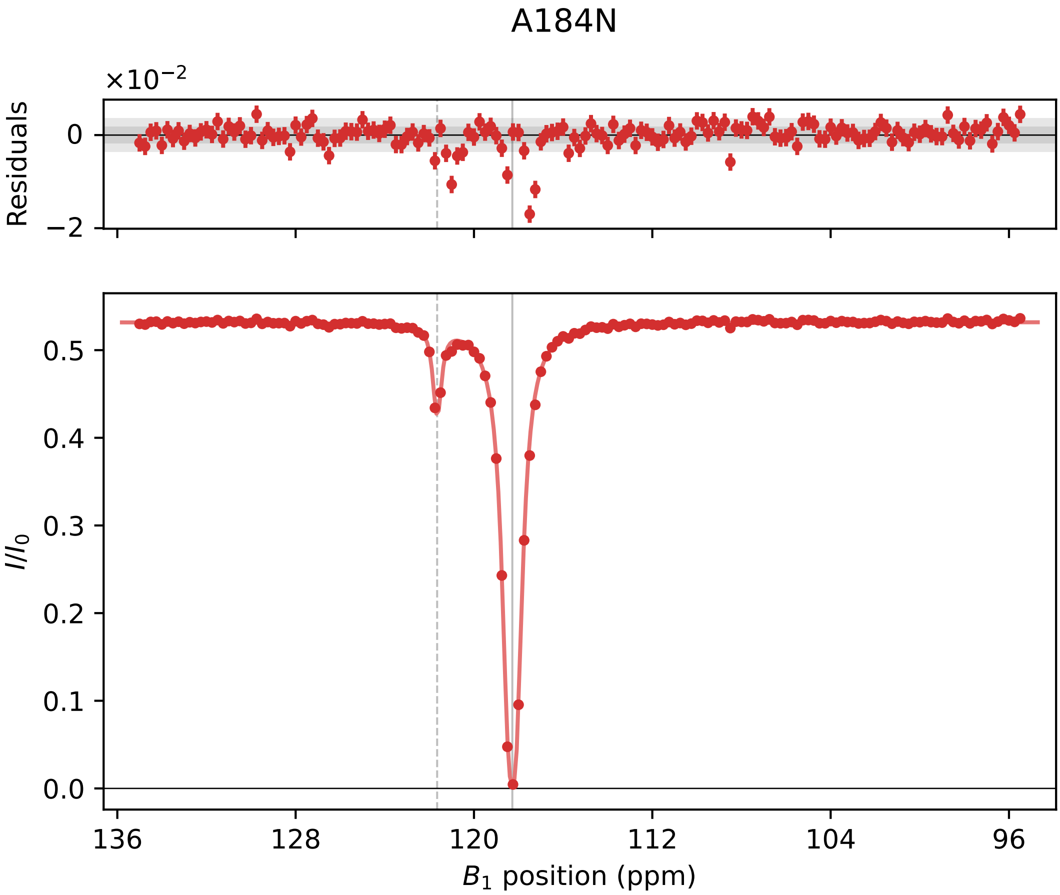
A184N
×10^−2
Residuals
0
−2
I/I0
0.5
0.4
0.3
0.2
0.1
0.0
136
128
120
112
104
96
B1 position (ppm)

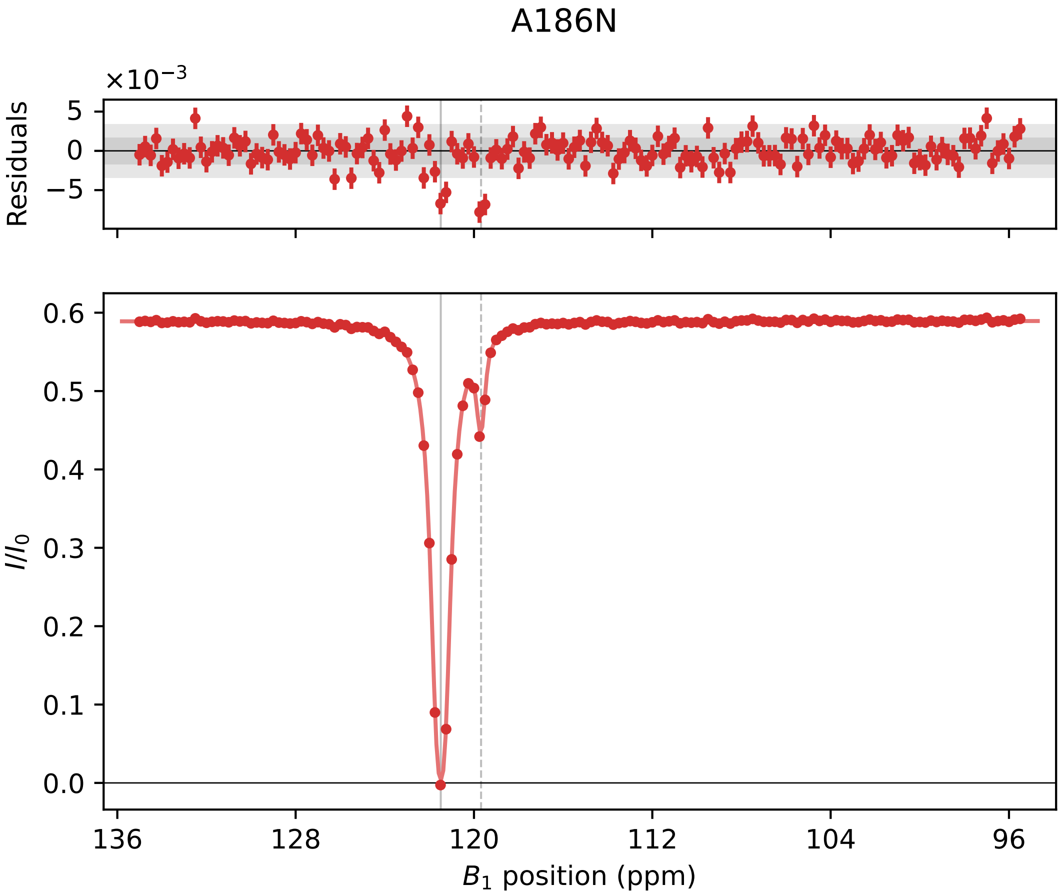

A186N
×10⁻³
Residuals
5
0
−5
I/I₀
0.6
0.5
0.4
0.3
0.2
0.1
0.0
136
128
120
112
104
96
B₁ position (ppm)

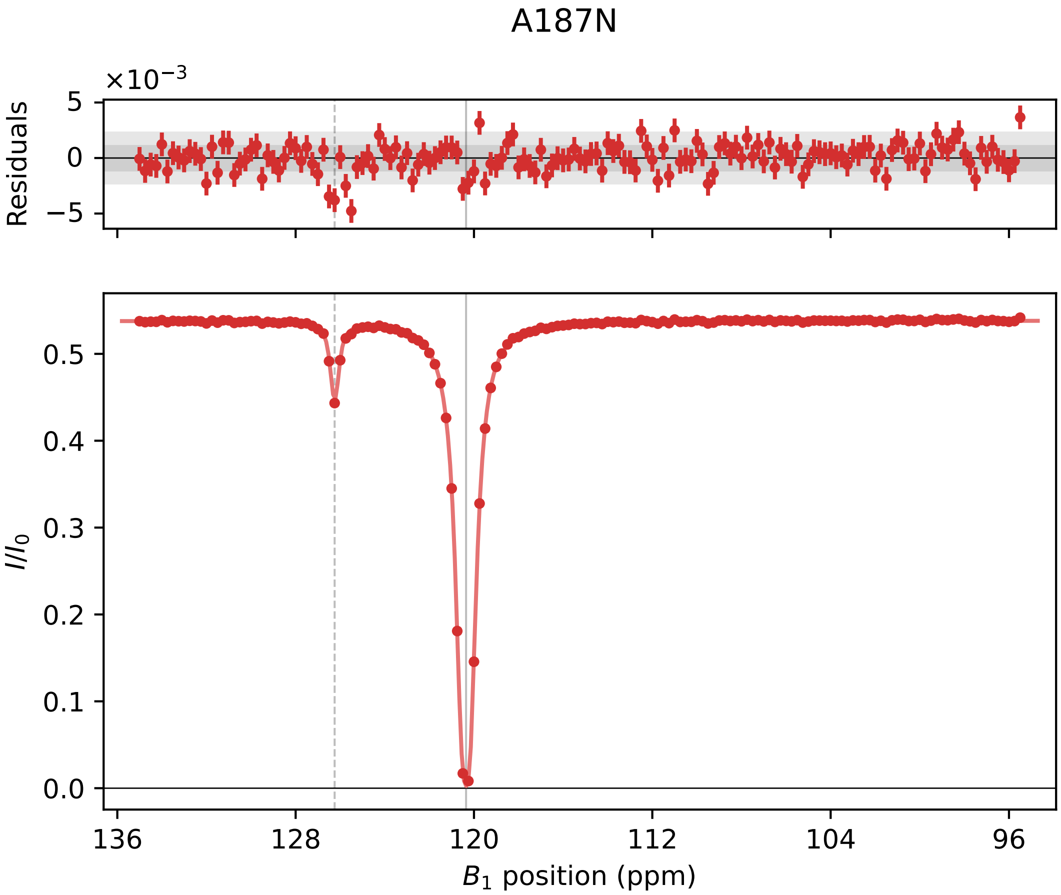

A187N
×10^−3
Residuals
5
0
−5
I/I0
0.5
0.4
0.3
0.2
0.1
0.0
136
128
120
112
104
96
B1 position (ppm)

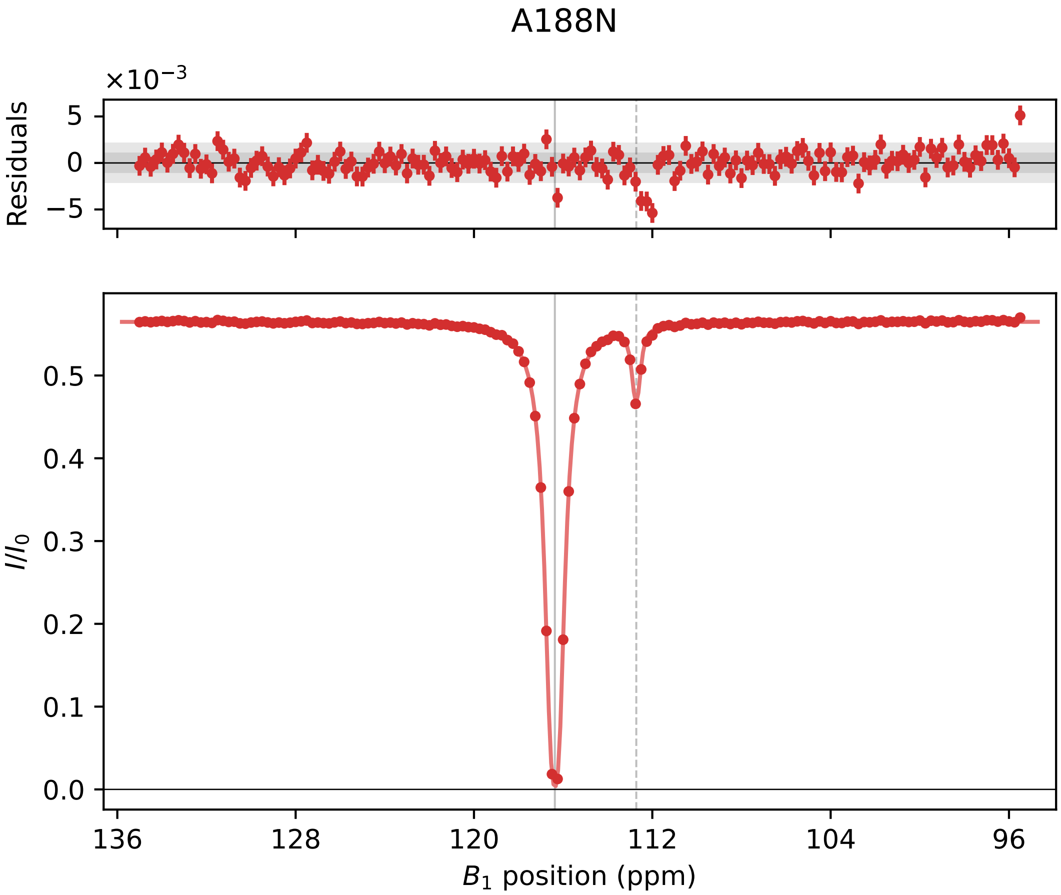

A188N
×10⁻³
Residuals
5
0
−5
I/I0
0.5
0.4
0.3
0.2
0.1
0.0
136
128
120
112
104
96
B1 position (ppm)

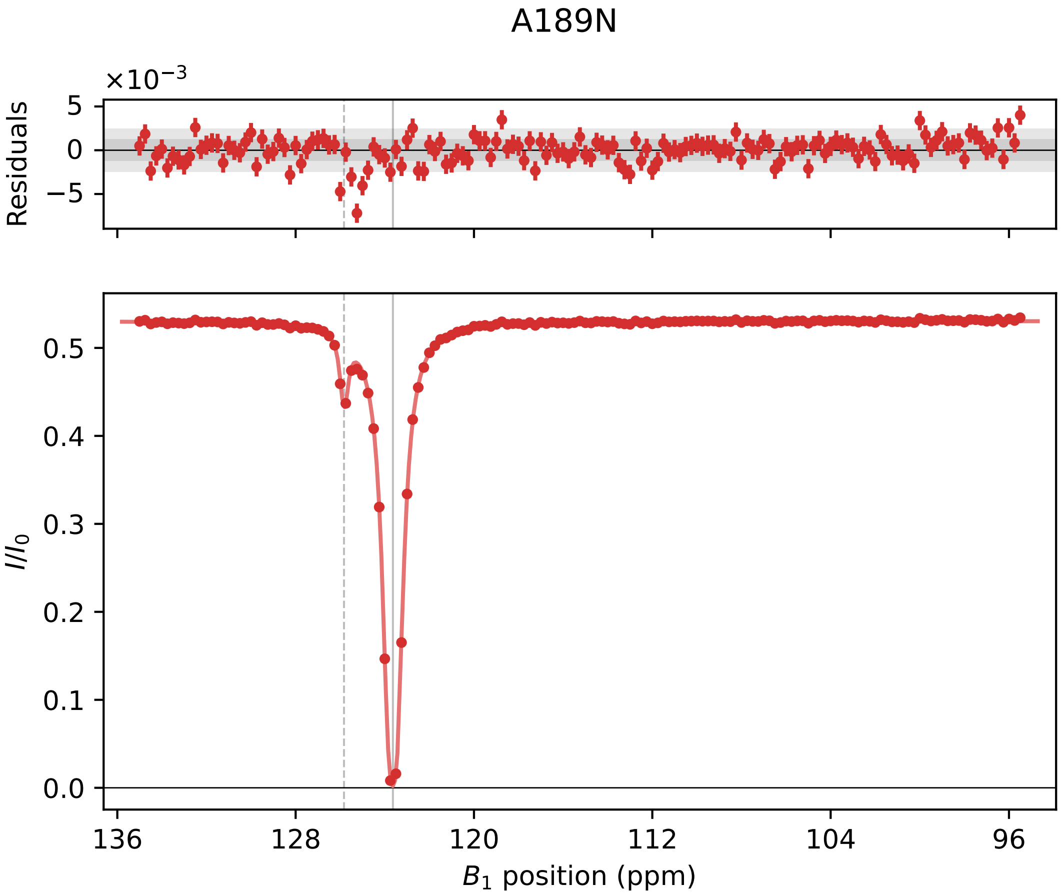

A189N
×10⁻³
Residuals
5
0
−5
I/I0
0.5
0.4
0.3
0.2
0.1
0.0
136
128
120
112
104
96
B1 position (ppm)

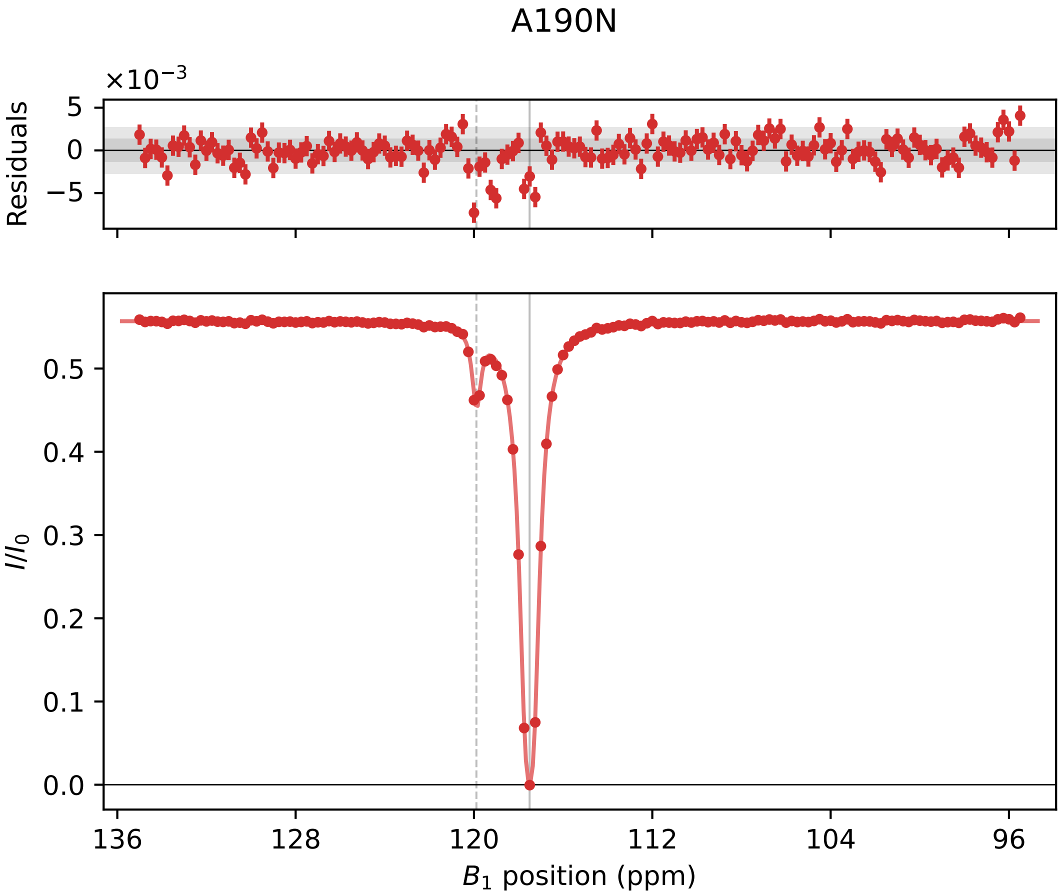

A190N
×10−3
Residuals
5
0
−5
0.5
0.4
0.3
0.2
0.1
0.0
I/I0
136
128
120
112
104
96
B1 position (ppm)

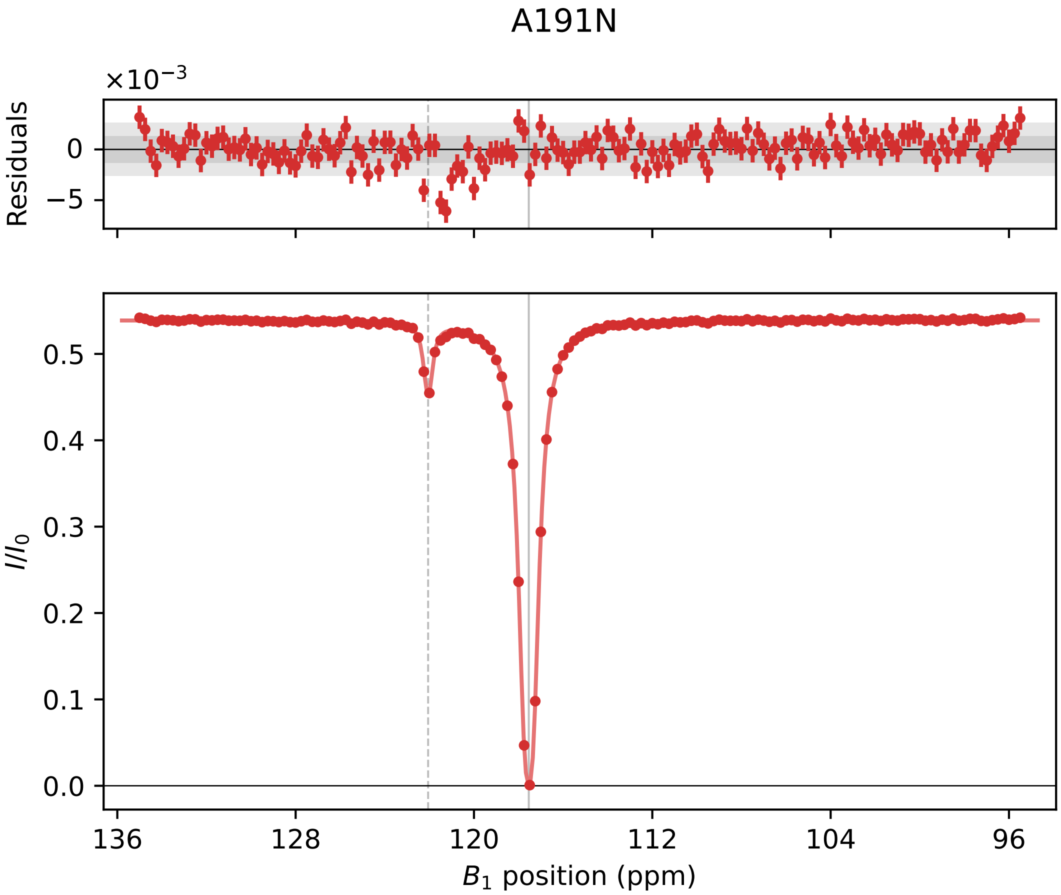

A191N
×10⁻³
Residuals
0
−5
I/I0
0.5
0.4
0.3
0.2
0.1
0.0
136
128
120
112
104
96
B1 position (ppm)

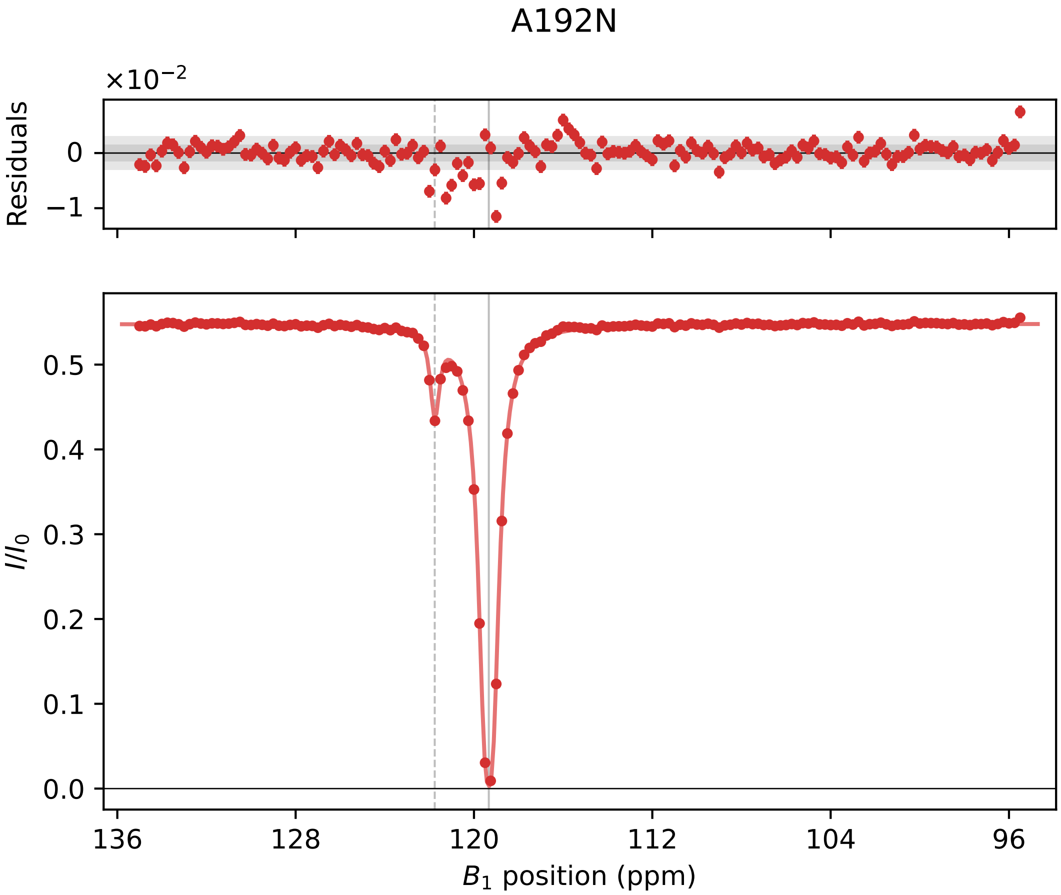

A192N
×10−2
Residuals
0
−1
I/I0
0.5
0.4
0.3
0.2
0.1
0.0
136
128
120
112
104
96
B1 position (ppm)

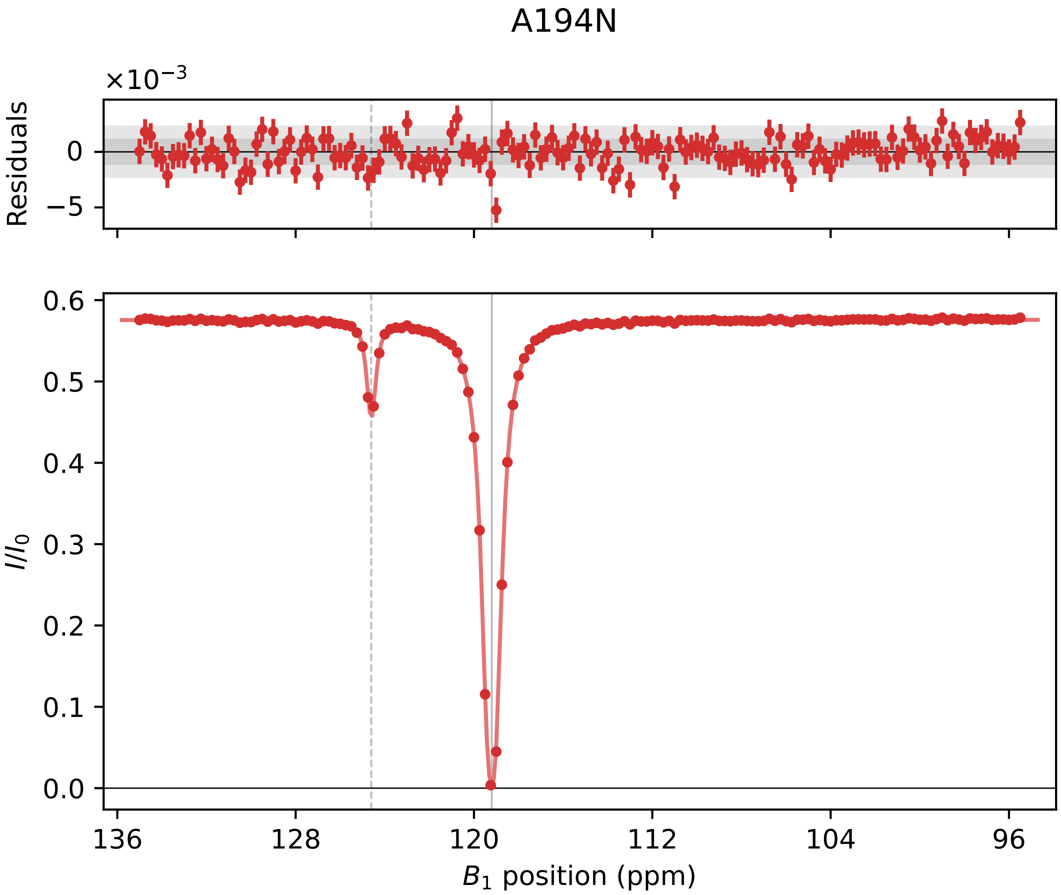

A194N
×10^−3
Residuals
0
−5
I/I0
0.6
0.5
0.4
0.3
0.2
0.1
0.0
136
128
120
112
104
96
B1 position (ppm)

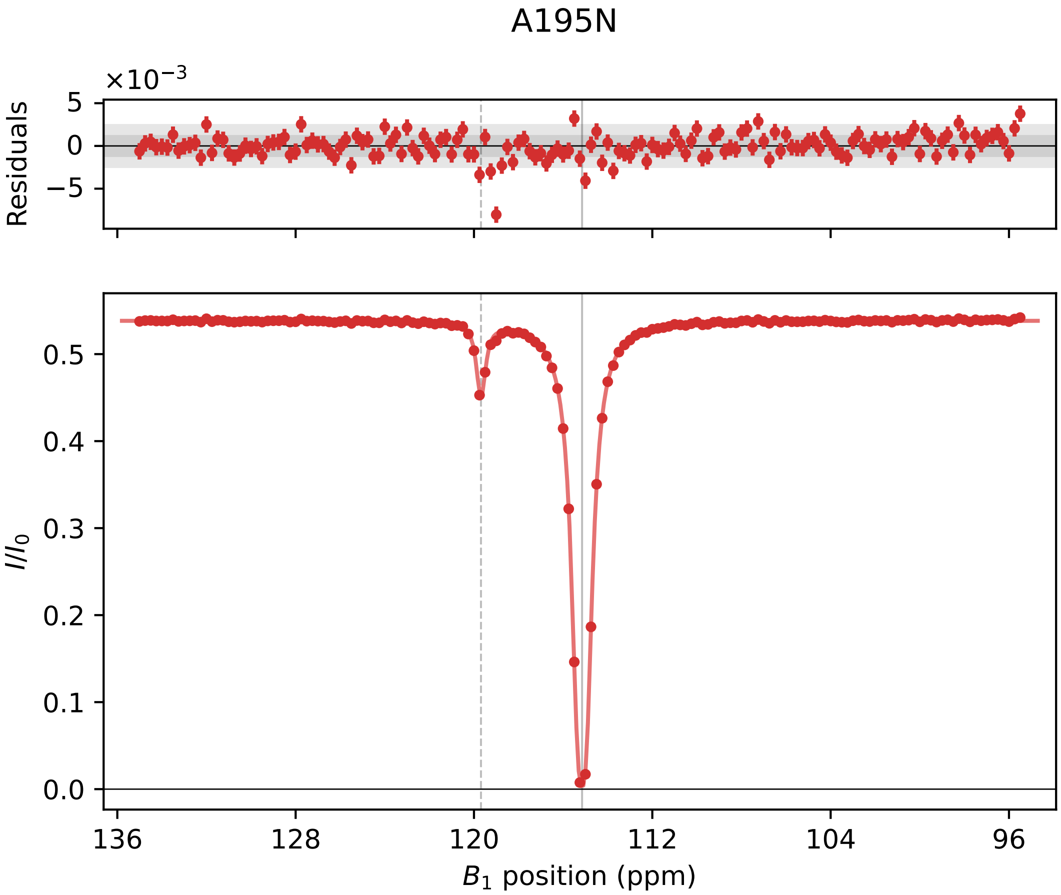

A195N
×10⁻³
Residuals
5
0
−5
0.5
0.4
0.3
0.2
0.1
0.0
I/I0
136
128
120
112
104
96
B1 position (ppm)

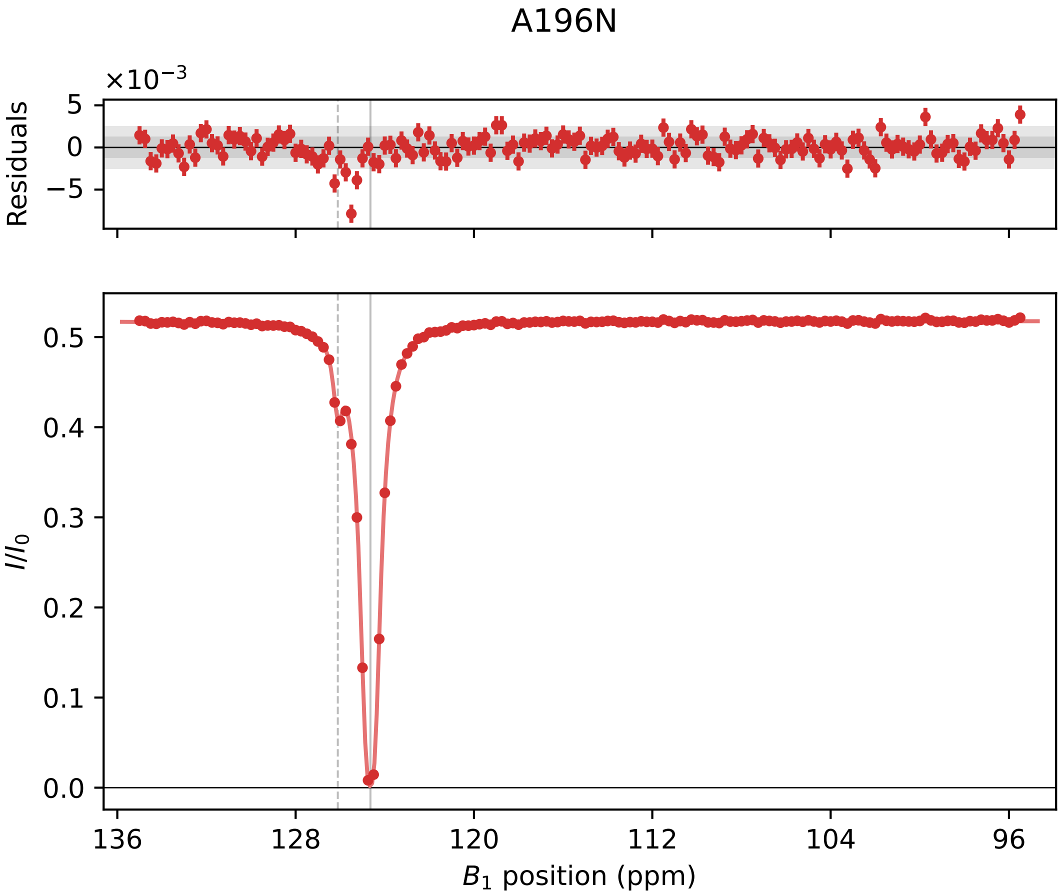
A196N
×10⁻³
Residuals
5
0
−5
0.5
0.4
0.3
0.2
0.1
0.0
I/I0
136
128
120
112
104
96
B1 position (ppm)

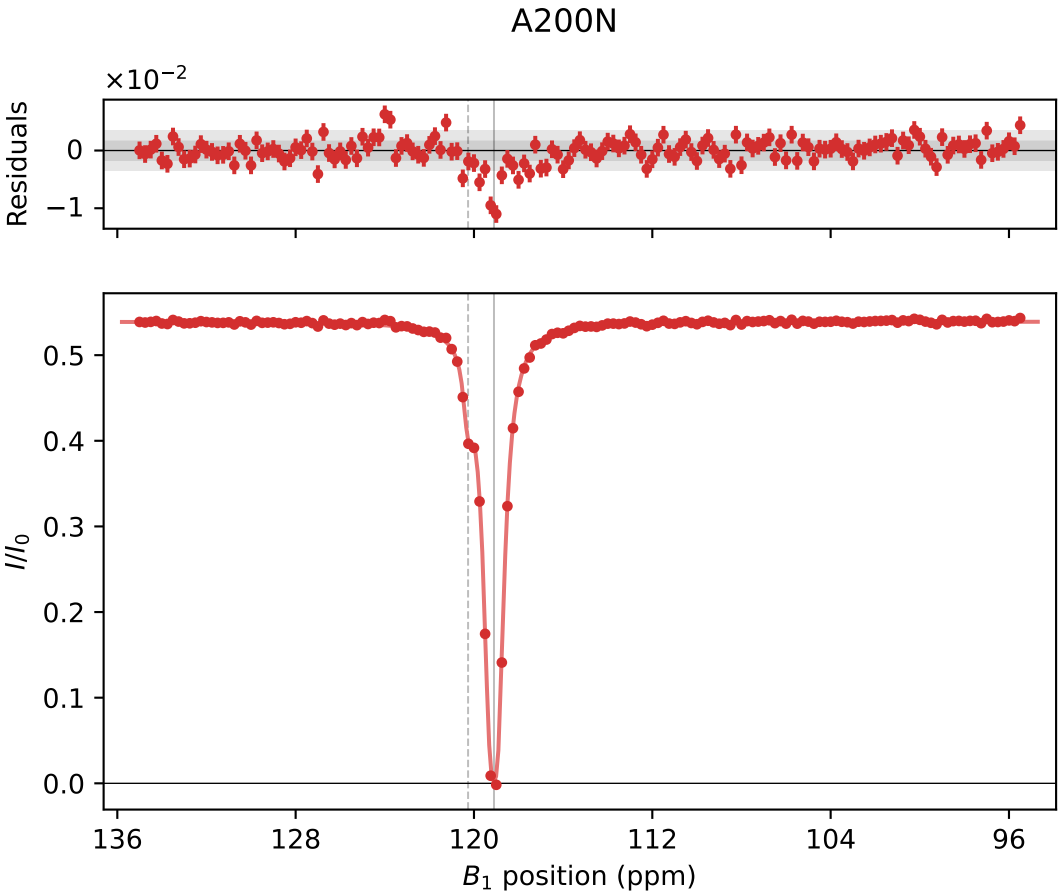
A200N
$\times10^{-2}$
Residuals
0
−1
$I/I_0$
0.5
0.4
0.3
0.2
0.1
0.0
136
128
120
112
104
96
$B_1$ position (ppm)

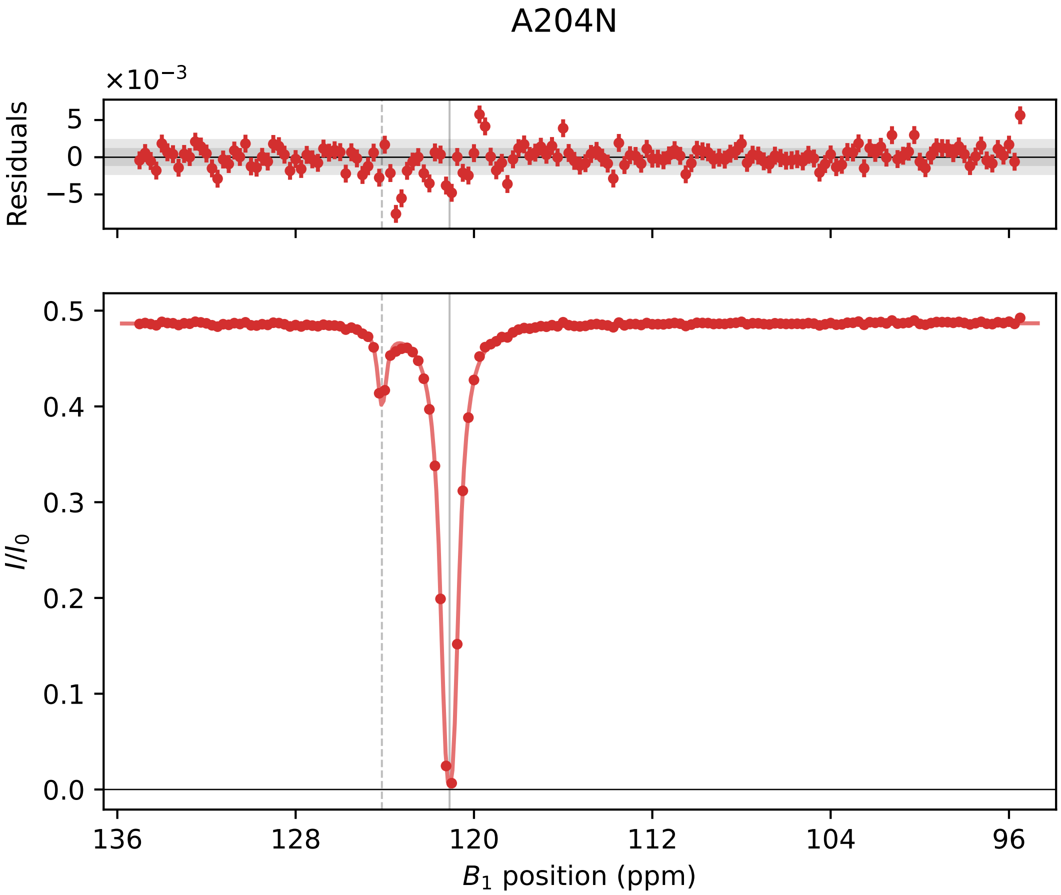

A204N
×10⁻³
Residuals
5
0
−5
0.5
0.4
0.3
0.2
0.1
0.0
I/I0
136
128
120
112
104
96
B1 position (ppm)

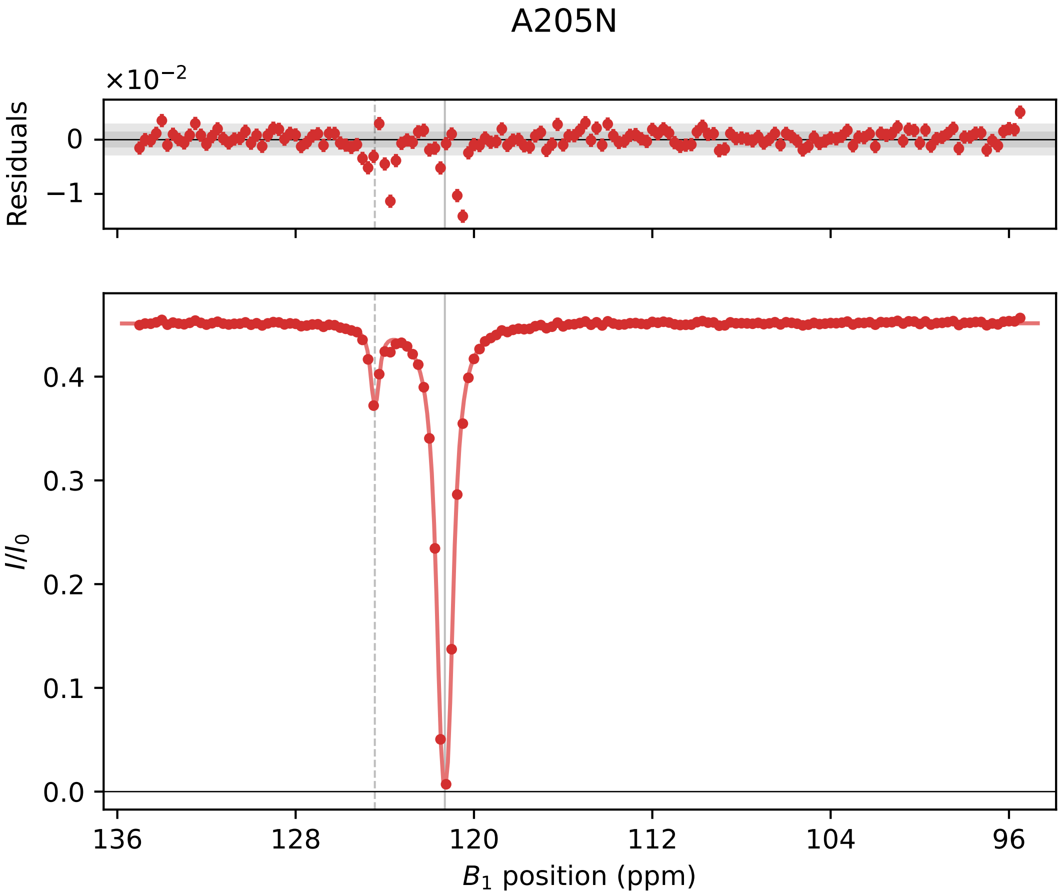

A205N
×10⁻²
Residuals
0
−1
I/I0
0.4
0.3
0.2
0.1
0.0
136
128
120
112
104
96
B1 position (ppm)

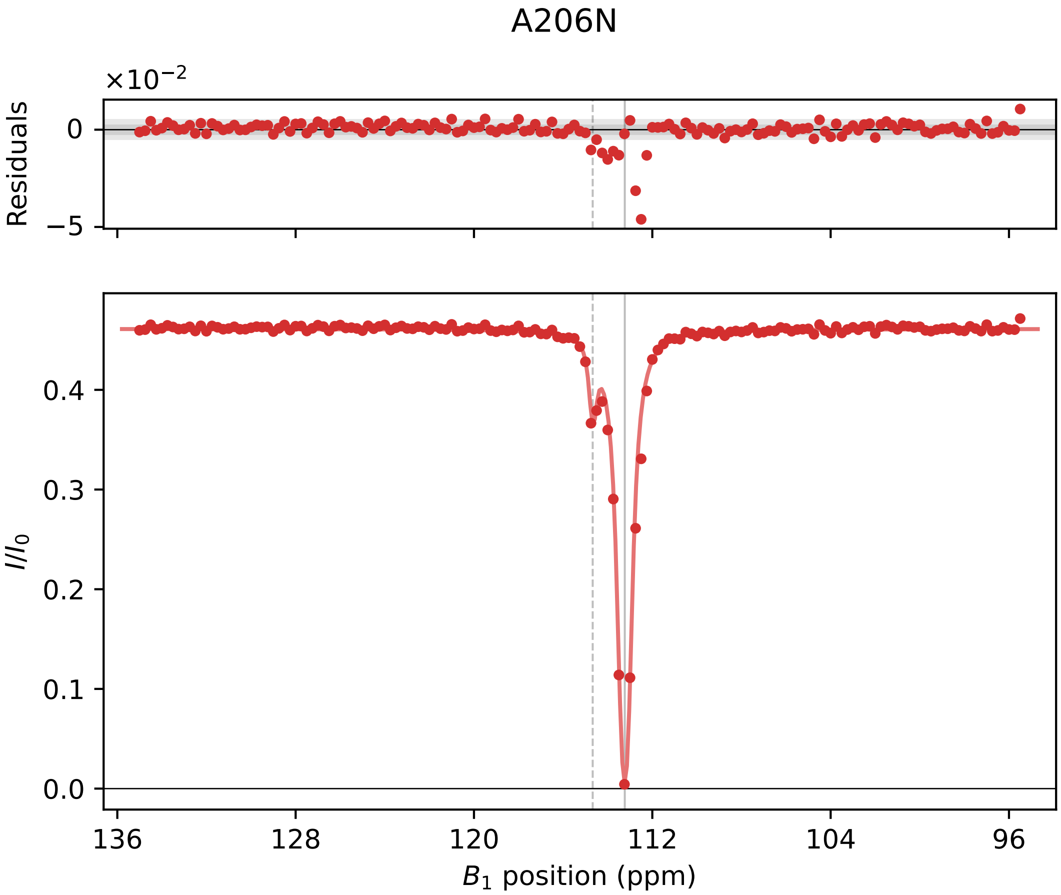

A206N
×10⁻²
Residuals
0
−5
I/I0
0.4
0.3
0.2
0.1
0.0
136
128
120
112
104
96
B1 position (ppm)
